# Primate sympatry shapes the evolution of their brain architecture

**DOI:** 10.1101/2022.05.09.490912

**Authors:** Benjamin Robira, Benoît Perez-Lamarque

**Author notes:** Correspondence: Benjamin Robira < >.

## Abstract

The main hypotheses on the evolution of animal cognition emphasise the role of conspecifics in affecting the socio-ecological environment shaping cognition. Yet, space is often simultaneously occupied by multiple species from the same ecological guild. These sympatric species can compete for food, which may thereby stimulate or hamper cognition. Considering brain size as a proxy for cognition, we tested whether species sympatry impacted the evolution of cognition in frugivorous primates. We first retraced the evolutionary history of sympatry between frugivorous primate lineages. We then fitted phylogenetic models of the evolution of the size of several brain regions in frugivorous primates, considering or not species sympatry. We found that the evolution of the whole brain or brain regions used in immediate information processing was best fitted with models not considering sympatry. By contrast, models considering species sympatry best predicted the evolution of brain regions related to long-term memory of interactions with the socio-ecological environment, with a decrease in their size the higher the sympatry. We speculate that species sympatry, by generating intense food depletion, might lead to an over-complexification of resource spatiotemporality that counteracts the benefits of high cognitive abilities and/or might drive niche partitioning and specialisation, thereby inducing lower brain region sizes. In addition, we reported that primate species in sympatry diversify more slowly. This comparative study suggests that species sympatry significantly contributes to shaping primate evolution.

## INTRODUCTION

Cognition evolution is shaped by the balance between socio-ecological drivers promoting cognitive abilities (González-Forero and Gardner 2018) and physiological and energetic constraints limiting them (Navarrete, van Schaik, and Isler 2011). Primates are pivotal species for cognitive studies (Byrne 2000) because their cognition (i.e. the set of mechanisms that enable them to perceive, learn and memorise information, and make decisions, Shettleworth 2010) is thought to be promoted by interactions of individuals with conspecifics within the social unit (Byrne 2018; Dunbar and Shultz 2017), among generations (Wilson 1991; Whiten and van Schaik 2007; Reader and Laland 2002; Herrmann et al. 2007; Tomasello 2019; van Schaik and Burkart 2011), between social units (Ashton, Kennedy, and Radford 2020), or with the rest of their environment (Clutton Brock and Harvey 1980; Milton 1981; Rosati 2017). However, space is often occupied by many primate species, which may have dietary overlaps and compete for the same resources (Kamilar and Ledogar 2011). Because of direct and indirect interactions linked to resource availability, the presence of heterospecifics is also likely to shape the evolution of species cognition.

Retracing the evolutionary history of cognitive abilities proves to be challenging because there is still no consensual measurement for cognition applicable across all species. Up to now, a raw approximation consists in considering the (relative) brain size as a proxy for cognitive abilities, with larger sizes considered equivalent to more advanced cognitive abilities (e.g. in mammalian carnivores: Benson-Amram et al. 2016; or primates: MacLean et al. 2014). Although the relevance of this assumption is heavily limited within species, in part because of plasticity (Gonda, Herczeg, and Merilä 2013), this holds when comparing different species within taxonomically similar groups (e.g. in primates, Reader and Laland 2002). In addition, instead of considering the brain as a whole, the multifaceted aspect of animal cognition is more precisely depicted by appreciating the mosaic nature of the brain (Barton and Harvey 2000). For instance, variations in the size of some specific brain regions have been robustly associated with variations in cognition related to the functions of these regions (Healy and Rowe 2007). The brain is a patchwork of areas cognitively specialised that may thus follow different evolutionary trajectories.

Because the coexistence of primate species in the same biogeographic area, henceforth referred to as sympatry, can affect multiple aspects of their social-ecological environment, brain regions may be differently affected by sympatry. Considering a simplistic functional picture of the primate brain, several hypotheses can be proposed on how species sympatry might, through direct or indirect competition or cooperation, influence the relative benefits of cognition (i.e. the balance between its benefits and its costs). As such, selective pressures upon cognitive abilities would affect the energy allocation toward the whole brain and/or the within-brain allocation toward the different brain regions, resulting in changes of their sizes relative to the whole body and/or the whole brain, respectively.

First, long-term memory stands as a valuable tool to infer food availability and location when food is rare and ephemeral but predictable (Milton 1981; Rosati 2017). Having several sympatric species competing for the same food resource often leads to an increase in food depletion of the shared resource compared with an environment with only one foraging species (Minot 1981). Such depletion may therefore complexify the patterns of resource distribution and availability in space and time. This complexification may in turn affect the selective pressures upon brain regions involved in the long-term storing of spatiotemporal information, such as the hippocampus (Burgess, Maguire, and O’Keefe 2002, Hypothesis 1: long-term memory is affected by sympatry). On the one hand, under reasonable food depletion, having better memory should be advantageous to better predict food availability and location. In addition, resource competition between heterospecifics should promote anticipatory behaviour, hence high cognition, as expected for within-species competition (Ashton, Kennedy, and Radford 2020). Thus, the relative size of the hippocampus, reflecting long-term memory abilities, should be larger the higher the sympatry (Prediction 1.1). On the other hand, intense food depletion may also increase environmental unpredictability to a point where long-term memory is not advantageous anymore. In addition, even when the dietary overlap between heterospecifics is inexistent, foragers may rely on phenological cross-correlations between plant species (e.g. if plant species A produces fruits earlier in the season than plant species B, thus the availability of B fruits can be predicted based on the availability of A fruits, even though A is not consumed by the forager, Janmaat et al. 2012; Robira et al. 2021): Intense food depletion may thus weaken these cross-correlations and make food availability predictions less reliable, therefore limiting the benefits of memory (Robira et al. 2021). Due to the energy constraints of maintaining a large brain, the hippocampus relative size could thus be smaller in species experiencing high levels of sympatry (Prediction 1.2). Furthermore, the competition between sympatric species from a given dietary guild may foster their specialisation toward different food resources, i.e. niche partitioning, which may impact cognition (Aristide et al. 2016). While a species might specialise in food that is difficult to access and requires high cognition (e.g. through the use of tools), specialisation is generally associated with reduced flexibility and thus lower cognitive abilities (Henke-von der Malsburg, Kappeler, and Fichtel 2020). Therefore, because of cognitive specialisation requires less long-term memory abilities, the hippocampus relative size could also be smaller in sympatric species (Prediction 1.2). Alternatively, intense levels of sympatry could also lead to a disruptive selection toward cognitive abilities, with the most successful competitors that may evolve enhanced cognition, while bad competitors may not. In this context, the hippocampus relative sizes may either increase or decrease for the different sympatric species (Prediction 1.3; character displacement).

Second, sympatric species may enrich the landscape with visual, olfactory, or acoustic cues usable to locate available food (e.g. Avarguès-Weber, Dawson, and Chittka 2013; Kashetsky, Avgar, and Dukas 2021). Consequently, such cues left out by heterospecifics may impact the selective pressures upon brain regions involved in processing more immediate sensory information, such as the Main Olfactory Bulb (MOB), the cerebellum (Koziol et al. 2014; Sokolov, Miall, and Ivry 2017), and the neocortex (Wiltgen et al. 2004) (Hypothesis 2: cue processing is affected by sympatry). Hence, sympatry could be associated with larger relative sizes of the MOB, the cerebellum, or the neocortex (Prediction 2). As brain regions may be interconnected, this enlargement could be detrimental to other brain regions. For instance, increased corticalisation could be associated with smaller hippocampus sizes (Kitamura et al., 2017).

Third, besides indirect interactions through foraging, cognition can also be triggered by direct “social” interactions with other individuals (Byrne 2018; Dunbar and Shultz 2017), such as with the formation of mixed-group species (Goodale et al. 2010). The striatum, a brain region stimulated during social interactions (Báez-Mendoza and Schultz 2013), or to a lesser extent, the hippocampus (Todorov et al. 2019), may therefore be affected by the increase in proximity, hence direct social interactions, between heterospecifics (Hypothesis 3: sociality is affected by sympatry), leading to a larger relative size of the striatum or hippocampus in sympatry (Prediction 3).

In any case, the evolution of brain size in primates likely impacted their dynamic of species diversification. Larger brain sizes are indeed found to be associated with higher diversification rates in birds (Sayol et al. 2019) and similar patterns have been suggested in primates (Melchionna et al. 2020). However, it remains unclear how brain size and diversification are interlinked in the context of sympatry.

Here, we investigated whether species sympatry affected the evolution of cognition using frugivorous primates as a study example. Frugivorous primates are an interesting group for such a question because fruit is the archetype of a hard-to-find resource yet predictable (Janmaat et al. 2016), for which cognition considerably shapes the foraging strategy (Trapanese et al. 2019). To infer the effect of species sympatry on cognition in frugivorous primates, we evaluated the support for phylogenetic models of brain size evolution accounting or not for species sympatry and investigated the directionality of the selection induced by sympatry on brain size evolution. Finally, we tested for correlative patterns between brain size or current sympatry and species diversification in all primates, to better understand the impact of cognition and interactions between primates on their evolutionary success.

## METHODS

Data processing, analyses, and plots were computed with R software (v.4.1.2, R Core Team 2020). The list and versions of packages are available in Supplementary Material, R packages used.

### Data Collection

#### Trait data

We pooled data from previous literature surveys (Pearce et al. 2013; Willems, Hellriegel, and van Schaik 2013; Grueter 2015; DeCasien, Williams, and Higham 2017; Powell, Isler, and Barton 2017; Navarrete et al. 2018; DeCasien and Higham, 2019; Powell, Barton, and Street 2019; Todorov et al. 2019, see Supplementary Material, Trait data collection). From the global endocranial brain volume, we obtained the Encephalization Quotient (EQ, _N*EQ, max*_ = 182) such as EQ = 1.036 x Brain size/0.085 × Body mass^0.775^ with the brain size in cm^3^, 1.036 g/cm^3^ being the assumed homogeneous brain density, and the body mass in g. EQ indicates whether the brain size ranges above (> 1) or below (< 1) what is expected given the body mass and correcting for allometry (DeCasien, Williams, and Higham 2017). We also focused on specific regions of the brain which were chosen because they were involved in immediate sensory information processing (MOB, N_*MOB,max*_ = 39), in movement and/or general information processing and retention (neocortex, N *_Neocortex,max_* = 69, Wiltgen et al. 2004; cerebellum, N *_Cerebellum,max_* = 70, Koziol et al. 2014; Sokolov, Miall, and Ivry 2017), or short-term working memory and long-term spatiotemporal memory (hippocampus, N *_Hippocampus,max_* = 63, Burgess, Maguire, and O’Keefe 2002). The striatum (N *_Striatum,max_* = 63) supports information processing during social interaction, reward assessment, planning, or goal-oriented behaviours (Báez-Mendoza and Schultz 2013; Johnson, Meer, and Redish 2007). To investigate their evolutionary history, we first used the ratio between their volume and body mass. As such, the use of brain region sizes relative to the body mass and not raw sizes depicts the evolution of cognitive abilities in terms of differential allocation rather than abilities per se (but see discussion in Deaner, Nunn, and van Schaik 2000). Second, we repeated the analyses considering the ratio between the brain region size and the whole-brain size, as this might reflect within brain energy reallocation.

For each primate species, broad dietary guilds (frugivores, folivores, nectarivores, granivores, insectivores, omnivores) are generally considered based on the main type of food they consume. We classified extant species as either “frugivorous” or “folivorous” based on the availability of percentage of frugivory and folivory, prioritising frugivory over folivory. We considered a species as frugivorous if the percentage of frugivory was at least above 20%. If not, or if the percentage of frugivory was unavailable, we considered the species as folivorous if the percentage of folivory was above 40%. Whenever the percentages were not available, we referred to the classification provided by DeCasien, Williams, and Higham (2017), partly based on anatomical criteria. We discarded any other cases (e.g. species feeding on invertebrates or tree exudates). We also replicated these diet assignments by considering a threshold of 40% for frugivory and 60% for folivory.

#### Ranging Data

The current biogeographic range of each primate species was assessed using ranging maps provided by the IUCN red list (IUCN 2021). Ranging data were available for 249 species. The availability of distribution range data is depicted in Supplementary Figure S1.

#### Phylogeny

We used chronogram trees of the primate taxon of the 10kTrees project (downloaded on May 2021, version 3), as well as a consensus tree of 1000 trees for the subsequent phylogenetic analyses. The trees contain 301 primate species. Note that in all these analyses, we discarded *Homo sapiens* and *Macaca sylvanus*. The latter was discarded because of its complete geographic isolation and repeated intervention of humans in population maintenance (Modolo, Salzburger, and Martin 2005). A summary of available data per species is presented in Supplementary Figure S1.

### Retracing past sympatry between primate species

Based on the biogeographic distribution of each extant primate species, we first reconstructed the history of past sympatry between primate lineages. To do so, we followed Drury et al. (2018) and first reconstructed the biogeographic history of each primate lineage to then retrace which pairs of primate lineages were likely to be simultaneously present at the same place. Leaning on Kamilar (2009), we considered that the biogeography of primates can be described by 12 discrete biogeographic areas with highly similar community structures shaped by both the environment geography and climatic correlates. These geographic areas, mapped using Google Earth Professional (v7.3.3), are represented in Figure 1. One to multiple biogeographic areas were assigned to each species as soon as 10% of their current distribution range overlapped on the surface with a given biogeographic area (see Supplementary Material, Spatial overlap assessment, Figure 2).

**Figure 1:**
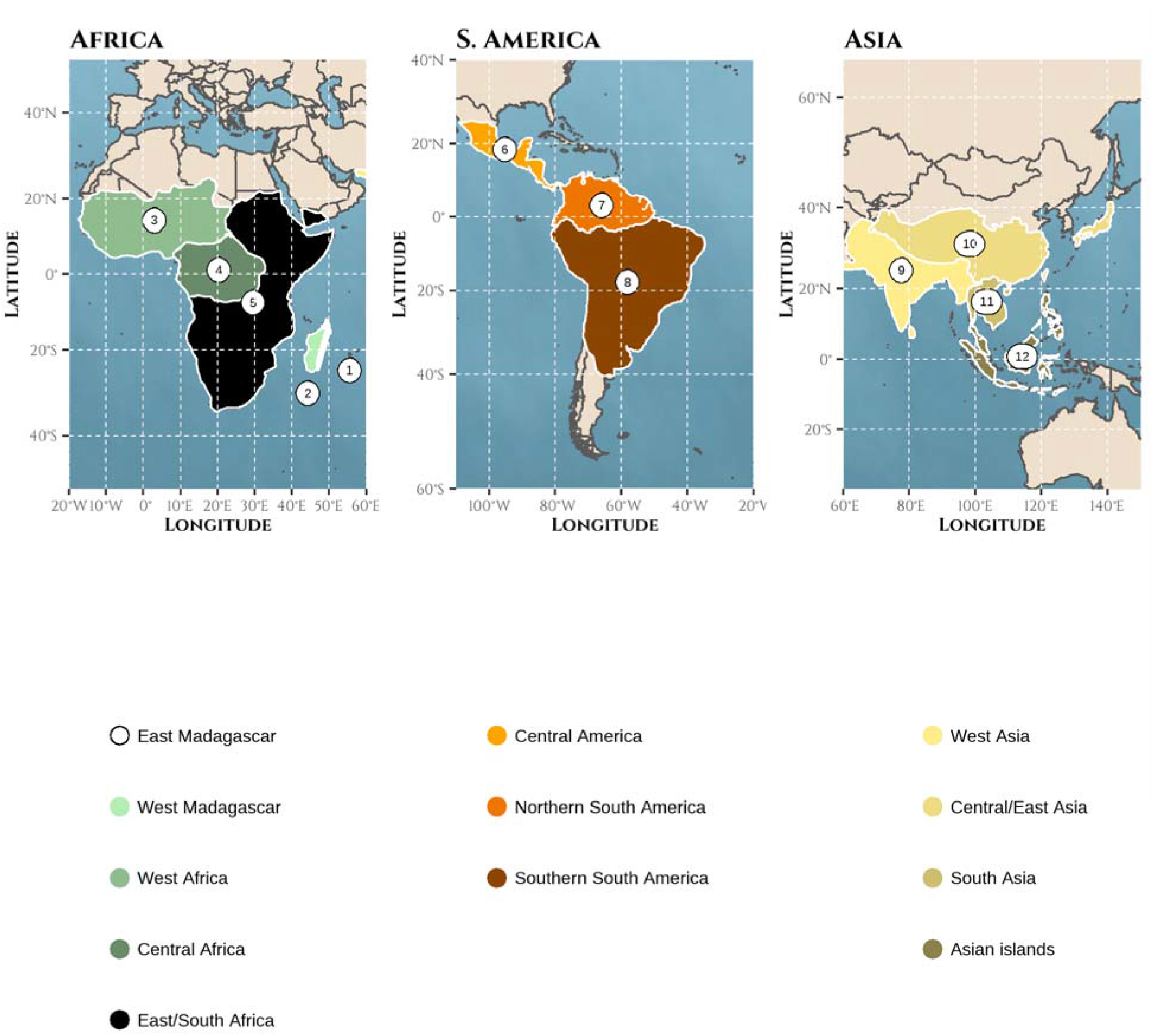
Biogeographic areas used for reconstructing the history of sympatry in frugivorous primates represented on the Mercator projection of the world | Areas were defined as a combination of geographic and environmental criteria relative to the primate taxonomy following results from Kamilar (2009): (1) East Madagascar (2) West Madagascar (3) West Africa (4) Central Africa (5) East/South Africa (6) Central America (7) North South-America (8) South South-America (9) West Asia (10) Central/East Asia (11) South Asia (12) Asian peninsula and islands. Note that northern Africa and southern Europe were discarded because *Macaca sylvanus* was not considered.

**Figure 2:**
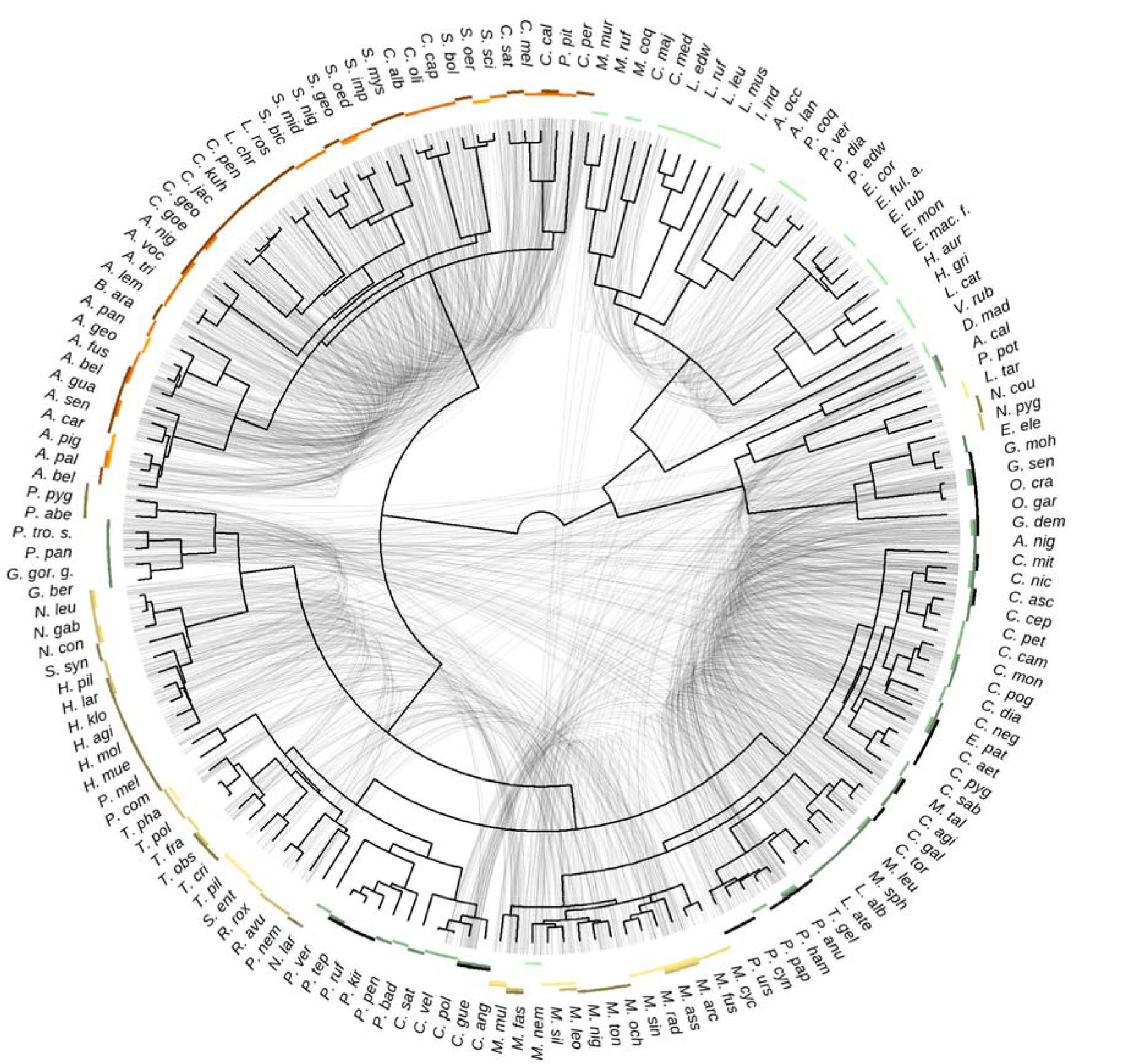
The level of species sympatry varies across the primate phylogeny | Primate phylogeny from the consensus tree of the 10kTrees project is depicted in the center, together with abbreviated species names. The corresponding non-abbreviated names can be found using Supplementary Figure S1. Sympatric frugivorous (based on a frugivory threshold of 20% and folivory threshold of 40%) species are linked by light grey lines. The biogeographic areas occupied by a species are depicted by coloured rectangles. Presence was assessed given an overlap between the species range and the biogeographic area of 10%.

Given these 12 biogeographic areas, we retraced the biogeographic history of primates with the *BioGeoBEARS* package (Matzke 2013), using the biogeographic stochastic mapping algorithm (Matzke 2016). We fitted non-time-stratified dispersal-extinction-cladogenesis (DEC) models. We fixed to three biogeographic areas the maximum number of areas that a lineage can simultaneously occupy since it offers the possibility to occupy a complete mainland continent while keeping computational time reasonable. We sampled 10 histories of primate biogeographic ranges to account for the uncertainty. We assumed that two primate lineages were in sympatry at a given time whenever these two lineages simultaneously occupied the same biogeographic area. We also replicated these biogeographic assignments using instead a larger threshold of 30% of overlap (instead of 10%; see Supplementary Figure S2 for a quantification of variations in biogeographic assignments due to the use of different thresholds).

### Inferring past diets of primate lineages

Next, we retraced the evolutionary history of frugivorous lineages in primates. Considering diet as a binary variable, we retraced the evolutionary history of frugivory versus folivory using extended Mk models (Bollback 2006) implemented in the “simmap” function of the *phytools* package (Revell 2012). We performed ancestral diet reconstructions using both combinations of dietary thresholds (20/40% and 40/60% for frugivory and folivory, respectively) and sampled 10 stochastic mapping of ancestral diet reconstructions per run. These ancestral reconstructions were used in combination with the histories of primate biogeographic ranges to assess whether a pair of frugivorous species was in sympatry or not (Drury et al. 2018): we thus obtained reconstructions of the evolutionary history of sympatry between frugivorous primate lineages.

### Phylogenetic models of trait evolution: does species sympatry shape brain size evolution?

We assessed the effect of sympatry on the evolution of frugivorous primate brain size using two approaches. First, we used phylogenetic models of trait evolution to assess the role of sympatry in the evolution of brain sizes. Second, we investigated how sympatry has influenced brain size evolution (i.e. selection towards smaller or larger brain sizes) by evaluating correlations between current levels of sympatry and brain sizes using linear models. In both approaches, we considered either the whole brain (using the EQ) or the size of the aforementioned specific brain regions relative (i) to the body mass or (ii) to the whole-brain size. Models testing the effect of species sympatry on one brain region relative to the body mass (models (i)) assess whether the total allocation towards a specific brain region is affected by species sympatry. In contrast, models testing the effect of species sympatry on one brain region relative to the whole-brain size (models (ii)) assess whether the within-brain energy reallocation is affected by species sympatry. Before fitting, relative brain sizes were log-transformed to account for allometry. Although it does not consider the potential inter-dependence between brain regions, we modelled independently each brain region to preserve the maximum power (i.e. the largest sample size) in each analysis.

#### (a)#Fitting models of trait evolution

For each brain region, we independently fitted different phylogenetic models of trait evolution. Following Drury et al. (2016), we considered three stochastic models accounting for sympatry that expand Brownian motion (BM) models of trait evolution. We used the “fit_t_comp” function from the *RPANDA* package (Morlon et al. 2016) to fit: a matching competition (MC) model (Nuismer and Harmon 2015) and linear and exponential density-dependent models (DD*_lin_* and DD*_exp_*, Drury et al. 2016). The MC model considers the repulsion of brain sizes of sympatric frugivorous lineages due to competition (character displacement), that is 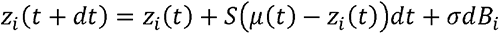 where *z* is the brain size of a species *i* at time *t*, *μ* is the mean brain size of all sympatric species, *S* is a negative value that reflects the strength of the effect of species sympatry, and *σdB_i_* is the drift (BM) with a constant evolutionary rate *σ* (Drury et al. 2016). Linear (DD*_lin_*) or exponential (DD*_exp_*) density-dependences (Drury et al. 2016; Weir and Mursleen 2013) assume that the evolutionary rate, *σ_i_*, of trait evolution, varies either positively or negatively as a function *f* of the number of frugivorous sympatric lineages, such as *σ_i_* = *f_in_(l)* = **σ*_0_* + *rl* and *σ_l_* = *f_exp_(l)* = **σ*_0_*exp(*rl*), where *σ*_0_ corresponds to the ancestral rate value, *l* indicates the number of lineages at a given time, and *r* controls the speed and direction of the dependency of the rate to the number of lineages (*r* > 0 leads to an increased rate of trait evolution in sympatry, while r < 0 leads to a slowdown of trait evolution in sympatry, as observed in adaptive radiations).

We compared the support of models considering species sympatry to the support of simpler models assuming no effect of species sympatry on the evolution of brain sizes: the Brownian motion (BM), the Ornstein-Uhlenbeck process (OU, a model with an optimum value, see Blomberg, Rathnayake, and Moreau 2020, for a review), or the Early-Burst model (EB), for assessing a time-dependence of the evolutionary rate, irrespectively of the levels of species sympatry (Blomberg, Garland, and Ives 2003). We fitted these models using the “fitContinuous” function from the *geiger* package (Slater et al. 2012; Pennell et al. 2014). We compared the six models accounting for sympatry or not within an information-theoretic framework (Burnham and Anderson 2002), based on the weights of Akaike Information Criteria corrected for small samples (AICc). The model weight depicts how well the model fits the observed data compared with the other tested models (i.e. a model set).

To account for uncertainty (in the data, Supplementary Figure S3, and fit), we fitted and compared these models (within each model set) 10 times for 10 different ancestral reconstructions of primate biogeography and diet considering all thresholds. Thus, for each brain region, we fitted all the models 10 (uncertainty on diet/biogeography ancestral reconstructions x 10 (uncertainty in brain/diet data) x 2 (biogeographic overlap threshold) x 2 (frugivory threshold) x 2 (folivory threshold) = 800 times. We nonetheless stopped computations when the calculation of the likelihood was excessively long (> 1 week) and the final sample size was 730 model sets.

#### (b)#Determining the effect of sympatry on brain sizes

Diversity-dependent models of trait evolution inform whether species sympatry has impacted brain size evolution by increasing or decreasing the tempo of trait evolution. Nonetheless, they do not tell the directionality of the effect (i.e. do brain sizes increase or decrease with sympatry?). To determine whether species sympatry positively or negatively affected the sizes of brain regions, we independently fitted Gaussian Pagel’s lambda phylogenetic regressions for each brain region of extant frugivorous species. This model is a derivative of the BM model, where the phylogenetic variance-covariance matrix has all coefficients, but its diagonal ones, multiplied by lambda (therefore relaxing the hypothesis of BM). To fit these models, we used a frequentist-based approach with the “phylolm” function from the *phylolm* package (Ho and Ane 2014). We considered the least stringent frugivory assessment, with the frugivory threshold fixed at 20% and the folivory threshold fixed at 40%.

The response variable was the log-transformed relative size of each brain region relative to the body mass or the whole-brain size. Doing so controlled for allometry while being in line with the phylogenetic models of trait evolution, which cannot account for additional variables and are thus limited to the use of relative brain sizes. Due to data variability, we took the mean of the possible values given the different datasets and assessed the sensitivity using non-averaged values (see Supplementary Material, Model stability). We used as covariates (i.e. continuous predictors) two explicit measures of the level of species sympatry for each extant frugivorous species: (1) the number of frugivorous sympatric species (square-rooted to reach symmetrical distribution to limit leverage effects) and (2) the average percentage of the overlapping current range (assessed based on IUCN data) with other sympatric frugivorous species. For a given species A, sympatry with another species B was considered when at least 10% of the range of species A overlaps with the range of species B. This was done to reduce the noise induced by coarse identification of species range.

### Body mass and sympatry

Body mass may itself be affected by species sympatry. Yet, as body mass was used to compute some relative brain size, to control for the potential confounding effect due to a relationship between sympatry and body mass, we repeated all model fitting (models of trait evolution and PGLS) with body mass as the output variable.

### Models of species diversification

Next, we also investigated the correlations between primate diversification rates and brain sizes or levels of sympatry to better understand the impact of cognition and interactions between primates on their evolutionary success. To do so, we inferred how primates diversified over time and across lineages. Lineage-specific net diversification rates (defined as speciation minus extinction rates) were estimated using an updated version of the *ClaDS* algorithm (Maliet, Hartig, and Morlon 2019) based on data augmentation techniques (Maliet and Morlon 2021). Particularly, we used *ClaDS2*, a model with constant turnover (i.e. constant ratio between extinction and speciation rates; see Supplementary Material, Primate diversification rate over time, for further explanations). We extracted the mean diversification rates through time and the lineage-specific diversification rate of each extant species.

We fitted Gaussian Pagel’s lambda phylogenetic regressions of the different relative brain sizes against the net diversification rates. Because assumptions for a frequentist-based approach were unmet, we performed Bayesian-based inferences instead. We used the “MCMCglmm” function of the *MCMCglmm* package (Hadfield 2010). Each chain had a burn-in period of 5000 iterations, a total length of 50,000 iterations, and was sampled every 50 iterations. We used the least informative priors. Fixed priors were set to default values (Gaussian distribution of mean 0 and variance 10). Again, we used the mean of the brain trait values for the main model and assessed the sensitivity by re-running the model several times using non-averaged values.

To determine whether species sympatry was associated with lower or larger diversification rates, we fitted frequentist-based Gaussian Pagel’s lambda phylogenetic regressions with the lineage-specific diversification rate as the output variable and used the two metrics for describing sympatry (the number of frugivorous sympatric species and the average percentage of overlapping range with other sympatric frugivorous species) as the tested variables.

### Model implementation and stability

Details on the implementation, stability, and uncertainty of phylogenetic regressions are provided in Supplementary Material, Model stability). Necessary assumptions on the normal distribution of residuals and homoscedasticity were visually assessed and pointed to no violation (see Supplementary Material, Model assumptions). In addition, we did not observe correlation issues among predictors (Variance Inflation Factor, VIF*_max_* < 2, Mundry 2014).

## RESULTS

The dataset we gathered contained between 34 to 182 frugivorous primate species (depending on the brain region considered). When considering the sizes relative to the body mass of the different brain regions, we observed ample variations (Figure 3). For instance, the lemuriformes, which are known to prioritize smell compared with other primate species, have the largest relative MOB size (lemuriformes: mean ± SE = 0.23 ± 0.07, other: 0.12 ± 0.04, 3). Similarly, the highest relative sizes of the striatum were found in platyrrhini (platyrrhini: mean ± SE = 0.91 ± 0.07, other: 0.59 ± 0.07, 3), which are known to form poly-specific associations (callitrichines in particular, Heymann and Buchanan-Smith 2000). In terms of sympatry, we observed that on average (±SE) the considered primate species had 52% (±2) of their range overlapping with other species. That ranged from 0% of overlap (*Macaca nigra*), to 100% of overlap (*Cercopithecus pogonias, Alouatta pigra, Loris tardigradus, Hylobates moloch, Cercocebus galeritus, Presbytis melalophos, Semnopithecus entellus*). In terms of the distribution range, the considered primate species co-occurred on average with 6.38 (±0.39) other primate species, ranging from 0 other species to 21.

**Figure 3:**
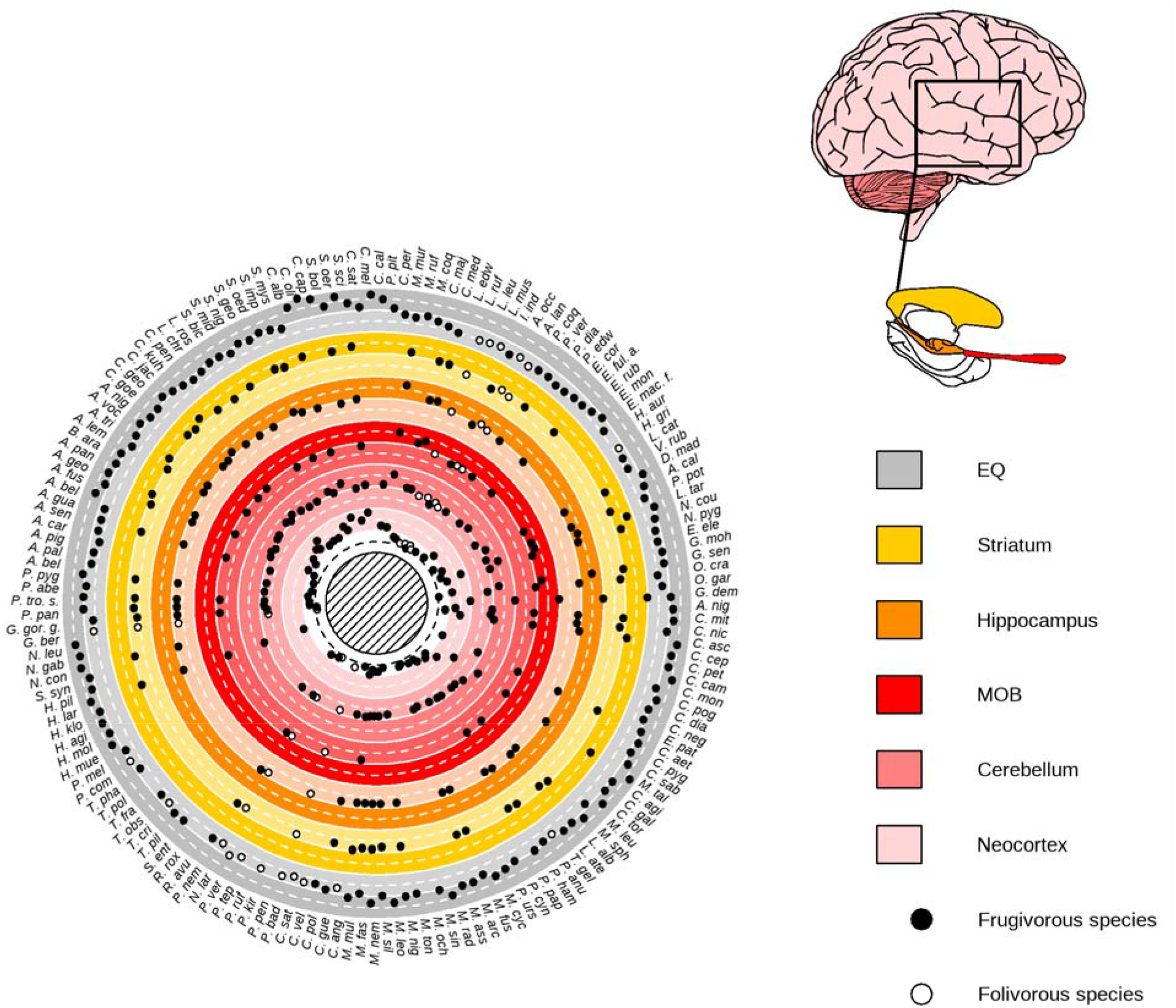
Variations in relative brain region sizes among extant frugivorous primates | (Left) Circular plot of the sizes of the different brain regions relative to the body mass. Colours indicate the rows for the different brain regions. The darker background emphasises when values are above average, while the lighter background emphasises when values are below average. The mean value (after scaling and based on one random sampling among possible values but see Supplementary Figure S3 for visualisation of measure variability) for the Encephalization Quotient (EQ) or relative size of brain regions, when available, is depicted by a plain circle for frugivorous species. The frugivorous threshold was fixed at 20% and the folivory threshold at 40%. (Right) The different studied brain regions (human brain as an illustration). In short, the MOB is involved in immediate olfactory information processing, the neocortex and the cerebellum support working memory and memory consolidation of immediate sensory information processing (Wiltgen et al. 2004; Koziol et al. 2014; Sokolov, Miall, and Ivry 2017), and the hippocampus supports working memory and long-term spatiotemporal memory (Burgess, Maguire, and O’Keefe 2002). The striatum is involved in social information processing (Báez-Mendoza and Schultz 2013).

To retrace past species sympatry between frugivorous lineages, we reconstructed primate biogeographic history (Figure 1) and their diet evolution. We found that the ancestors of primates were likely to be frugivorous and that folivory evolved more than five times, in particular in cercopithecidae and memuridae. Estimated diet history was consistent with fossil evidence (see Supplementary Material, Diet reconstruction, Supplementary Figure S4). We then modelled the evolution of the size of the whole brain (EQ) or separate brain regions (neocortex, cerebellum, MOB, hippocampus, and striatum) when considering species sympatry or not. Weighting each brain region by the whole-body mass and weighting by whole-brain size yielded similar results (see Supplementary Material, Weighting the size of brain regions by whole-brain size, Tables S1 and S4, Figures S5 and S6), therefore we only present the case when weighting by body mass. We found that models not considering species sympatry best described the evolutionary history of the EQ, the neocortex, and the cerebellum (Figures 3 and 4), two brain regions specifically involved in immediate sensory information processing (Wiltgen et al. 2004; Koziol et al. 2014; Sokolov, Miall, and Ivry 2017), and also in memory consolidation for the neocortex (Wiltgen et al. 2004). The fact that these biggest brain regions are best described by an Ornstein-Uhlenbeck process suggests a stabilisation towards an optimal relative size, which may illustrate the trade-off between the costs and benefits of brain development (Isler and van Schaik 2009). Yet, given the broad range of the study species, and the likely difference of their environments, this could hide a potentially multi-peak stabilisation that occurred towards different optima for different clades (Mahler et al. 2013). By contrast, density-dependent models considering species sympatry (DD_*lin*_ and DD_*exp*_) were best supported in the foraging-related and social-related brain regions respectively: the hippocampus, specialised in spatiotemporal memory (Burgess, Maguire, and O’Keefe 2002), and the striatum, involved in social interactions (Báez-Mendoza and Schultz 2013). The fact that we inferred positive rates *r* of density-dependence (Figure 4) suggested an acceleration of the evolutionary tempo of trait evolution together with an increased number of frugivorous sympatric lineages for the hippocampus and the striatum. The MOB, the brain region involved in sensory abilities, also tended to be best fitted by models considering sympatry as a whole. Yet, Brownian motion (BM) was as likely as density-dependent or matching competition models, preventing firm conclusions on whether sympatry affected or not MOB size evolution (Figures 3 and 4), especially since this coincided with the most reduced sample size we had (N_*species*_ = 34 to 39).

**Figure 4:**
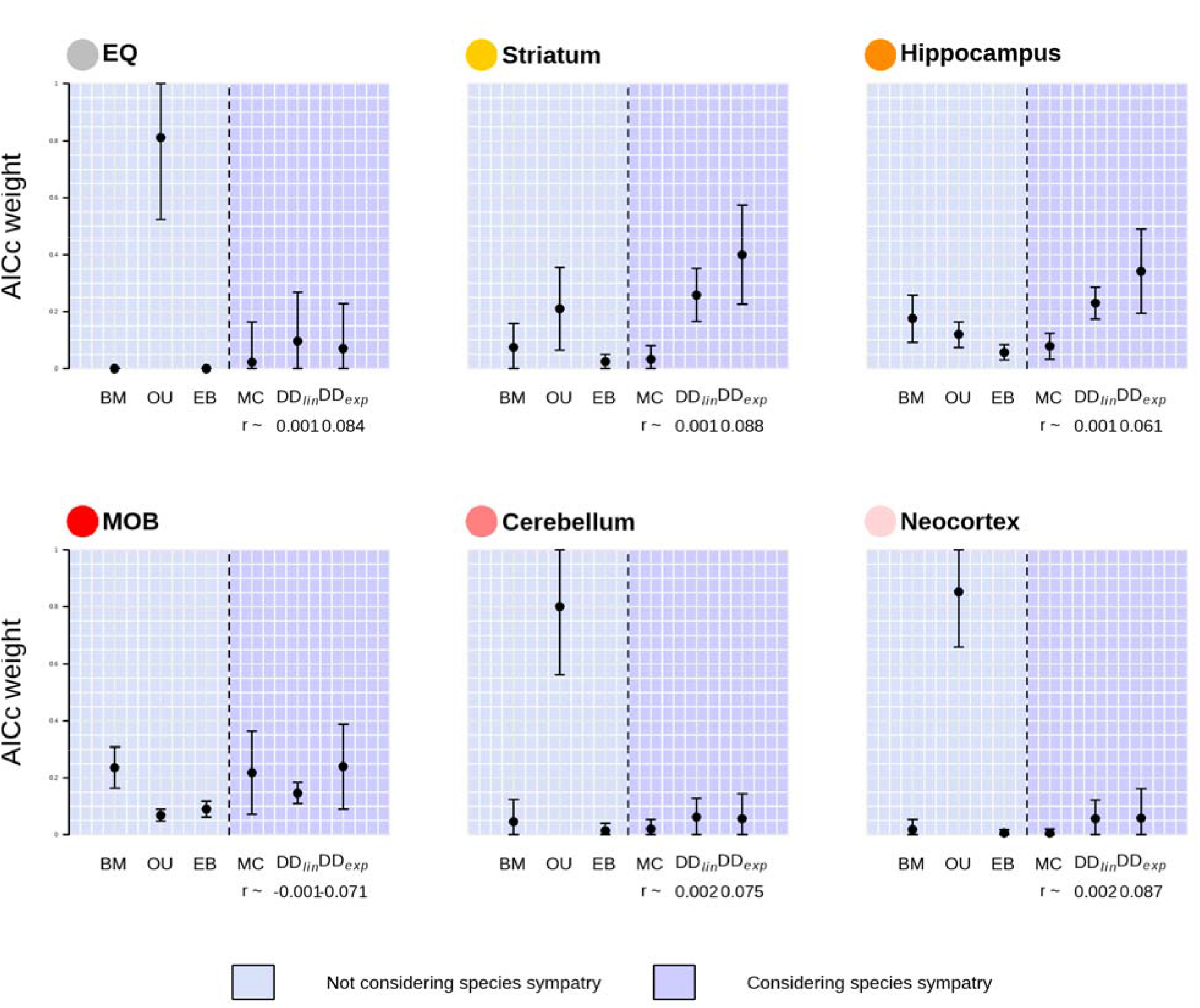
The evolution of the hippocampus and striatum in frugivorous primates are best fitted by models of trait evolution considering species sympatry | Plotted is the AICc weight, a measure of relative support for a given model, for models not considering species sympatry (BM, OU, EB) or considering species sympatry (MC, DD, DD). The points represent the average AICc weight obtained (when considering the six models from the same run), while the vertical bars indicate the standard deviation given all tested conditions (see Phylogenetic models of trait evolution: does species sympatry shape brain size evolution?).

Next, we assessed whether higher levels of species sympatry led to “bigger” or “smaller” brain regions. To do so, we fitted phylogenetic regressions in extant frugivorous species between the relative sizes of the different brain regions and two measures of sympatry (1) the average percentage of overlapping range with other frugivorous sympatric species, and (2) the number of such sympatric frugivorous species across their current entire distribution range. When weighting each brain region by body mass, the number of sympatric species never significantly influenced the relative brain sizes (Table 1; Supplementary Figures S7 and S10). Conversely, we found that the average percentage of overlapping range significantly correlated, or tended to correlate, with the relative size of brain regions that were better fitted with models considering sympatry: the hippocampus and the striatum (hippocampus: *t* = -1.94, p = 0.058; striatum: *t*= -2.26, p = 0.028). The correlations were all negative (hippocampus: est. = -0.39, CI95% = [-0.76,-0.01]; striatum: est. = -0.4, CI95% = [-0.77,-0.04]), which means that higher range overlap between sympatric species associates with lower relative size, insensitively to data and phylogenetic uncertainties (Supplementary Table S3, Supplementary Figures S7 and S10). The acceleration of the tempo of trait evolution with species sympatry (*r* > 0 in the density-dependent models) suggests that, compared with isolated species, sympatric species are subject to a positive selection towards smaller brains, and not to a less intense selection for advanced cognitive abilities.

**Table 1:**
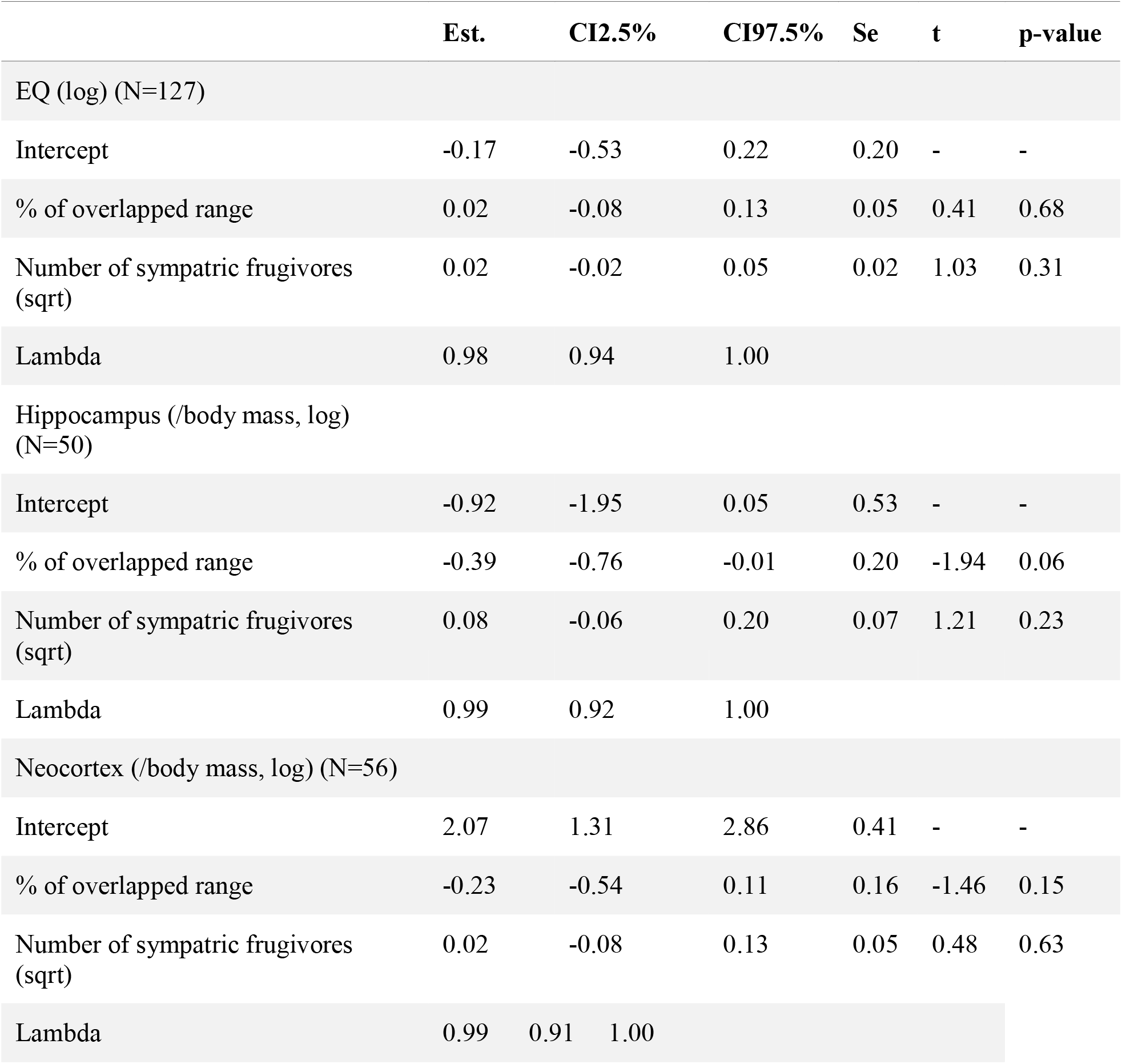
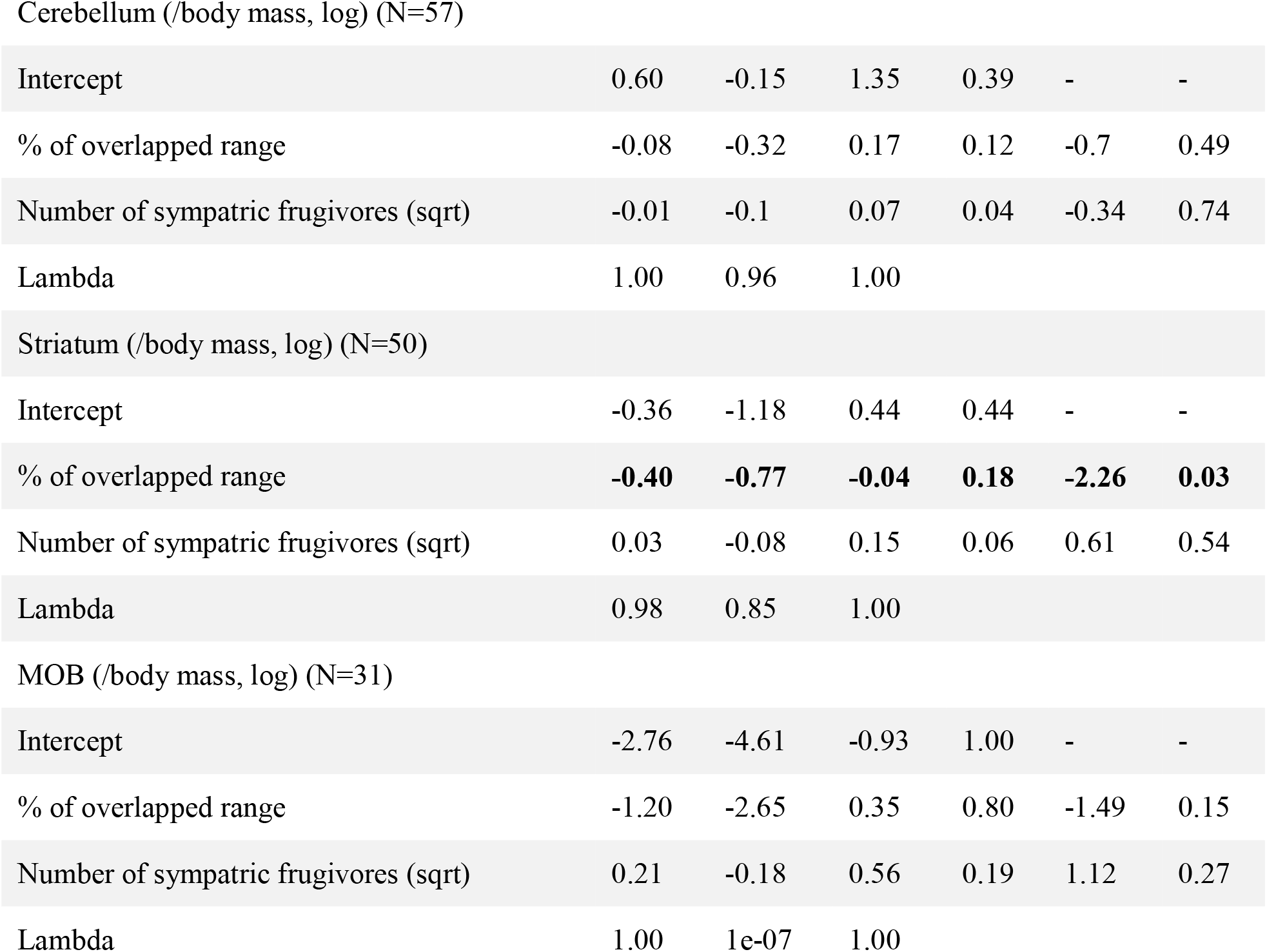
Species sympatry correlates negatively with the relative size of some brain regions in extant frugivorous primate species | Model estimates and significance of phylogenetic regressions to assess the relationship between relative brain sizes and species sympatry. Est.=Estimate, CI2.5%=Lower border of the CI95%, CI97.5%=Upper border of the CI95%, Se=Standard error, t=Statistics t-value. The brain regions (as well as the associated sample sizes) are indicated before each list of estimates. The transformations applied to variables are indicated between parentheses (logarithm, log, or square-root, sqrt), as well as the weighting by body mass (/body mass).

|  | Est. | CI2.5% | CI97.5% | Se | t | p-value |
| --- | --- | --- | --- | --- | --- | --- |
| EQ (log) (N=127) |  |  |  |  |  |  |
| Intercept | -0.17 | -0.53 | 0.22 | 0.20 | - | - |
| % of overlapped range | 0.02 | -0.08 | 0.13 | 0.05 | 0.41 | 0.68 |
| Number of sympatric frugivores (sqrt) | 0.02 | -0.02 | 0.05 | 0.02 | 1.03 | 0.31 |
| Lambda | 0.98 | 0.94 | 1.00 |  |  |  |
| Hippocampus (/body mass, log) (N=50) |  |  |  |  |  |  |
| Intercept | -0.92 | -1.95 | 0.05 | 0.53 | - | - |
| % of overlapped range | -0.39 | -0.76 | -0.01 | 0.20 | -1.94 | 0.06 |
| Number of sympatric frugivores (sqrt) | 0.08 | -0.06 | 0.20 | 0.07 | 1.21 | 0.23 |
| Lambda | 0.99 | 0.92 | 1.00 |  |  |  |
| Neocortex (/body mass, log) (N=56) |  |  |  |  |  |  |
| Intercept | 2.07 | 1.31 | 2.86 | 0.41 | - | - |
| % of overlapped range | -0.23 | -0.54 | 0.11 | 0.16 | -1.46 | 0.15 |
| Number of sympatric frugivores (sqrt) | 0.02 | -0.08 | 0.13 | 0.05 | 0.48 | 0.63 |
| Lambda | 0.99 | 0.91 | 1.00 |  |  |  |
| Cerebellum (/body mass, log) (N=57) |  |  |  |  |  |  |
| Intercept | 0.60 | -0.15 | 1.35 | 0.39 | - | - |
| % of overlapped range | -0.08 | -0.32 | 0.17 | 0.12 | -0.7 | 0.49 |
| Number of sympatric frugivores (sqrt) | -0.01 | -0.1 | 0.07 | 0.04 | -0.34 | 0.74 |
| Lambda | 1.00 | 0.96 | 1.00 |  |  |  |
| Striatum (/body mass, log) (N=50) |  |  |  |  |  |  |
| Intercept | -0.36 | -1.18 | 0.44 | 0.44 | - | - |
| % of overlapped range | **-0.40** | **-0.77** | **-0.04** | **0.18** | **-2.26** | **0.03** |
| Number of sympatric frugivores (sqrt) | 0.03 | -0.08 | 0.15 | 0.06 | 0.61 | 0.54 |
| Lambda | 0.98 | 0.85 | 1.00 |  |  |  |
| MOB (/body mass, log) (N=31) |  |  |  |  |  |  |
| Intercept | -2.76 | -4.61 | -0.93 | 1.00 | - | - |
| % of overlapped range | -1.20 | -2.65 | 0.35 | 0.80 | -1.49 | 0.15 |
| Number of sympatric frugivores (sqrt) | 0.21 | -0.18 | 0.56 | 0.19 | 1.12 | 0.27 |
| Lambda | 1.00 | 1e-07 | 1.00 |  |  |  |

Furthermore, we also investigated the influence of sympatry on body mass, which could be a confounding explanation for the observed patterns (Smaers et al. 2021). We found that the evolutionary history of body mass is best explained by MC models (Supplementary Figure S6). In other words, species in sympatry tend to diverge towards being large or small. There is thus no overall linear relationship between body mass and sympatry, explaining why we found no effect of sympatry variables on body mass in PGLS; Supplementary Tables S2 and S7).

Finally, we investigated the evolutionary consequences of cognition and species sympatry by evaluating whether brain sizes and levels of sympatry correlated with the lineage-specific net diversification rates of primates (Figure 5). Overall, species diversification rates particularly boomed in the early and late Miocene, around 25 and 11 Myr ago (Figure 5). When accounting for phylogenetic dependence, no significant relationship between the net diversification rate and the relative size of brain regions was found in extant primates (Table 2, Supplementary Figure S8; see robustness in Supplementary Table S5). Although diversification was uncorrelated with brain size in frugivorous primates, it was influenced by the sympatry context. In particular, phylogenetic regressions highlighted a negative effect of the number of sympatric species on the diversification rate (est. = -5.04e-03, CI95% = [-0.01,1.34e-04], t = 2.56e-03, p < 0.001, Table 3, Supplementary Figure S9, see robustness in Supplementary Table S6). In other words, the higher the number of sympatric species, the lower the diversification rate.

**Figure 5:**
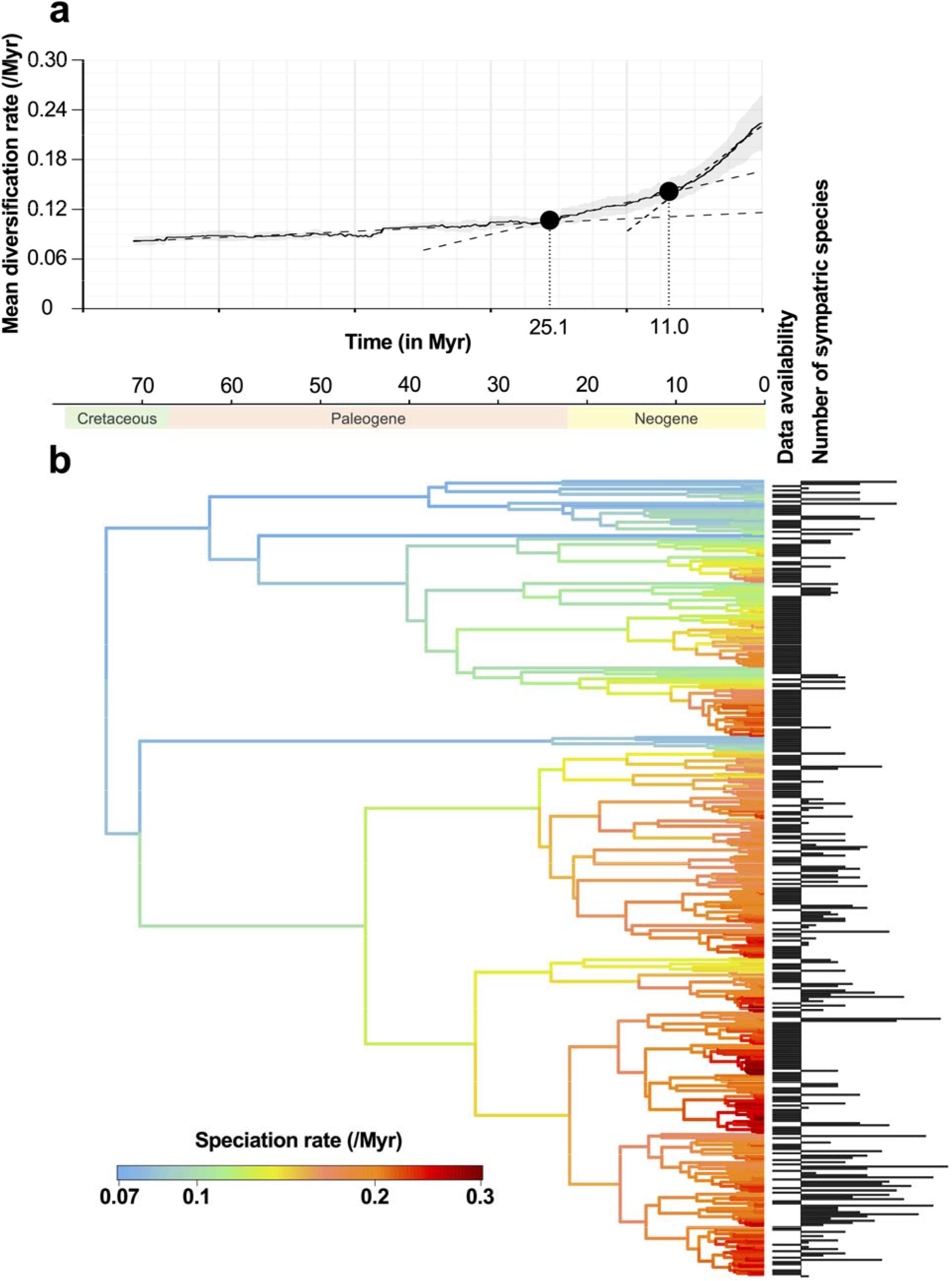
Species sympatry in primates is negatively associated with species diversification | a) Net diversification rate over time in the primate taxon. The average diversification rate estimated based on an assumed sampling fraction of primate species ranging from 60 to 95% (at a step of 10%; then 5% from 90%) is depicted by the plain line. The grey background depicts the standard deviation. The two breakpoints, depicted by the plain dots and the vertical dotted bars, were estimated based on a three-linear regression segmentation using the *strucchange* package [Zeileis et al. (2002); Zeileis et al. (2003); Zeileis (2006); see the vignette package for statistical details]. The three fitted regressions are displayed by the dashed lines. The choice of two breakpoints was first assessed by choosing the number of breakpoints minimising the Bayesian Information Criterion. b) The branches of the primate phylogenetic trees are colored according to their speciation rates estimated using ClaDS2: red colors indicate lineages that speciate frequently, whereas blue colors indicate lineages that rarely speciate. At the tips of the tree, we reported for each extant species whether we had data on its biogeography (column ‘Data availability’) and if so, we indicated the number of sympatric species it has. Using PGLS, we found a significant relationship between lineage-specific diversification rates and the numbers of sympatric species at present.

**Table 2:** Relative brain sizes do not correlate with primate species diversification in frugivorous primates | Model estimates and significance of Bayesian phylogenetic regressions to assess the correlation between the net diversification rates and the relative brain sizes. Est.=Estimate, HDP2.5%=Lower border of the 95% Highest Posterior Density, HDP97.5%=Upper border of the 95% Highest Posterior Density, Eff. Samp.=Effective sample (adjusted for autocorrelation). The brain regions (as well as the associated sample sizes) are indicated before each list of estimates. The (log) indicates log-transformed variables, while the (/body mass) indicates variables weighted by body mass.

|  | Est. | HDP2.5% | HDP97.5% | Eff. Samp. | pMCMC |
| --- | --- | --- | --- | --- | --- |
| Diversification EQ (N=148) |  |  |  |  |  |
| Intercept | 0.12 | 0.08 | 0.16 | 900.00 | - |
| EQ (log) | 0.02 | -7.91e-03 | 0.05 | 789.25 | 0.15 |
| Lambda | 0.83 | 0.76 | 0.9 |  |  |
| Diversification Hippocampus (N=61) |  |  |  |  |  |
| Intercept | 0.13 | 0.09 | 0.18 | 900.00 | - |
| Hippocampus (/body mass, log) | 9.10e-03 | -9.48e-03 | 0.03 | 900.00 | 0.34 |
| Lambda | 0.73 | 0.6 | 0.85 |  |  |
| Diversification Neocortex (N=67) |  |  |  |  |  |
| Intercept | 0.1 | 0.04 | 0.17 | 991.53 | - |
| Neocortex (/body mass, log) | 7.26e-03 | -0.02 | 0.03 | 900.00 | 0.56 |
| Lambda | 0.74 | 0.6 | 0.86 |  |  |
| Diversification Cerebellum (N=68) |  |  |  |  |  |
| Intercept | 0.12 | 0.07 | 0.16 | 900.00 | - |
| Cerebellum (/body mass, log) | 3.94e-03 | -0.02 | 0.03 | 989.21 | 0.76 |
| Lambda | 0.74 | 0.6 | 0.86 |  |  |

| Diversification Striatum (N=61) |  |  |  |  |  |
| --- | --- | --- | --- | --- | --- |
| Intercept | 0.12 | 0.08 | 0.17 | 900.00 | - |
| Striatum (/body mass, log) | 9.11e-03 | -0.01 | 0.03 | 900.00 | 0.44 |
| Lambda | 0.73 | 0.59 | 0.85 |  |  |
| Diversification MOB (N=37) |  |  |  |  |  |
| Intercept | 0.11 | 0.05 | 0.17 | 900.00 | - |
| MOB (/body mass, log) | -4.79e-03 | -0.02 | 0.01 | 900.00 | 0.59 |
| Lambda | 0.65 | 0.46 | 0.83 |  |  |

**Table 3.** : Species sympatry slowdowns primate diversification | Model estimates and significance of phylogenetic regressions to assess the correlation between diversification rate and species sympatry. Est.=Estimate, CI2.5%=Lower border of the CI95%, CI97.5%=Upper border of the CI95%, Se= Standard error, t= Statistics t-value. The brain regions (as well as the associated sample sizes) are indicated before each list of estimates. The transformation (logarithm or square-root) is indicated in parentheses by the abbreviation (log or sqrt).

|  | Est. | CI2.5% | CI97.5% | Se | t | p-value |
| --- | --- | --- | --- | --- | --- | --- |
| Diversification Sympatry (N=128) |  |  |  |  |  |  |
| Intercept | 0.15 | 0.10 | 0.2 | 0.03 | - | - |
| % of overlapped range | -5.40e-03 | -0.02 | 9.35e-03 | 8.14e-03 | -0.66 | 0.51 |
| Number of sympatric frugivores (sqrt) | **-5.04e-03** | **-0.01** | **1.34e-04** | **2.56e-03** | **-1.97** | **0.05** |
| Lambda | 0.96 | 0.89 | 0.99 |  |  |  |

## DISCUSSION

### Sympatry of frugivorous primate species impacts the evolution of their hippocampus and striatum sizes

Although developmental constraints can induce correlation in the evolution of different brain regions (Gómez-Robles, Hopkins, and Sherwood 2014), brain regions underpin different cognitive functions and can thus be under different, independent, selective pressures (Barton and Harvey 2000). The functional regionalisation is for instance evidenced here by the differences in relative sizes across lineages in the MOB, with larger sizes in the lemuriformes that mostly rely on smell to forage. Our analyses also highlighted such differences in evolutionary trajectories for the different brain regions by the variations in the best-fit models of size evolution. We indeed show that sympatry is one factor that affects the evolutionary regime under which only some brain regions evolve: although the brain as a whole (i.e., Encephalization Quotient, EQ), the cerebellum and the neocortex were not significantly affected by species sympatry, the latter nonetheless induced a change in the relative sizes of the hippocampus and the striatum.

Our results were similar whether the relative sizes were obtained by weighting by body mass or whole-brain size. Theoretically, the two weighting methods are expected to give insights into differences in resource allocation between body tissues or differences in resource allocation within the brain tissue, respectively. Nonetheless, we observed no differences in the response to sympatry by weighting by body mass or whole-brain size, despite weak to moderate correlations between sizes relative to the body mass and sizes relative to the whole-brain size (see Supplementary Figure S5). Therefore, our results suggest that species sympatry mostly affected within-brain energy partitioning (as weighting by body mass can depict reallocation changes towards the whole brain or only within the brain), and in particular reallocations to two relatively small brain regions: the hippocampus and the striatum.

The hippocampus and the striatum are brain regions involved in individual-based and social-based information processing, pinpointing that these two components might be under strong selection in primates (DeCasien, Williams, and Higham 2017; Powell, Isler, and Barton 2017; González-Forero and Gardner 2018). The hippocampus in particular may have played a considerable role in the evolution of primate-like behaviours (Schilder, Petry, and Hof 2020), driven by the changes that primates faced in their ecological environment (e.g. the spatiotemporal distribution of the food, DeCasien and Higham 2019), or by the social environment that they faced (e.g. the number of conspecifics they interact with, Todorov et al. 2019; DeCasien and Higham 2019).

The fact that the hippocampus, particularly relevant to processing and memorising spatiotemporal information, is negatively sensitive to sympatry, is consistent with the idea of an effect of sympatric species on resource spatiotemporality (Hypothesis 1). Competition is generally the first-thought factor to describe community structures (de Almeida Rocha et al. 2015) because it might affect the environment in which species evolve. We show that a higher level of sympatry is associated with smaller sizes of the hippocampus (following Prediction 1.2), while the neocortex and cerebellum are not, at odds with the hypothesis that this is the mere consequence of corticalisation (Wiltgen et al., 2004). This suggests that indirect competition for food might contribute to convoluting the environment, and such an over-complexification of the resource spatiotemporality may render cognitive foraging not advantageous anymore. As a result, it might even generate a selection for smaller brain regions involved in foraging. In parallel with the complexification of their environment, species might have also narrowed frugivorous primates’ niches, which might have synergistically affected their cognition. Indeed, the support for matching competition when modelling the evolution of the body mass of frugivorous primates indicates that sympatric primate species tended to diverge in terms of body mass, evolving towards lower or higher body mass in sympatry. This is consistent with the idea of niche partitioning in sympatry, e.g. where light primate species would occupy the canopy layer, while heavier primate species would occupy the ground. Such niche partitioning may be accompanied by dietary specialisation (Schreier et al. 2009) and impact cognitive abilities (Aristide et al. 2016), as dietary specialisation often requires lower cognitive abilities and thus smaller brain region sizes.

In contrast to direct/indirect competition, potential indirect facilitation between species due to “social” cues (Hypothesis 2), is ruled out by the absence of an effect of sympatry on brain regions involved in immediate sensory information processing (e.g. cerebellum or neocortex). This absence of effect can stem from two possibilities. Either foragers do not exploit cues left out by sympatric heterospecifics, or they discarded environment cues in favour of social cues, as it has been shown that foragers tend to use social information over environmental (i.e. personal) information, in particular in non-perfectly predictable environments (Rafacz and Templeton 2003; Dunlap et al. 2016). In both cases, stimulation intensity of the MOB, the cerebellum, or the neocortex would somehow remain equivalent when in sympatry or not. Further work should explicitly test for these possibilities.

As expected (Hypothesis 3), the striatum size was relatively larger in callitrichines, particularly known to form mixed-species groups (Heymann and Hsia 2015). Yet, overall, the striatum size was negatively affected by sympatry. This puzzle might take root in secondary, but key, functions supported by the striatum, namely reward expectation, goal-directed behaviour, and planning abilities (Johnson, Meer, and Redish 2007). These three functions might as well be advantageous when foraging. As for the hippocampus, then, the increase in environmental unpredictability could diminish the benefits of these future-oriented skills.

### Sympatry of frugivorous primate species is associated with slower diversification

Given the context-dependence of the direction of selection (towards bigger sizes when sympatry is low, smaller sizes otherwise), there is no surprise that we do not observe a correlation between the net diversification rate and the three brain regions affected by species sympatry. Surprisingly however, we found no positive association between the net diversification rate and the EQ, the cerebellum, or the neocortex, which were insensitive to species sympatry. By contrast, a positive association between brain size and diversification was also found in birds (Sayol et al. 2019) given that bigger brains act as a buffer to environmental challenges (Sol et al. 2007). A visual inspection of the regressions evidenced a positive trend if not considering phylogeny (EQ and neocortex, Supplementary Figure S8). Sudden encephalisation in primates is clearly associated with a limited number of closely related species (DeCasien, Williams, and Higham 2017; Melchionna et al. 2020). Thus, this limits the statistical power of our analyses accounting for phylogenetically relationships, as we cannot decipher whether larger brain size and faster species diversification result from a true biological link or appeared simultaneously but independently. This means that a positive association between brain size and species diversification remains a likely possibility (as previously suggested in primates, Melchionna et al. 2020). Species sympatry, however, induced a significant slowdown in primate diversification, a density-dependence trend frequently observed in many tetrapod clades (Condamine, Rolland, and Morlon 2019). This frames coherently with a competitive scenario, where the tempo of species diversification decreases when ecological niches are progressively filled up (Rabosky and Lovette 2008). Species competing for resources are thought to contribute to limiting competitors’ range (Price and Kirkpatrick 2009), hence constraining population size and diversification rate (Pigot and Tobias 2013).

### Limits

The use of brain size as a proxy for cognition is a central debate with no optimal solution (see grounded criticism from Deaner, Nunn, and van Schaik 2000; Healy and Rowe 2007; Logan et al. 2018; van Schaik et al. 2021). The current flourishment of consortia, allowing for much more detailed and standardised anatomical measurements (e.g. in primates: Milham et al. 2018), or standardised behaviourally-explicit comparisons (e.g. on captive, Many Primates et al. 2019; or wild primates, Janmaat et al. 2021) might alleviate biases stemming from brain size analysis, but this will take time to generate large-enough datasets. In the meanwhile, brain size is a proxy much appreciated in practice, because of its easy accessibility for a “large” number of species, while the multifaceted aspect of cognition can simply be taken into account by considering the brain as a mosaic of singular and independent regionalised areas that are cognitively specialised. Yet, this approach implies collating datasets from different sources, which can induce a high intra- and inter-species variability, as the methodology has changed over time (discussed in Navarrete et al. 2018). We thus sampled measures among the available datasets and repeated the analysis several times to account for the variability of the data sets. The results remained robust to this approach. This supports the idea that intra-species variation is negligible compared with inter-species variation, a necessary condition for brain size to be an honest signal of cognitive differences across species, despite differences in measurement protocols.

Not only can the nature of the data be biased, but the methods themselves can suffer from significant limitations. In our case, these limitations are particularly true for the reconstructions of diet history, ancestral biogeography, or species diversification that we performed, which infer the most likely evolutionary histories based on observations at present alone. To verify the realism of these inferences, we confronted the diet history reconstruction with fossil evidence. Dental microwear textures in tooth fossils can indeed be used to reconstruct past diets and we found that our estimates of ancestral primate diets were consistent with the available fossils (Ramdarshan, Merceron, and Marivaux 2012; Merceron et al. 2009, see Supplementary Figure S4). Similarly, we found that our estimates of species diversification rates from molecular phylogenies were consistent with estimates of past primate diversity reconstructed from all available fossil data (Springer et al. 2012). In addition, the identified breakpoints matched estimates from previous studies (Arbour and Santana 2017; Springer et al. 2012). We did not use fossils for the biogeographic reconstruction because, although this may help the reconstruction, the overall benefices are minor (Wisniewski, Lloyd, and Slater 2022) and highly influenced by the fossil data quality (e.g. spatial coverage), which is still heavily limited and biased. We nevertheless accounted for the uncertainty in all reconstructions in our analyses by reporting results based on multiple sets of ancestral reconstructions.

Furthermore, the effectiveness of a method also depends on the proxy and associated definitions, which may impact the insights of such large-scale analyses. For instance, this issue is clearly illustrated with group size being used as a proxy of social complexity in social cognition (Dunbar and Shultz 2007), despite social complexity being multifactorial. This rejoins our major assumption of brain size as an honest signal for cognition, a multifaceted trait that is likely to be affected by diverse genes. In this study, further simplifications were considered: brain regions were associated with one main function, an overlap of diets was considered within the whole frugivorous primate guild, and sympatry was considered to be based as soon as spatial overlap occurred. Instead, each brain region can be considered as a Russian doll, with possibly redundant sub-functionalisation between spatially distant regions, frugivorous primates can be differentially selective in the fruits they eat (Campera et al. 2019), and species may not necessarily encounter each other at fine spatiotemporal scales (Deane et al. 2013). Although we cannot exclude that more accurate definitions might change our results, it is important to note that such a simplification is imposed by many constraints: from the sake of theorising to computational time. In addition, these large definitions also have some advantages. First, they can capture a broad variety of situations. For instance, simply defining sympatry as co-occurrence in a given biogeographic area enables considering both, direct and indirect interactions. Second, more stringent definitions might induce strong initial constraints that would eventually bias inferences on past history. For instance, the considered range areas to define sympatry are large and are unlikely to capture smaller-scale spatial segregation between species (e.g. due to habitat-specific segregation or vertical segregation). Yet, a finer mapping of species range may poorly reflect the “realised niche,” as primate populations suffer intense local extinction in recent years due to human activity (Estrada et al. 2017; Pavoine et al. 2019). Furthermore, we also took care of varying the stringency of our definitions (e.g. thresholds used to consider a species frugivorous based on the percentage of frugivory, thresholds used to consider two species as sympatric based on range overlap) and results were robust to such arbitrary choices (see Supplementary Material, Model stability).

## CONCLUSION

We showed that species sympatry is an important factor shaping the brain evolutionary history of frugivorous primates. It now seems crucial to scrutinise more carefully how sympatry fits with other socio-ecological variables that also influence brain size evolution in this clade (e.g. diet, group size, home range, etc., see DeCasien, Williams, and Higham 2017; DeCasien and Higham 2019; Powell, Isler, and Barton 2017). In addition, dietary overlap and food competition might not only happen between frugivorous primates but between any frugivores in the same area. In fact, it is very likely that any hypotheses on cognition evolution, generally discussed within species, could be broadened to a between-species context: foraging facilitation between species does exist (Olupot, Waser, and Chapman 1998; Havmøller et al. 2021), and so do polyspecific social associations (Porter 2001), as well as inter-species territory and resource defence (Drury, Cowen, and Grether 2020; Losin et al. 2016) or imitation and copying (Persson, Sauciuc, and Madsen 2018; Pepperberg 2002). Similarly, prey-predator races could shape selection on cognitive abilities (Shultz and Dunbar 2006). As Alice said, “It’s a great huge game of chess that’s being played—all over the world” (Carroll 1871, chap. II) and all individuals are just pieces to play with or against.

## ACKNOWLEDGEMENTS

We considerably value the help provided by J. Drury in making some scripts available functions in the *RPANDA* package in *R* and helping us run them, and that of M-C. Quidoz for assistance in using the CEFE cluster. We thank S. Benhamou and M. Clairbaux for discussions on spatial projections, M. Quéroué, V. Lauret, A. Caizergues, and C. Teplistky for feedback on Bayesian computations, and H. Morlon for discussions on models of trait evolution. We also acknowledge Editor F. Condamine, as well as P. Gonzalez, O. S. Todorov, and three other anonymous reviewers for their constructive comments on our manuscript that helped us improve it. Finally, this work could not have been possible without prior data collection from the IUCN Red List (primate ranging), the 10kTrees project (phylogenetic trees), and A. R. DeCasien and collaborators, L. E. Powell and collaborators, O. S. Todorov and collaborators, E. P. Willems and collaborators, F. Pearce and collaborators, A. F. Navarrete and collaborators, and C. C. Grueter who provided primate trait data we used as (supplementary) material with their articles. Their indirect input is therefore tremendous.

## AUTHORS’ CONTRIBUTION

BR conceived the study, gathered, cleaned, and analysed the data, drew the figures, wrote the first version of the manuscript, and subsequently revised it. BP-L implemented the ClaDS algorithm for our data, helped with running other analyses and drawing figures, and revised the manuscript multiple times.

## FUNDING

Both authors were supported by a doctoral grant from the *École Normale Supérieure*, Paris.

## CONFLICT OF INTEREST

The authors declare having no conflict of interest. All authors gave final approval for publication and agree to be held accountable for the work performed therein.

## DATA, SCRIPT, AND CODE AVAILABILITY

The data and codes that support the findings of this study are openly available at https://github.com/benjaminrobira/Meta_analysis_cognition_primates.

## Notes

https://github.com/benjaminrobira/Meta_analysis_cognition_primates

### Competing Interest Statement

The authors have declared no competing interest.

### Summary of Updates

/

