## Supplementary Material for "Primate sympatry shapes the evolution of their brain architecture"

^1^ CEFE, Univ Montpellier, CNRS, EPHE, IRD, Montpellier, France.
^2^ Eco-anthropologie (EA), Muséum National d’Histoire Naturelle, CNRS, Université de Paris, Musée de l’Homme, Paris, France.
^3^ Institut de Biologie de l’École Normale Supérieure (IBENS), École Normale Supérieure, CNRS, INSERM, Université PSL, Paris, France.
^4^ Institut de Systématique, Évolution, Biodiversité (ISYEB), Muséum National d’Histoire Naturelle, CNRS, Sorbonne Université, EPHE, Université des Antilles, Paris, France.

### Trait data collection

Collected data (Figure S1) were gathered within the published literature: DeCasien and Higham (2019) for the whole brain and all other brain regions mentioned (cerebellum, hippocampus, Main Olfactory Bulb (MOB), neocortex, striatum), from Powell, Isler, and Barton (2017) and from Powell, Barton, and Street (2019) for the whole brain, cerebellum, and neocortex size, from Todorov et al. (2019) for the hippocampus and neocortex size, from Grueter (2015) for the whole brain, and from Navarrete et al. (2018) for the whole brain, cerebellum, hippocampus, and striatum size. Body mass was obtained from DeCasien, Williams, and Higham (2017), from Powell, Isler, and Barton (2017), from Grueter (2015), and from Pearce et al. (2013). We obtained the percentages of frugivory and/or folivory from DeCasien, Williams, and Higham (2017), from Powell, Isler, and Barton (2017), and from (2013).

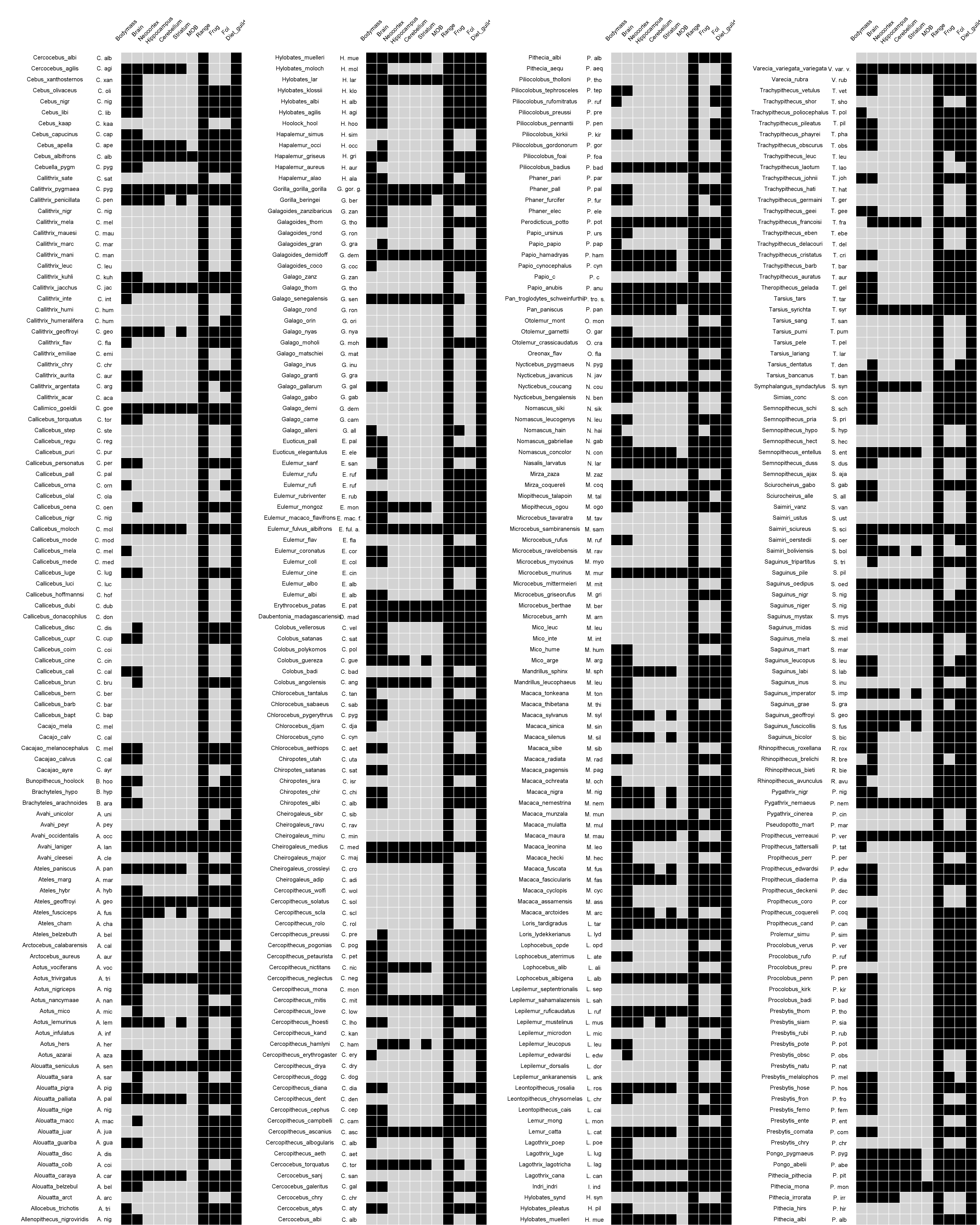

Figure S1: Data availability | Black boxes indicate data availability while grey boxes indicate the absence of data.

### Data variability

We present below the results of the assessments of data variability depending on the considered thresholds (for frugivory, folivory, or overlap) and the data set that is used, specifically related to distribution ranges, or anatomical/behavioural traits.

#### Sensitivity to variation in distribution ranges

The chosen biogeographic areas correspond to Central America, the North and the South of South America respectively, West Africa, Central Africa, and East/South Africa, East and West of Madagascar respectively, West Asia, Central/East Asia, South Asia, and the Asian Islands. The chosen scale for the areas is large because (i) retracing the history of a large number of areas necessitates considerable computational means. In addition, this drastically increases the computational time for fitting the phylogenetic models of brain trait evolution too. Furthermore (ii), all species and particularly primate species suffer(ed) from recent extinction (Pavoine et al. 2019), with a reduction of ranging areas at an unprecedented speed rate. Finer geographic characterization would therefore give too much weight to such anthropogenic effects that recently altered species distribution (e.g. evidenced in the North American fauna in Pineda-Munoz et al. 2021).

  The overlap between a species range and a biogeographic area was used to assess whether the species occupied the area or not. We chose overlaps of 10% (low threshold) or 30% (high threshold), to consider a species presence within a given area. We present now variations that occurred depending on the considered threshold.

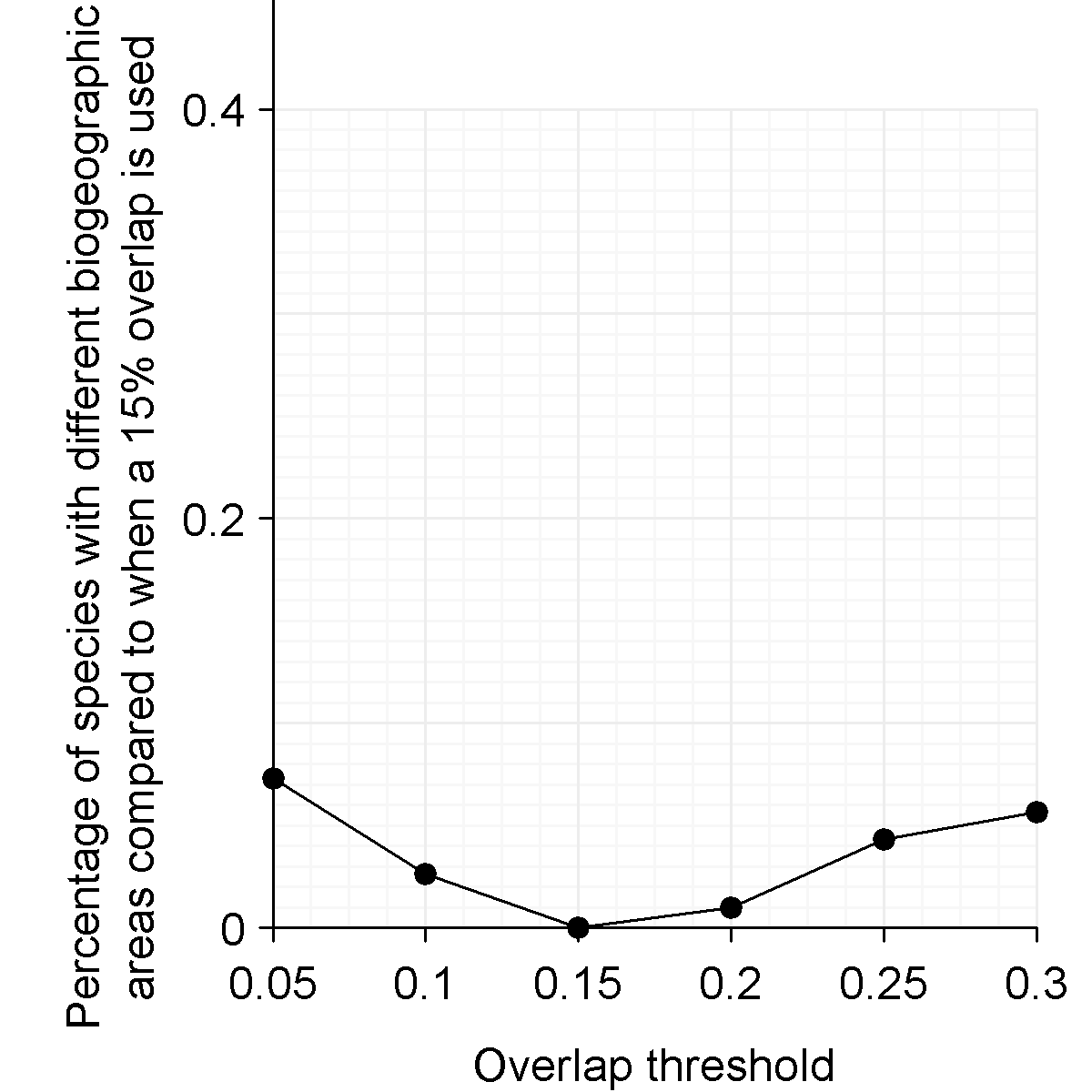

Figure S2: Percentage of species with differently identified biogeographic areas as a function of the overlap threshold (reference is an overlap threshold of 15%) | For a given species, a biogeographic area difference means that at least one biogeographic area considers the absence/presence of the species while this was not the case with the 15% threshold. 15% was chosen as the reference since halfway to the chosen maximum of 30%. 30% was chosen as the maximum because based on current observations, a species occupied at best three different biogeographic areas.

#### Sensitivity to variations in trait values

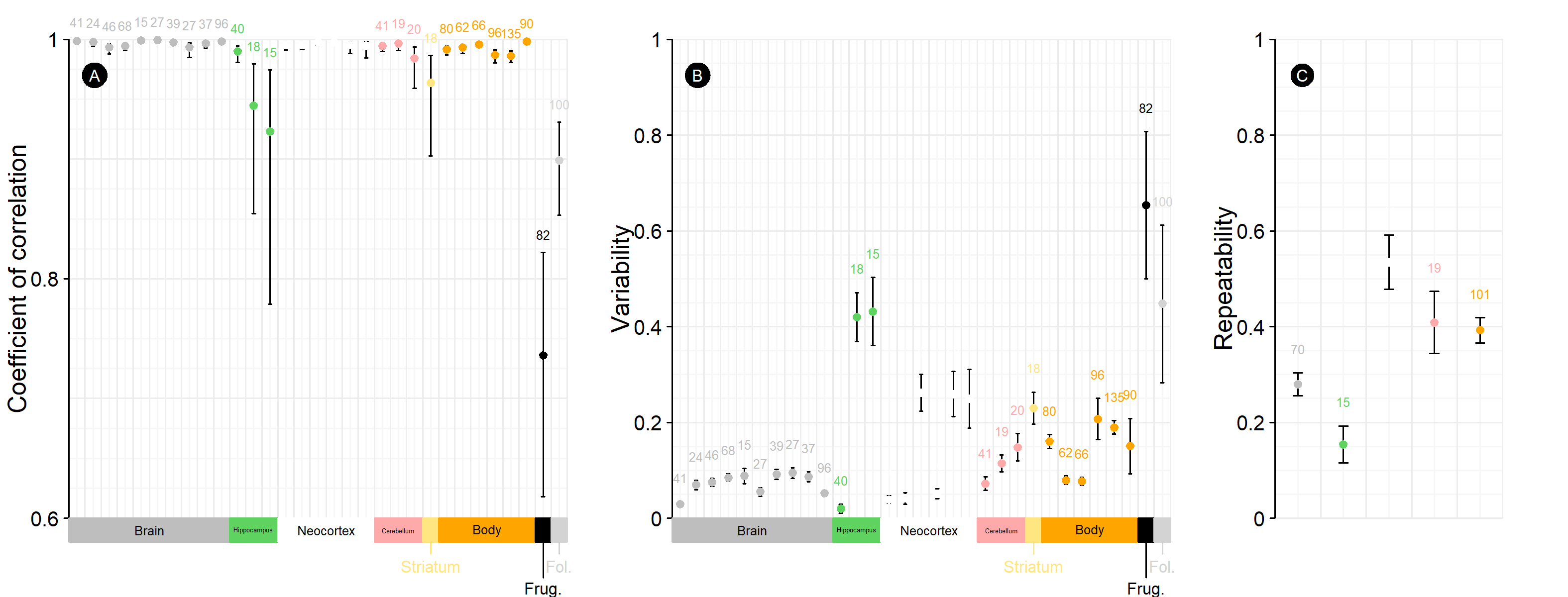

Figure S3: Variations in trait values among reference datasets | Colours are associated with a specific trait: Brain, hippocampus, neocortex, and cerebellum refer to the volume of the area (in mm$\boldsymbol{}^{\boldsymbol{3}}$), Body refers to the body mass (in g), Frug. indicates the percentage of frugivory and Fol. indicates the percentage of folivory. (A) Correlation: The points depict the coefficient of correlation while the bar depicts the 95% confidence interval (CI). (B) Variability: The points depict the average of the mean ratio $\boldsymbol{m}$ of the absolute of differences with paired values; If we reduce the equation, we have $\boldsymbol{m}\mathbf{=}\left| \left( \boldsymbol{v}_{\boldsymbol{1}}^{\boldsymbol{2}}\mathbf{-}\boldsymbol{v}_{\boldsymbol{2}}^{\boldsymbol{2}} \right) \right|\mathbf{/}\left( \boldsymbol{2}\boldsymbol{v}_{\boldsymbol{1}}\boldsymbol{v}_{\boldsymbol{2}} \right)$, where $\boldsymbol{v}_{\boldsymbol{1}}$ and $\boldsymbol{v}_{\boldsymbol{2}}$ are the two paired values from two different datasets and are different from 0. If $\boldsymbol{v}_{\boldsymbol{1}}$ and $\boldsymbol{v}_{\boldsymbol{2}}$ equal 0, then $\boldsymbol{m}\mathbf{=}\boldsymbol{0}$. If $\boldsymbol{v}_{\boldsymbol{1}}$ or $\boldsymbol{v}_{\boldsymbol{2}}$ equals 0 (case for the diet rates constrained between [0,1]), then we fixed the null value to 0.01. The bar depicts the standard error. (C) Repeatability: Repeatability was assessed for traits that were included in at least three datasets. Before calculation, traits were pondered *within* species by the *within* species max value. The point represents the mean repeatability $\boldsymbol{r}$ calculated as $\boldsymbol{\sigma}_{\boldsymbol{between}}^{\boldsymbol{2}}\mathbf{/}\left( \boldsymbol{\sigma}_{\boldsymbol{between}}^{\boldsymbol{2}}\mathbf{+}\boldsymbol{\sigma}_{\boldsymbol{within}}^{\boldsymbol{2}} \right)$, with the $\boldsymbol{\sigma}_{\boldsymbol{between}}^{\boldsymbol{2}}$ and $\boldsymbol{\sigma}_{\boldsymbol{within}}^{\boldsymbol{2}}$ corresponding the variance *between* or *within* species. The bar depicts the standard error. For all graphics, sample sizes are indicated above the upper value of the CI/error interval.

In addition to variability between datasets, we varied thresholds used to classify species diets. Illustratively, for example, if a species reached 30% of frugivory, and 65% of folivory (the rest being due, for instance, to insectivory), then, it would be categorized as “frugivorous” if low thresholds are considered, or as “folivorous” in case high thresholds are considered. If we suppose now another species with 30% of frugivory and 50% of folivory, but considered by DeCasien, Williams, and Higham (2017) as frugivorous, then, the species would be “frugivorous” for a low threshold, and still “frugivorous” with high a threshold.

  Frugivory was prioritized over folivory because we considered that since fruits are a highly palatable food source, they would be the key items that drive the foraging strategy (and associate consequence(s) on brain selection), even if less consumed. Additionally, to consider frugivory, we used a lower threshold than for folivory for two reasons. First, such a static percentage does not reflect potential seasonality in fruit-eating (e.g. Masi, Cipolletta, and Robbins 2009), which is generally shorter, hence a lower overall percentage of frugivory. Second, the percentage of frugivory is likely to be underestimated in part because primates generally spend more time feeding on leaves than fruits, while rates are often based on relative feeding time, or observation frequency at the individual or group unit of feeding events. Finally, the methodology to obtain this percentage could additionally vary (e.g. in addition to the two aforementioned estimations, one could also rely on the proportion of species targeted for their fruits/leaves). For all these reasons, we used two threshold levels (low, 20%, or high, 40%) to classify a species as frugivorous, as well as two threshold levels (low, 40%, or high, 60%) to classify a species as folivorous.

### Supplementary methods

#### **Spatial overlap assessment**

Overlaps of primate current ranges with biogeographic areas were calculated with the “gIntersection” function from the *rgeos* package (Bivand and Rundel 2021) applied to Mercator-projected data to get the overlapping contour, and the “area” function from the *geosphere* package (Hijmans 2021), applied directly on unprojected longitudinal-latitudinal data for area size calculation.

#### Primate diversification rate over time

Lineage-specific diversification rates were estimated using an updated version of the *ClaDS* algorithm (Maliet, Hartig, and Morlon 2019) based on data augmentation techniques (Maliet and Morlon 2021). This Bayesian approach considers rate heterogeneity by modelling small shifts in the rate at speciation events. In other words, the two new lineages are assumed to inherit new speciation rates that are sampled from a log-normal distribution with an expected mean value $log\left( \alpha\lambda\right)$ (where $\lambda$ represents the ancestral speciation rate and $\alpha$ is a trend parameter), and a standard deviation $\sigma$. Three independent chains were run until their convergence was validated by a Gelman-Rubin diagnostic criterion (Gelman and Rubin 1992). The analysis relied on the use of a consensus tree of primate phylogeny from Dos Reis et al. (2018). The latter provides a robust phylogenetic tree for 367 primate species (while the 10kTrees primate phylogeny has only 301 species).

  Such analysis necessarily depends on the fraction of sampled taxa (present in the phylogenetic tree) among all possible existing ones. Estrada et al. (2017) estimated that, given current knowledge, the primate clade should be composed of 504 species. This means that the current sampling fraction is around 73%. We thus parameterized the *ClaDS* algorithm with this value for the estimated sampling fraction. Yet, given that the extant number of primate species is subject to controversy, and because the estimated sampling fraction may affect diversification rate estimations, we replicated our analyses with a range of sampling fractions from 95% down to 60%. At the end of each run, we extracted the maximum *a posteriori* net diversification rate of each extant primate species, as well as the mean diversification rate (given all lineages) through time.

#### Evolutionary models

##### Model implementation

(i) Effect of sympatry on brain sizes

Models were fitted using the “phylolm” function from the *phylolm* package (Ho and Ane 2014), with the lambda parameter (with $\lambda$ indicating the strength of the phylogenetic signal, where $\lambda$=1 corresponds to Brownian Motion, i.e. the maximal influence of the phylogenetic history on the trait evolution) estimated by maximum-likelihood (argument “model” set to “lambda”). Bootstrapping over 1000 independent replicates was done to obtain confidence intervals. Other function parameters were set to default. Prior to fitting, covariates were transformed to reach more symmetrical distributions when adequate. Necessary assumptions on the normal distribution of residuals and homoscedasticity were visually assessed and pointed out no violation (see Supplementary methods Model assumptions). We did not observe any correlation issue among predictors either (VIF${}_{max}$ < 2, Mundry 2014).

(ii) Diversification and brain size

We could not compute phylogenetic regressions to link diversification and brain traits in frugivorous primates using a frequentist-based approach because it led to a violation of homoscedasticity. Instead, we fitted Bayesian phylogenetic regressions using the “MCMCglmm” function of the *MCMCglmm* package (Hadfield 2010). Each chain was based on a burn-in period of 5000 iterations, among a total of 50 000 iterations, and was sampled every 50 iterations. We used the least informative priors. Fixed priors were set to default values (Gaussian distribution of mean 0 and variance ${10}^{8}$). Priors on random effects and residuals were set to follow an inverse-Wishart distribution with a variance at a limit ($V$) of 1, and a degree of belief ($\nu$) of 0.02. We checked model convergence by fitting three chains, and calculated the Gelman-Rubin criterion (max value < 1.0042, Gelman and Rubin 1992), as well as checked autocorrelation (max absolute value < 0.07) using the respective “gelman.diag” and “autocorr.diag” functions from the *coda* package (Plummer et al. 2006). In Supplementary methods Model assumptions, we present traces and distributions of posterior estimates. We further checked the quality of the posterior by visually assessing the Q-Q plot of the posterior with that of a Gaussian distribution of mean 0 and sd 1 (see Supplementary methods Model assumptions). We present the estimate together with the 95% credibility interval centered on the mode (Highest Density Posterior, HDP), together with an MCMC p-value (pMCMC) that corresponds to the probability that the estimate ($\beta$) is positive if the mean estimate ($\hat{\beta}$) is negative (i.e. $P\left( \beta>0|\hat{\beta}<0 \right)$), or if the mean estimate is positive, the probability that the estimate is negative (i.e. $P\left( \beta<0|\hat{\beta}>0 \right)$).

(iii) Diversification and sympatry

We fitted phylogenetic regressions as explained in (i). In particular, verifications of model assumption and stability pointed out no source of worry (see Phylogenetic regressions: results, stability, and assumption).

##### Model stability

###### Dealing with data uncertainty and parameterisation sensitivity in models of trait evolution

In our analyses, uncertainty can stem from two sources. First, the reconducted evolutionary histories (in particular ancestral diet and biogeography) are susceptible to be uncertain at some points. Thus, we used 10 reconstructions for running each of our analyses.

  Second, for each species, trait estimates could vary slightly among datasets (see Supplementary Figure S3). Particularly, although correlations between measures from the different datasets seem good enough, there is a variation in absolute measurements (Supplementary Figure S3). We are aware that this could contribute to blurring results (Wartel, Lindenfors, and Lind 2019; Hooper, Brett, and Thornton 2021). In addition, this study is based on several arbitrary thresholds, namely (i) to assess species sympatry and (ii) to assess the species dietary guild which can cause sensitivity of the results to the chosen parameters. To account for these three sources of variability, we refitted several times the six models of trait evolution (BM, OU, EB, MC, DD${}_{lin}$ and DD${}_{exp}$) with (1) random samples of the dietary and brain traits in case of multiple values available (i.e. equal probability for each possible value to be selected), (2) used the low or high threshold for assessing frugivory, folivory, and species sympatry, and (3) various biogeography and dietary evolutionary history reconstructions.

  Each model thus represents the average of 10 (uncertainty on diet/ranging evolution) x 10 (uncertainty in brain/diet rate data) x 2 (geographic overlap threshold) x 2 (frugivory threshold) x 2 (folivory threshold) = 800 sub-models. We stopped computations when the calculation of the likelihood was excessively long (> 1 week). The final sample size thus was 730 models.

###### Assessing model stability in PGLS

We present below statistical indicators related to changes in estimates when re-fitting the model considering sub-samples (i.e. DfBetas and Cook’s distance), as well as when accounting for data variability (i.e. re-sampling among possible values given all datasets) or when using different parameterisation (i.e. “sampling fraction” of known species for diversification analyses).

  To assess frequentist model stability with regards to singular points, we computed the DfBetas (variation in estimates) by discarding one observation at a time of the “standard” dataset used to fit the main model, based on the consensus tree.

  To assess the sensitivity to (i) the variability in data and (ii) phylogeny uncertainty, we refitted the models using 50 phylogenetic trees among the 10000 possible trees from the 10kTrees project. For each of these trees, we fitted the model 50 times, allowing random sampling for data when we had multiple values (e.g. if body mass was provided by different datasets etc.). For the diversification analysis specifically, we also assessed the sensitivity to changes in primate sampling fraction by refitting the models for values ranging between 60 to 95% (as specified before), using the “standard” dataset and the consensus tree.

### Supplementary results

#### Diet reconstruction

We retraced the evolutionary history of frugivory versus folivory based on a continuous Markov process (extended Mk models; Bollback 2006) using Bayesian inference and internally setting up the prior probability of trait, but with no prior on the transition matrix.

We present in Supplementary Figure S4 the reconstruction of primate diets, considering only extant frugivorous and folivorous species. We compared it to fossil evidence, highlighting that the method used is consistent with the latter.

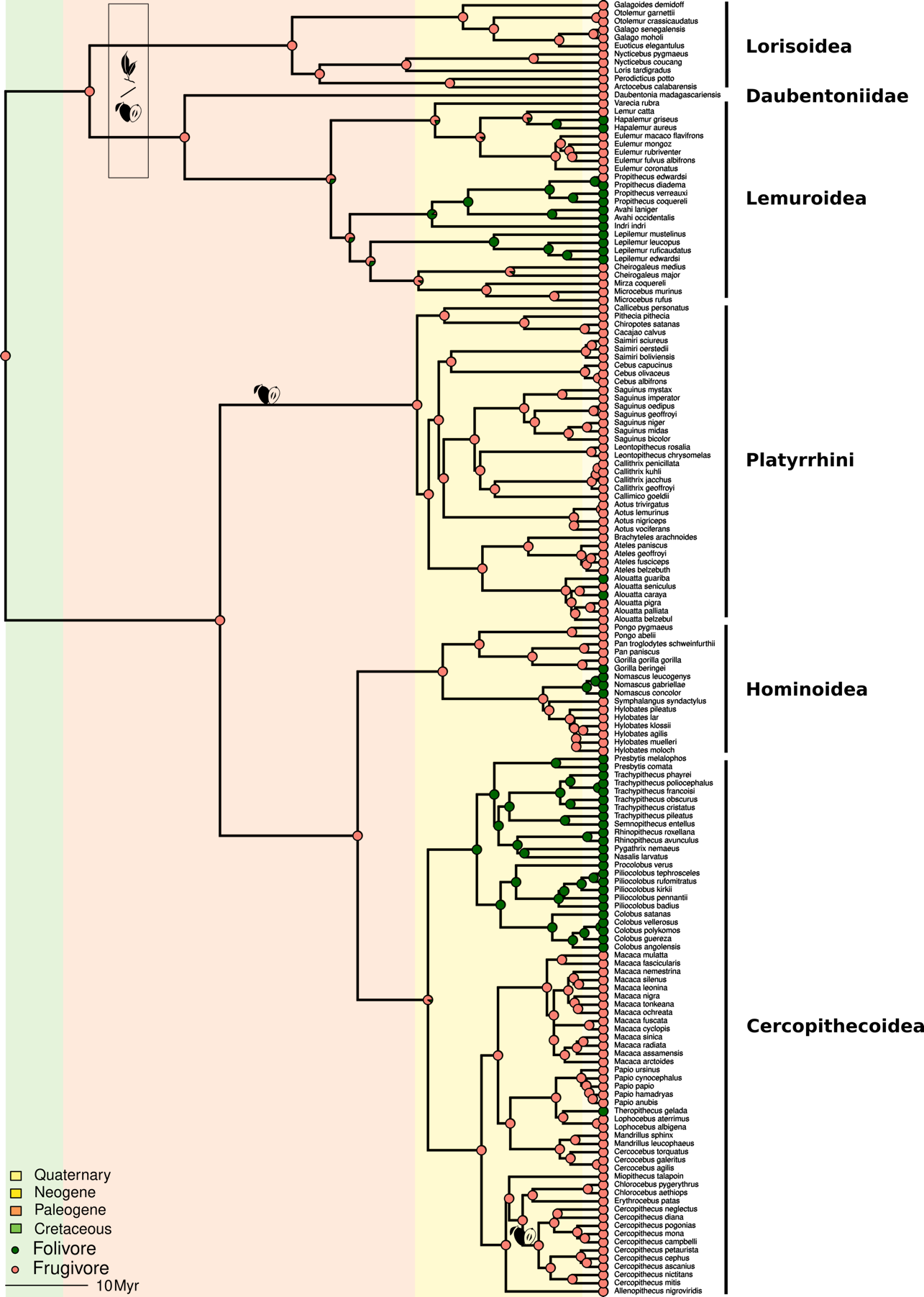

**Figure S4: Reconstruction of diet evolution along the primate phylogeny | A binary classification of diet (frugivory vs. folivory) was considered, with the highest threshold for frugivory and folivory classification used (see Methods). The diagram at each node of the phylogenetic tree indicates the estimated proportion of reconstructions in which the past species belonged to each dietary guild. The estimated diet of a few fossils (*Leptadapis magnus*, seasonal frugivorous/folivorous and *Pseudoloris pavulus*, frugivorous (Ramdarshan, Merceron, and Marivaux 2012), and *Mesopithecus*, frugivorous (fruit seeds, Merceron et al. 2009)) were added with a fruit or leaf symbol adequately.**

#### Weighting the size of brain regions by whole-brain size

We repeated the analyses considering relative brain size as the ratio between the sizes of the brain region of interest and the whole brain. We show that results are analogous to what is found when making the ratio with body mass instead, despite the correlations between the two ratios being not substantial (Supplementary Figure S5). Results are presented in Supplementary Figure S6 and Supplementary Table S1.

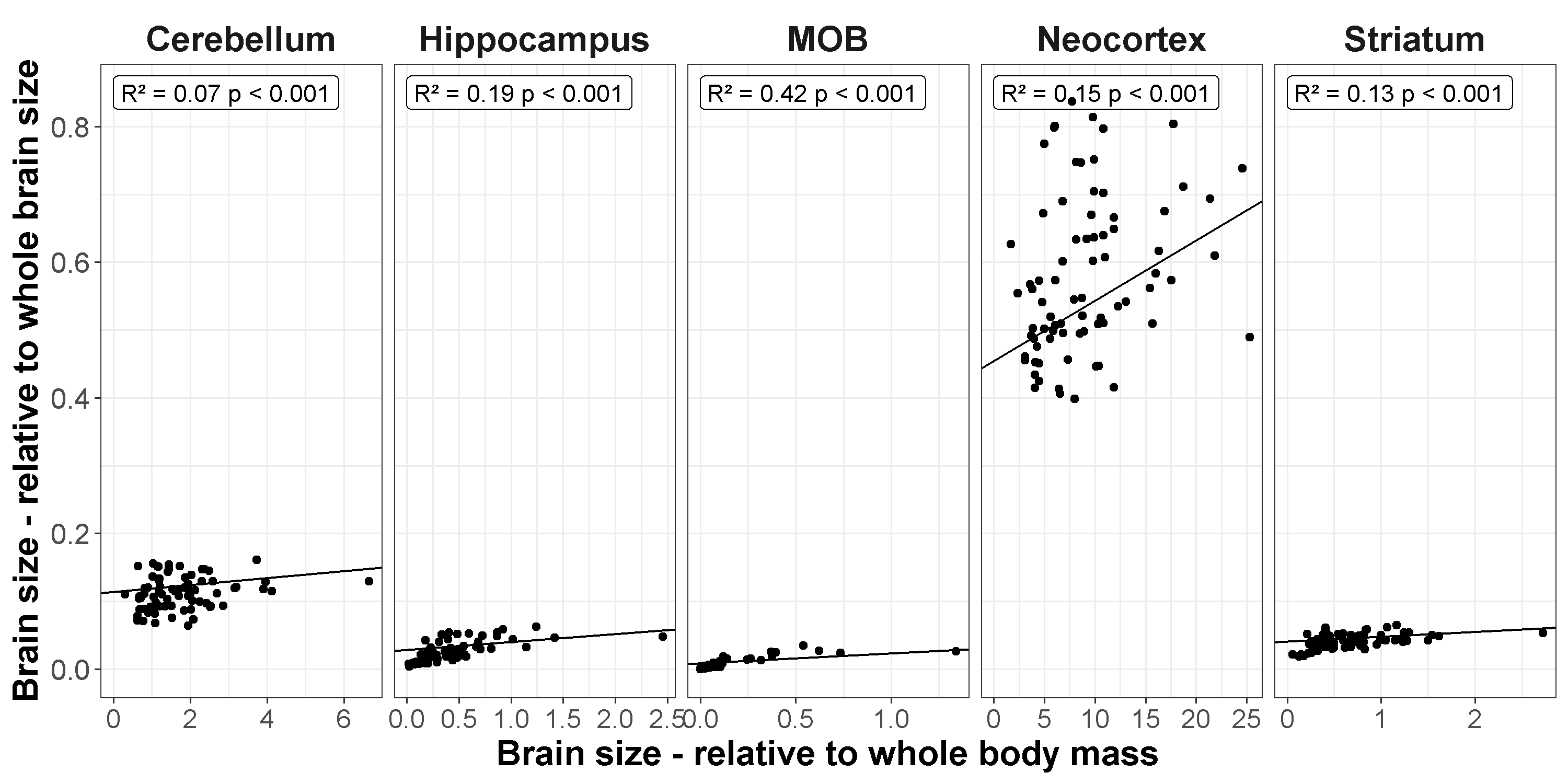

Figure S5: Relative sizes of brain regions based on body mass or whole-brain size do not substantially correlate | Each point represents one species. It was obtained by averaging the sizes of the brain regions, the whole-brain size, and body mass among all datasets.

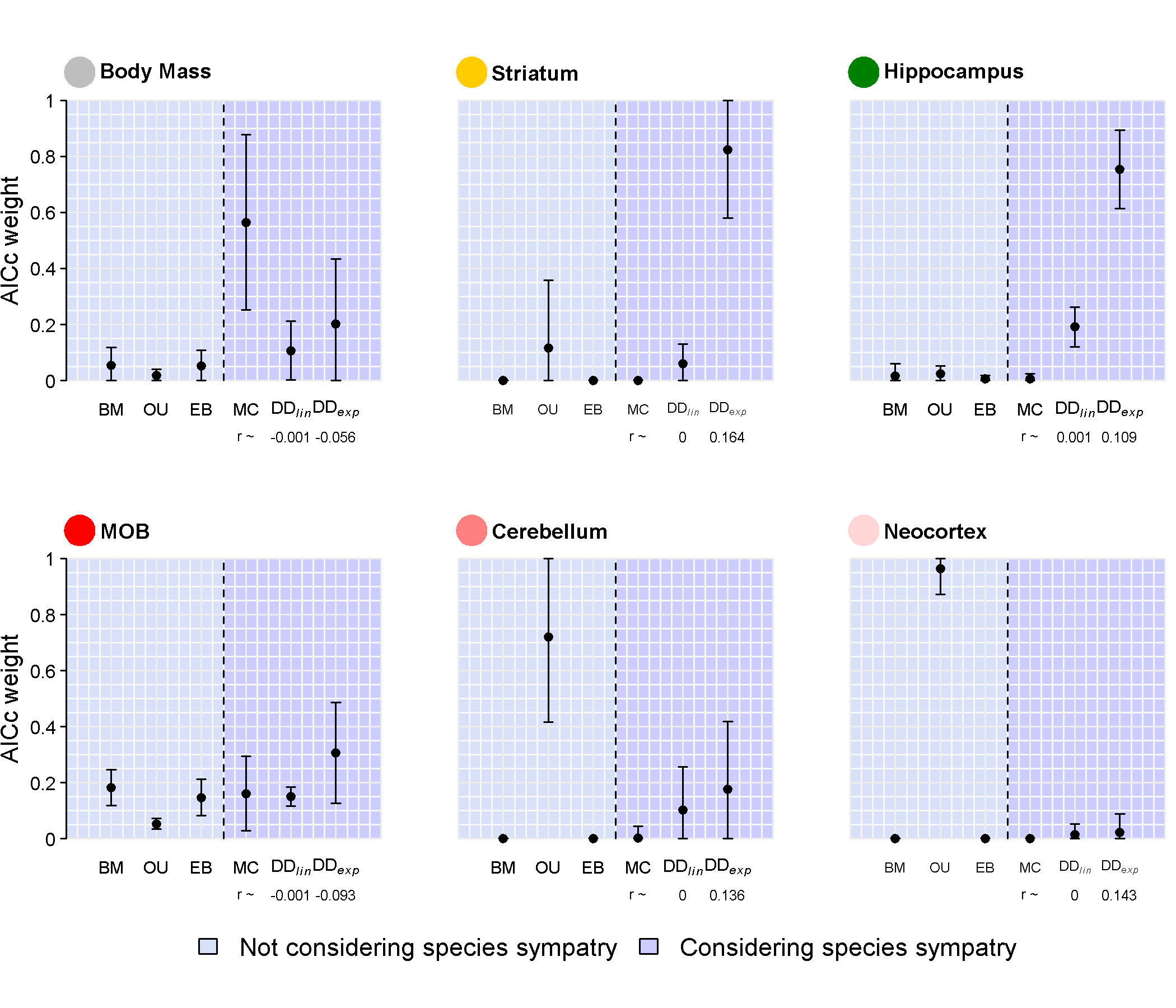

Figure S6: The evolution of body mass is best predicted by MC models, but estimated evolutionary patterns of the relative size of brain regions, when weighting by whole brain size, is unchanged compared to when weighting by body mass (Main Figure 4) | Plotted are the AICc weights, a measure of relative support for a given model, for models not considering species sympatry (BM, OU, EB) or considering species sympatry (MC, DD$\boldsymbol{}_{\boldsymbol{lin}}$, DD$\boldsymbol{}_{\boldsymbol{exp}}$). The points represent the average AICc weights obtained (when considering the six models from the same run), while the vertical bars indicate the standard deviation given all tested conditions. Depending on the brain region and the frugivory threshold we considered, the models were fitted on different sample sizes: EQ: 148 to 182, striatum: 56 to 63, MOB: 34 to 39, neocortex: 61 to 69, hippocampus: 56 to 63, cerebellum: 62 to 70 frugivorous species. For a given set of models (i.e. within a brain region), the sample was strictly identical, allowing within-set model comparisons.

Table S1: Species sympatry correlates negatively with the size of some brain regions of extant frugivorous primate species when weighting by the whole-brain size | Model estimates and significance of phylogenetic regressions to assess the relationship between relative brain sizes (weighted by the whole-brain size) and species sympatry. Est.=Estimate, CI2.5%=Lower border of the CI95%, CI97.5%=Upper border of the CI95%, Se=Standard error, t=Statistics t-value. The brain regions (as well as the associated sample sizes) are indicated prior to each list of estimates. The transformations applied to variables are indicated between parentheses (logarithm, log, or square-root, sqrt), as well as the weighting by the whole-brain size (/whole-brain size).

|  | **Est.** | **CI2.5%** | **CI97.5%** | **Sd** | **t** | **p-value** |
| --- | --- | --- | --- | --- | --- | --- |
| **Hippocampus (/whole-brain size, log) (N=50)** |  |  |  |  |  |  |
| Intercept | -3.52 | -4.13 | -2.96 | 0.32 | - | - |
| % of overlapped home range | -0.29 | -0.62 | 0.06 | 0.18 | -1.64 | 0.11 |
| Number of sympatric frugivorous (sqrt) | 0.05 | -0.05 | 0.14 | 0.05 | 0.98 | 0.33 |
| Lambda | 0.93 | 0.54 | 1.00 |  |  |  |
| **Neocortex (/whole-brain size, log) (N=56)** |  |  |  |  |  |  |
| Intercept | -0.76 | -0.95 | -0.56 | 0.10 | - | - |
| % of overlapped home range | 0.15 | -0.02 | 0.34 | 0.09 | 1.72 | 0.09 |
| Number of sympatric frugivorous (sqrt) | 0.03 | -0.01 | 0.07 | 0.02 | 1.47 | 0.15 |
| Lambda | 0.44 | 1e-07 | 0.76 |  |  |  |
| **Cerebellum (/whole-brain size, log) (N=57)** |  |  |  |  |  |  |
| Intercept | -2.18 | -2.4 | -1.96 | 0.11 | - | - |
| % of overlapped home range | 0.06 | -0.09 | 0.21 | 0.08 | 0.74 | 0.46 |
| Number of sympatric frugivorous (sqrt) | 0.01 | -0.02 | 0.05 | 0.02 | 0.71 | 0.48 |
| Lambda | 0.80 | 1e-07 | 0.96 |  |  |  |
| **Striatum (/whole-brain size, log) (N=50)** |  |  |  |  |  |  |
| Intercept | -3.03 | -3.29 | -2.76 | 0.15 | - | - |
| % of overlapped home range | -0.24 | -0.46 | -0.03 | 0.10 | -2.35 | 0.02 |
| Number of sympatric frugivorous (sqrt) | 0.02 | -0.03 | 0.07 | 0.03 | 0.66 | 0.51 |
| Lambda | 0.82 | 1e-07 | 0.99 |  |  |  |
| MOB (/whole-brain size, log) (N=31) |  |  |  |  |  |  |
| Intercept | -5.34 | -6.7 | -3.98 | 0.74 | - | - |
| % of overlapped home range | -0.84 | -1.91 | 0.30 | 0.59 | -1.43 | 0.16 |
| Number of sympatric frugivorous (sqrt) | 0.13 | -0.16 | 0.38 | 0.14 | 0.92 | 0.37 |
| Lambda | 1.00 | 1e-07 | 1.00 |  |  |  |

#### Visual representation of sympatry on relative brain size (weighting by whole-body mass)

(a) Phylogenetic regressions: effect of sympatry on brain sizes

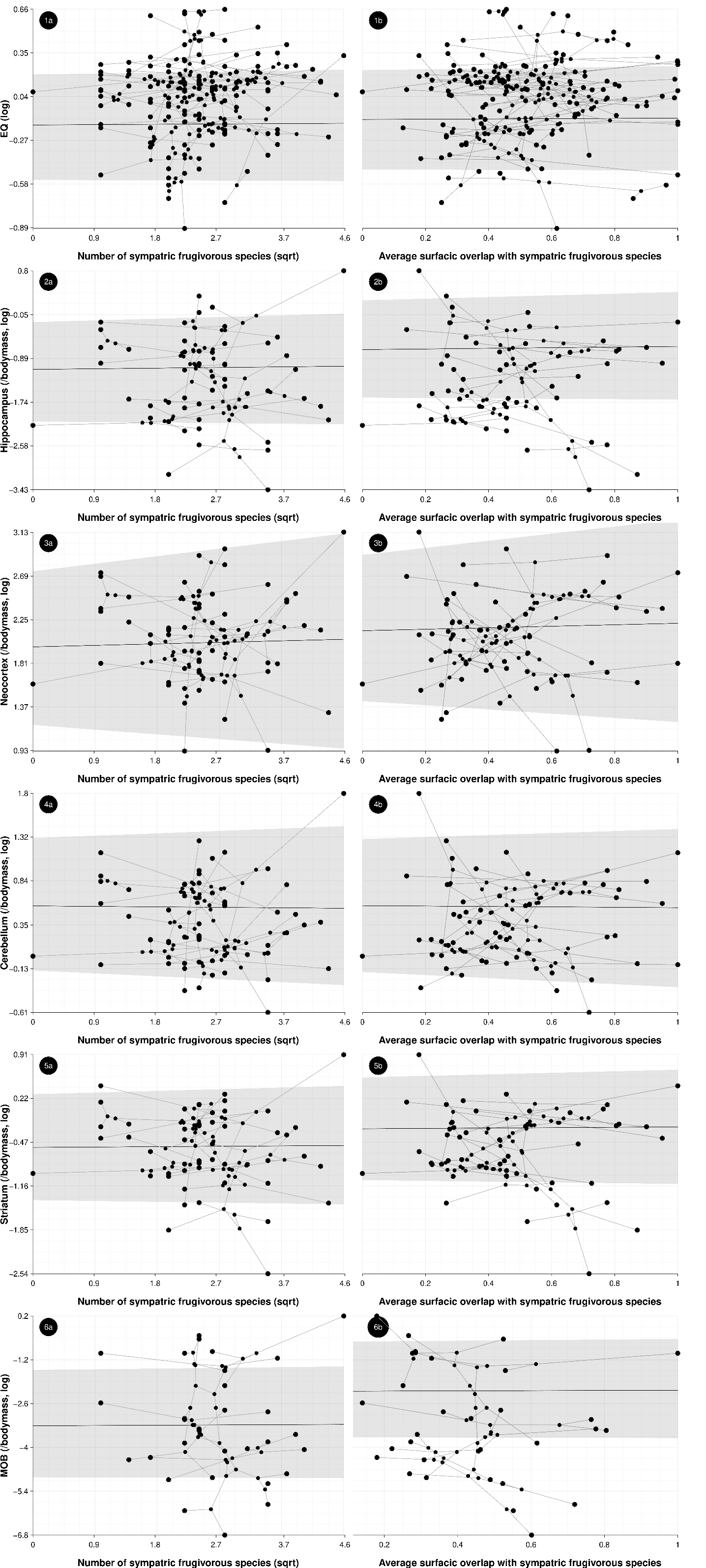

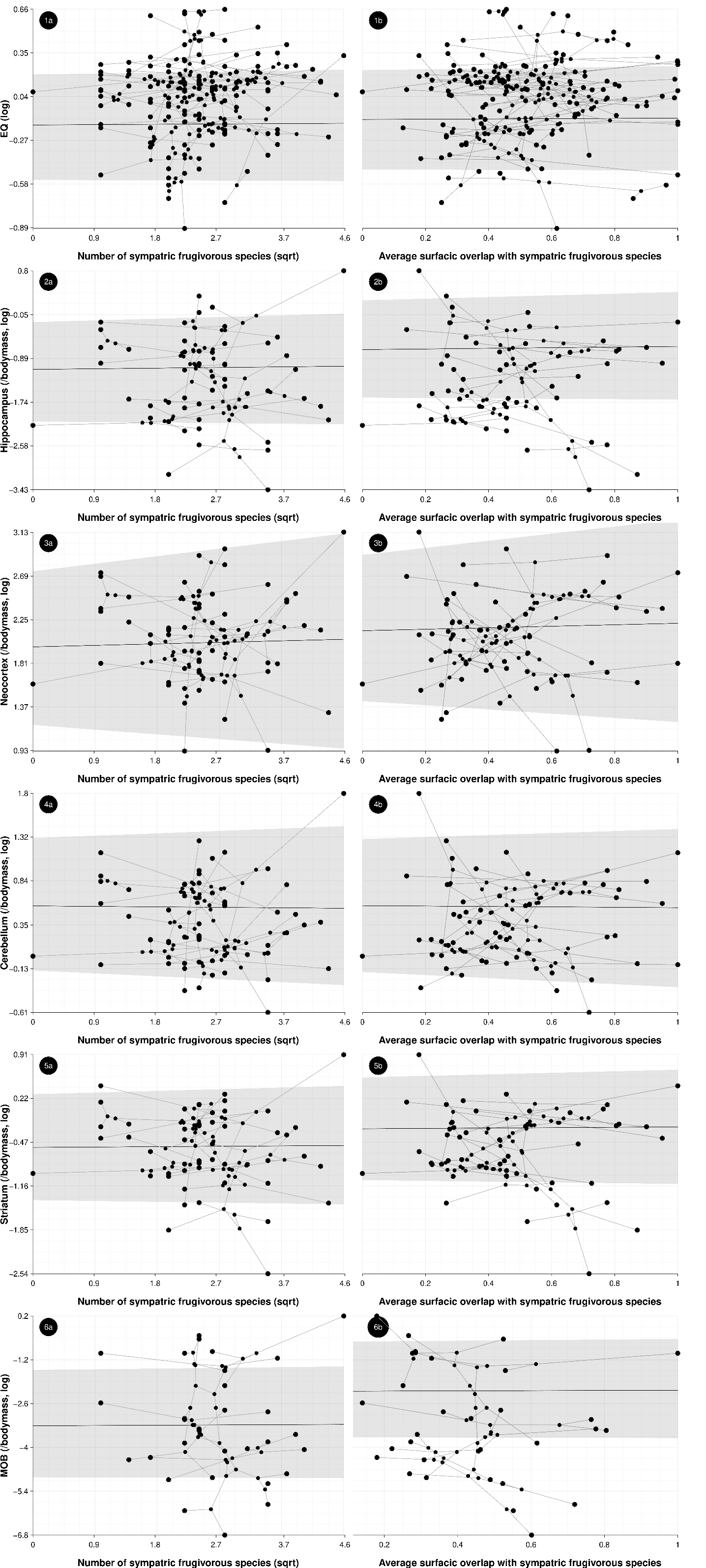

Figure S7: Phylogenetic regressions of relative brain size as a function of the indices of sympatry level| Left graphics show the correlation between the number of sympatric species on the brain size, when the effect of the average percentage of overlapping current range with sympatric frugivorous species (surfacic overlap) is averaged, while the right graphics do the opposite. Raw data are depicted with points, while the segments that link them correspond to the projected phylogenetic tree. The model fit is shown with the plain black line and the associated 95% confidence interval is depicted by the transparent grey background.

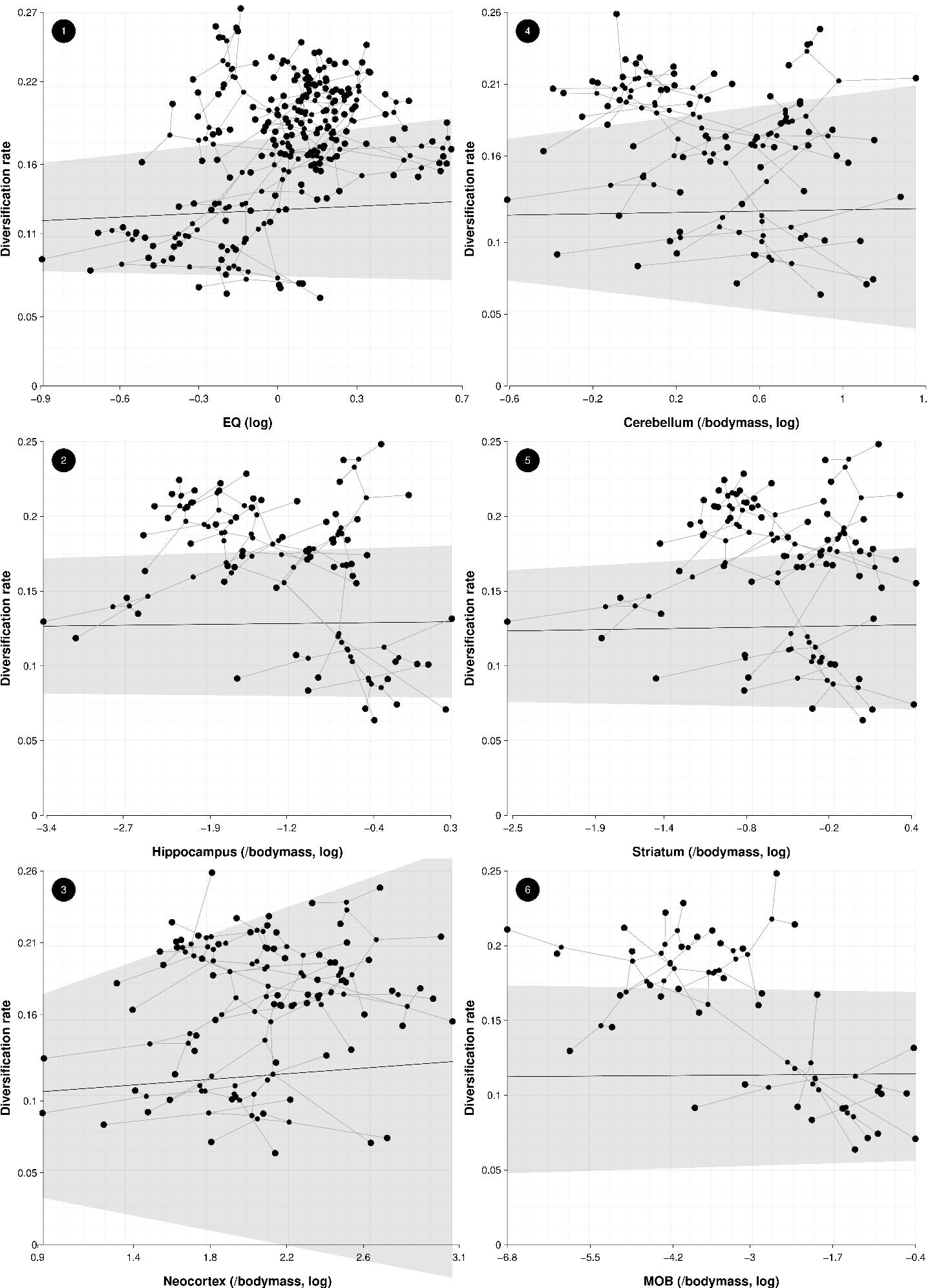
(b) Phylogenetic regressions: effect of brain size on diversification

Figure S8: Phylogenetic regressions of the net diversification rate as a function of the size of the different brain regions | Raw data are depicted with points, while the segments that link them correspond to the projected phylogenetic tree. The model fit is shown with the plain black line and the associated 95% highest density posterior is depicted by the transparent grey background.

(c) Phylogenetic regressions: effect of sympatry on diversification

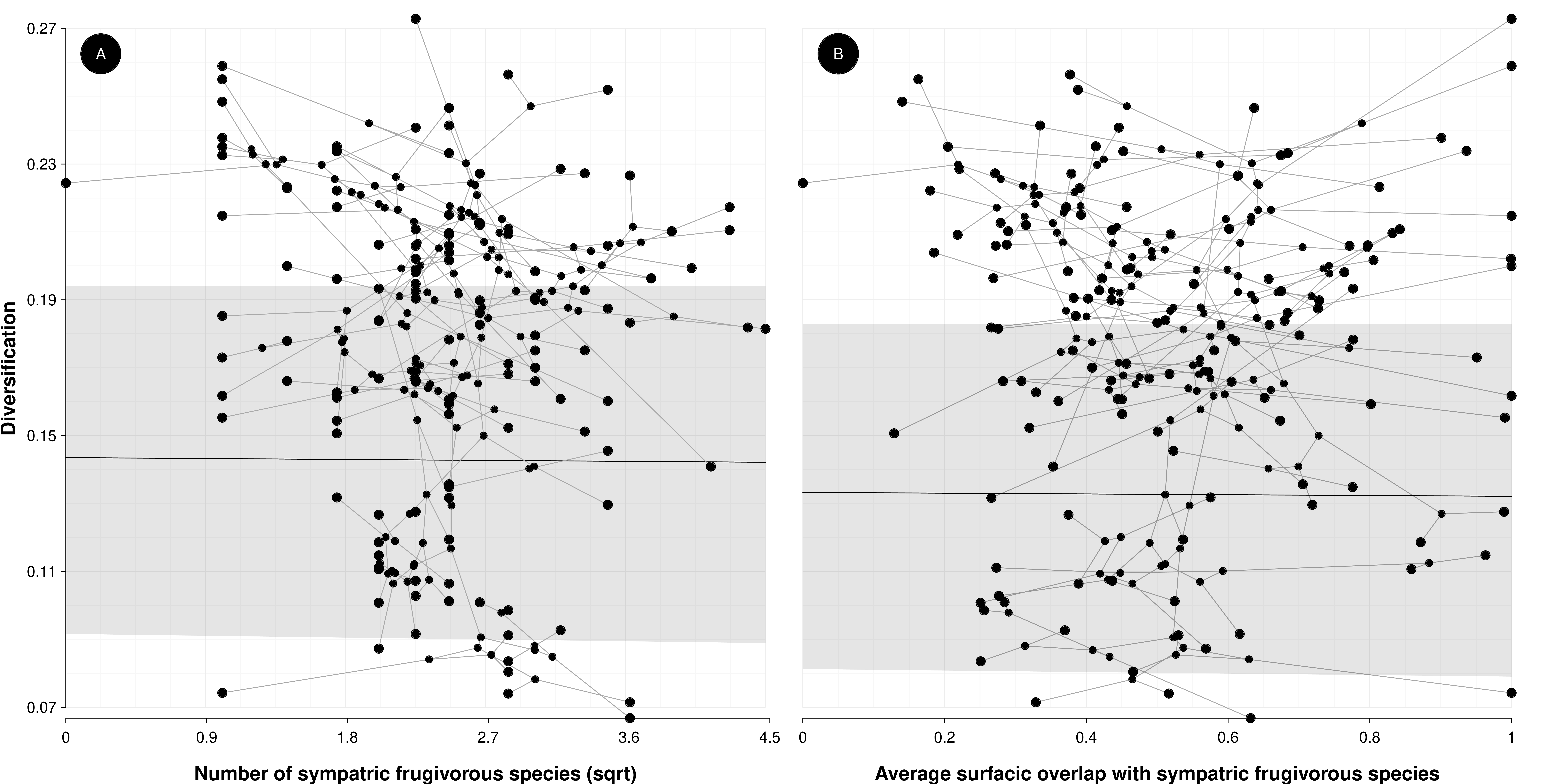

Figure S9: Phylogenetic regressions of the net diversification rate as a function of the indices of sympatry level | The left graphic depicts the correlation between the number of sympatric species and the net diversification rates, when the effect of the average percentage of overlapping current range with sympatric frugivorous species (surfacic overlap) is averaged, while the right graphic does the opposite. Raw data are depicted with points, while the segments that link them correspond to the projected phylogenetic tree. The model fit is shown with the plain black line and the associated 95% confidence interval is depicted by the transparent grey background.

(d) Forest plot of estimates

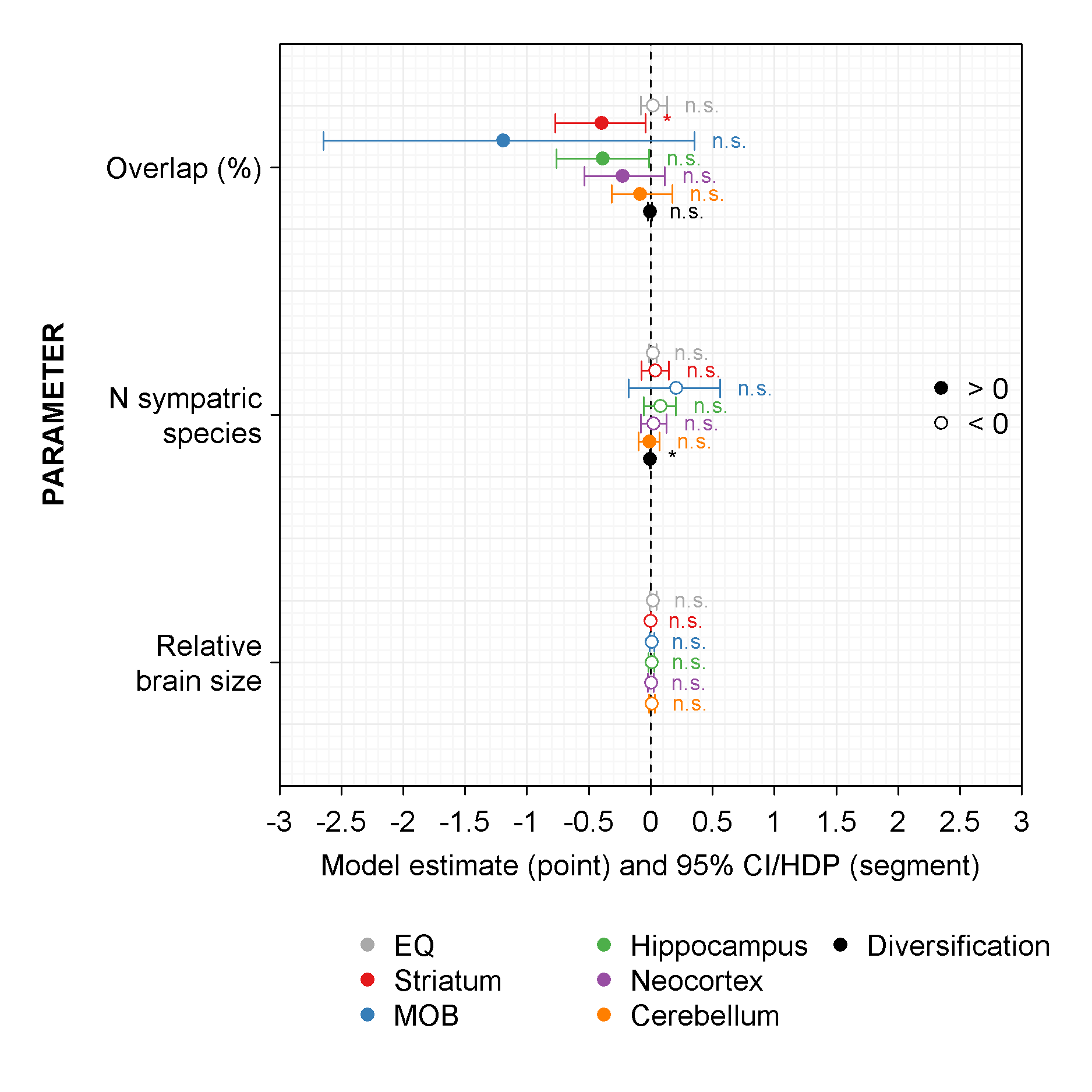

Figure S10: Forest plot of the phylogenetic regressions (when relative brain size is predicted by overlap with other species’ range and number (N) of sympatric species, when diversification is predicted by relative brain size) | CI: 95% Confidence Interval, HDP: Highest Posterior Density (when brain size is the predictor, they are barely visible because reduced). Plain dots depict negative effects, open dots depict positive effects.

#### Does sympatry shape body mass evolutionary history?

By affecting resource quantity and accessibility, it is possible that sympatry also shapes body mass. We replicated analyses on relative brain size using body mass as the variable of interest. We found that body mass was non-linearly sensitive to sympatry (evolutionary models: Supplementary Figure S6; PGLS: Supplementary Table S2).

Table S2: Model estimates and significance of phylogenetic regressions to assess the selection gradient direction | Est.=Estimate, CI2.5%=Lower border of the CI95%, CI97.5%=Upper border of the CI95%, Se= Standard deviation, t= Statistics t-value. The brain regions (as well as the associated sample size) are indicated prior to each list of estimates. the transformation (logarithm or square-root) if indicated in parentheses by the abbreviation (log or sqrt).

|  | **Est.** | **CI2.5%** | **CI97.5%** | **Se** | **t** | **p-value** |
| --- | --- | --- | --- | --- | --- | --- |
| Body mass (log) (N=130) |  |  |  |  |  |  |
| Intercept | 7.23 | 6.00 | 8.36 | 0.61 | - | - |
| % of overlapped home range | 0.03 | -0.21 | 0.26 | 0.12 | 0.26 | 0.79 |
| Number of sympatric frugivorous (sqrt) | 0.02 | -0.05 | 0.10 | 0.04 | 0.59 | 0.56 |
| Lambda | 1.00 | 0.99 | 1.00 |  |  |  |

#### Results robustness

##### Model stability

(a.1) Phylogenetic regressions: effect of sympatry on brain sizes (weighting by body mass)

Table S3: Sensitivity analysis of phylogenetic regressions to assess the relationship between relative brain sizes and species sympatry | Depicted is the minimum and maximum of estimates when one observation was removed at a time (DfBetas) or when varying the used phylogenetic tree and the data sampling (Phylogeny/Data).

| **Regression** | | **DfBetas** | | | **Phylogeny/Data** | | |
| --- | --- | --- | --- | --- | --- | --- | --- |
| **Trait** | **Variable** | **Est. min.** | **Est.** | **Est. max.** | **Est. min.** | **Est.** | **Est. max.** |
| Cerebellum (/body mass, log) | Intercept | 0.55 | 0.60 | 0.67 | 0.22 | 0.60 | 0.7 |
|  | % of overlapping range | -0.16 | -0.08 | -0.03 | -0.49 | -0.08 | -4.84e-03 |
|  | Number of sympatric frugivores | -0.04 | -0.01 | 5.27e-03 | 0.01 | -0.01 | 0.17 |
|  | Lambda | 1 | 1.00 | 1 | 0.3 | 1.00 | 1 |
| EQ (log) | Intercept | -0.19 | -0.17 | -0.13 | -0.41 | -0.17 | -0.05 |
|  | % of overlapping range | 7.14e-04 | 0.02 | 0.06 | -0.38 | 0.02 | 0.09 |
|  | Number of sympatric frugivores | 5.37e-03 | 0.02 | 0.02 | 2.7e-03 | 0.02 | 0.1 |
|  | Lambda | 0.98 | 0.98 | 0.99 | 0.35 | 0.98 | 1 |
| Hippocampus (/body mass, log) | Intercept | -1.03 | -0.92 | -0.82 | -1.16 | -0.92 | -0.41 |
|  | % of overlapping range | -0.5 | -0.39 | -0.2 | -1.37 | -0.39 | -0.28 |
|  | Number of sympatric frugivores | 0.04 | 0.08 | 0.1 | 0.03 | 0.08 | 0.21 |
|  | Lambda | 0.99 | 0.99 | 1 | 0.79 | 0.99 | 1 |
| MOB (/body mass, log) | Intercept | -3.23 | -2.76 | -2.62 | -2.99 | -2.76 | -2.55 |
|  | % of overlapping range | -1.83 | -1.20 | -0.8 | -1.5 | -1.20 | -0.87 |
|  | Number of sympatric frugivores | 0.11 | 0.21 | 0.33 | 0.13 | 0.21 | 0.26 |
|  | Lambda | 1 | 1.00 | 1 | 1 | 1.00 | 1 |
| Neocortex (/body mass, log) | Intercept | 1.95 | 2.07 | 2.23 | 1.73 | 2.07 | 2.32 |
|  | % of overlapping range | -0.31 | -0.23 | -0.03 | -0.55 | -0.23 | -0.02 |
|  | Number of sympatric frugivores | -0.02 | 0.02 | 0.06 | -0.04 | 0.02 | 0.15 |
|  | Lambda | 0.98 | 0.99 | 1 | 0.23 | 0.99 | 1 |
| Striatum (/body mass, log) | Intercept | -0.45 | -0.36 | -0.26 | -0.76 | -0.36 | -0.07 |
|  | % of overlapping range | -0.46 | -0.40 | -0.28 | -0.92 | -0.40 | -0.22 |
|  | Number of sympatric frugivores | 4.03e-03 | 0.03 | 0.06 | 0.01 | 0.03 | 0.18 |
|  | Lambda | 0.98 | 0.98 | 1 | 0.79 | 0.98 | 1 |

(a.2) Phylogenetic regressions: effect of sympatry on brain sizes (weighting by brain size)

Table S4: Sensitivity analysis of phylogenetic regressions to assess the relationship between relative brain sizes and species sympatry | Depicted is the minimum and maximum of estimates when one observation was removed at a time (DfBetas) or when varying the used phylogenetic tree and the data sampling (Phylogeny/Data).

| **Regression** | | **DfBetas** | | | **Phylogeny/Data** | | |
| --- | --- | --- | --- | --- | --- | --- | --- |
| **Trait** | **Variable** | **Est. min.** | **Est.** | **Est. max.** | **Est. min.** | **Est.** | **Est. max.** |
| Cerebellum (/whole-brain size, log) | Intercept | -2.2 | -2.18 | -2.14 | 0.09 | -2.18 | 0.73 |
|  | % of overlapping range | -6.68e-03 | 0.06 | 0.08 | -0.68 | 0.06 | -0.07 |
|  | Number of sympatric frugivores | 8.87e-03 | 0.01 | 0.02 | 0.02 | 0.01 | 0.2 |
|  | Lambda | 0.79 | 0.80 | 0.86 | 0.31 | 0.80 | 1 |
| Hippocampus (/whole-brain size, log) | Intercept | -3.58 | -3.52 | -3.42 | -1.27 | -3.52 | -0.44 |
|  | % of overlapping range | -0.42 | -0.29 | -0.2 | -1.51 | -0.29 | -0.43 |
|  | Number of sympatric frugivores | 8.99e-03 | 0.05 | 0.06 | 0.02 | 0.05 | 0.26 |
|  | Lambda | 0.92 | 0.93 | 0.98 | 0.84 | 0.93 | 1 |
| MOB (/whole-brain size, log) | Intercept | -5.65 | -5.34 | -5.23 | -2.92 | -5.34 | -2.56 |
|  | % of overlapping range | -1.12 | -0.84 | -0.58 | -1.45 | -0.84 | -0.86 |
|  | Number of sympatric frugivores | 0.07 | 0.13 | 0.19 | 0.14 | 0.13 | 0.27 |
|  | Lambda | 1 | 1.00 | 1 | 1 | 1.00 | 1 |
| Neocortex (/whole-brain size, log) | Intercept | -0.79 | -0.76 | -0.64 | 1.72 | -0.76 | 2.28 |
|  | % of overlapping range | 0.06 | 0.15 | 0.2 | -0.59 | 0.15 | 0.01 |
|  | Number of sympatric frugivores | 3.54e-03 | 0.03 | 0.04 | -0.02 | 0.03 | 0.15 |
|  | Lambda | 0.35 | 0.44 | 0.6 | 0.13 | 0.44 | 1 |
| Striatum (/whole-brain size, log) | Intercept | -3.06 | -3.03 | -2.97 | -0.67 | -3.03 | -0.05 |
|  | % of overlapping range | -0.28 | -0.24 | -0.19 | -0.97 | -0.24 | -0.41 |
|  | Number of sympatric frugivores | 3.44e-03 | 0.02 | 0.03 | -0.01 | 0.02 | 0.18 |
|  | Lambda | 0.78 | 0.82 | 0.87 | 0.75 | 0.82 | 1 |

(b) Phylogenetic regressions: diversification and brain size (weighting by body mass)

Table S5: Sensitivity analysis of phylogenetic regressions to assess the relationship between species diversification rates and relative brain sizes | Depicted is the minimum and maximum of estimates when varying the used phylogenetic tree and the data sampling (Phylogeny/Data) or when the sampling fraction varied (Sampling fraction).

| **Regression** | | **Phylogeny/Data** | | | **Sampling fraction** | | |
| --- | --- | --- | --- | --- | --- | --- | --- |
| **Model** | **Variable** | **Est. min.** | **Est.** | **Est. max.** | **Est. min.** | **Est.** | **Est. max.** |
| Cerebellum (/body mass, log) | Intercept | 0.12 | 0.12 | 0.12 | 0.11 | 0.12 | 0.13 |
|  | Trait | 7.60e-04 | 3.94e-03 | 4.17e-03 | 1.91e-04 | 3.94e-03 | 6.13e-03 |
|  | Lambda | 0.69 | 0.74 | 0.74 | 0.72 | 0.74 | 0.75 |
| EQ (log) | Intercept | 0.12 | 0.12 | 0.12 | 0.11 | 0.12 | 0.13 |
|  | Trait | 7.51e-03 | 0.02 | 0.02 | 6.04e-03 | 0.02 | 0.02 |
|  | Lambda | 0.77 | 0.83 | 0.83 | 0.8 | 0.83 | 0.85 |
| Hippocampus (/body mass, log) | Intercept | 0.13 | 0.13 | 0.13 | 0.12 | 0.13 | 0.14 |
|  | Trait | 3.73e-03 | 9.10e-03 | 9.10e-03 | 4.07e-03 | 9.10e-03 | 9.10e-03 |
|  | Lambda | 0.69 | 0.73 | 0.73 | 0.72 | 0.73 | 0.75 |
| MOB (/body mass, log) | Intercept | 0.1 | 0.11 | 0.11 | 0.1 | 0.11 | 0.12 |
|  | Trait | -9.89e-03 | -4.79e-03 | -4.76e-03 | -7.64e-03 | -4.79e-03 | -4.10e-03 |
|  | Lambda | 0.6 | 0.65 | 0.65 | 0.64 | 0.65 | 0.65 |
| Neocortex (/body mass, log) | Intercept | 0.1 | 0.1 | 0.11 | 0.1 | 0.1 | 0.12 |
|  | Trait | 4.69e-03 | 7.26e-03 | 7.26e-03 | 1.60e-03 | 7.26e-03 | 7.26e-03 |
|  | Lambda | 0.69 | 0.74 | 0.74 | 0.72 | 0.74 | 0.75 |
| Striatum (/body mass, log) | Intercept | 0.12 | 0.12 | 0.13 | 0.12 | 0.12 | 0.14 |
|  | Trait | 5.89e-03 | 9.11e-03 | 9.41e-03 | 6.27e-03 | 9.11e-03 | 9.13e-03 |
|  | Lambda | 0.69 | 0.73 | 0.73 | 0.72 | 0.73 | 0.75 |

(b) Phylogenetic regressions: diversification and sympatry

Table S6: Sensitivity analysis of phylogenetic regressions to assess the relationship between species diversification rates and sympatry | Depicted is the minimum and maximum of estimates when one observation was removed at a time (DfBetas), when varying the used phylogenetic tree and the data sampling (Phylogeny/Data), or when the sampling fraction varied (Sampling fraction)

| **Regression** | **DfBetas** | | | **Phylogeny/Data** | | | **Sampling fraction** | | |
| --- | --- | --- | --- | --- | --- | --- | --- | --- | --- |
| **Variable** | **Est. min.** | **Est.** | **Est. max.** | **Est. min.** | **Est.** | **Est. max.** | **Est. min.** | **Est.** | **Est. max.** |
| Intercept | 0.14 | 0.15 | 0.15 | 0.15 | 0.15 | 0.15 | 0.14 | 0.15 | 0.17 |
| % of overlapping range | -8.58e-03 | -5.40e-03 | 5.85e-05 | -5.73e-03 | -5.40e-03 | -5.73e-03 | -8.6e-03 | -5.40e-03 | 7.71e-03 |
| Number of sympatric frugivores | -6.03e-03 | -5.04e-03 | -4.24e-03 | -5.2e-03 | -5.04e-03 | -5.2e-03 | -9.06e-03 | -5.04e-03 | -6.14e-03 |
| Lambda | 0.96 | 0.96 | 0.97 | 0.96 | 0.96 | 0.96 | 0.94 | 0.96 | 0.99 |

(d) Phylogenetic regressions: effect of sympatry on body mass

Table S7: Sensitivity analysis of phylogenetic regressions to assess the relationship between body mass and species sympatry | Depicted is the minimum and maximum of estimates when one observation was removed at a time (DfBetas) or when varying the used phylogenetic tree and the data sampling (Phylogeny/Data).

| **Regression** | | **DfBetas** | | | **Phylogeny/Data** | | |
| --- | --- | --- | --- | --- | --- | --- | --- |
| **Trait** | **Variable** | **Est. min.** | **Est.** | **Est. max.** | **Est. min.** | **Est.** | **Est. max.** |
| Body mass (log) | Intercept | 7.17 | 7.23 | 7.28 | 7.22 | 7.23 | 7.22 |
|  | % of overlapping range | -0.02 | 0.03 | 0.08 | 0.04 | 0.03 | 0.04 |
|  | Number of sympatric frugivores | 9.91e-03 | 0.02 | 0.03 | 0.03 | 0.02 | 0.03 |
|  | Lambda | 1 | 1.00 | 1 | 1 | 1.00 | 1 |

##### Model assumptions

We present below the visual assessment of linear modelling assumptions (non-Bayesian regression: histogram of residuals, Q-Q plot, and scatterplot of fitted values vs residuals; Bayesian regression: trace of trait and estimated density distribution).

(a.1) Phylogenetic regressions: effect of sympatry on brain sizes (weighting by body mass)

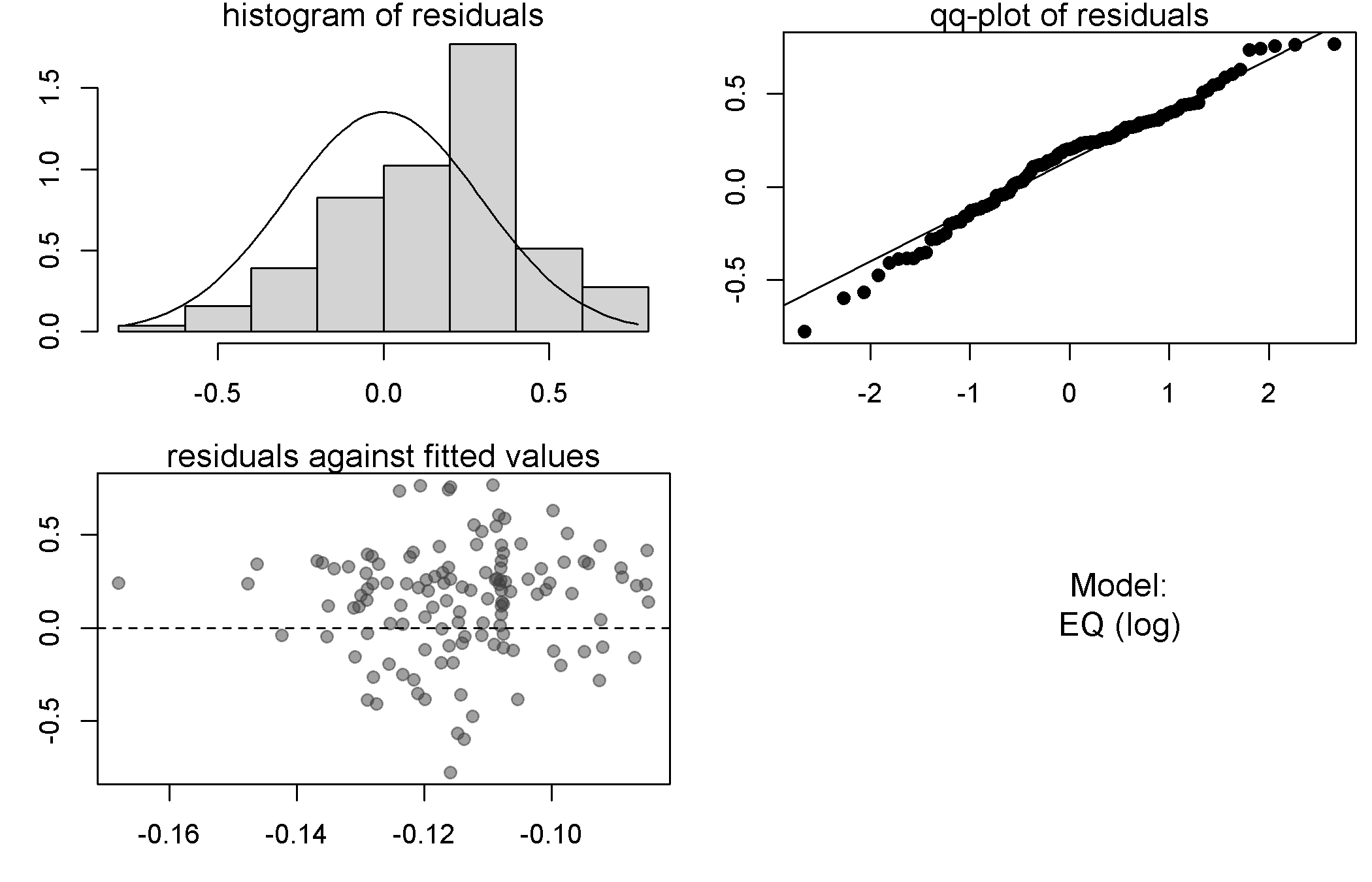

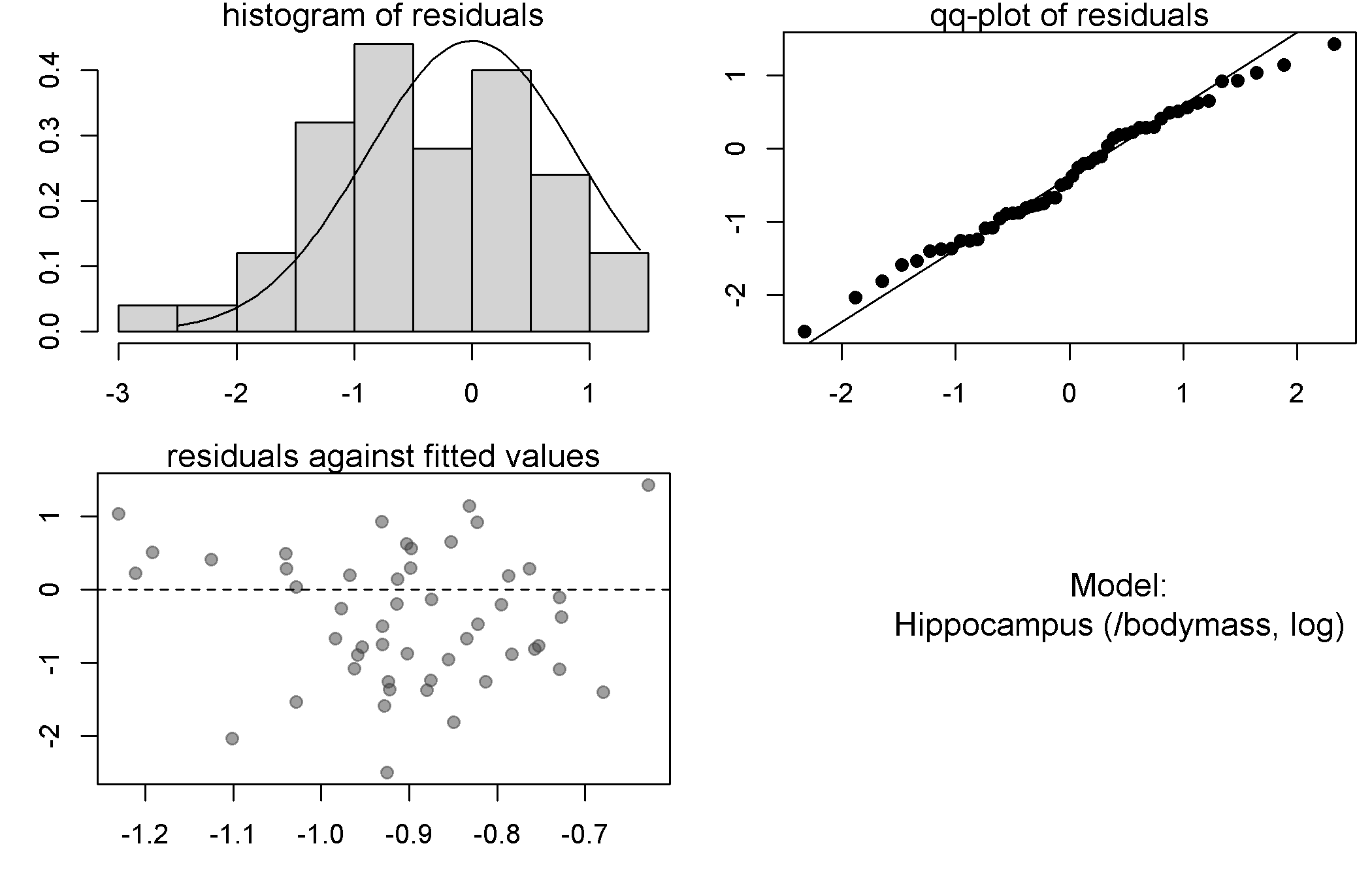

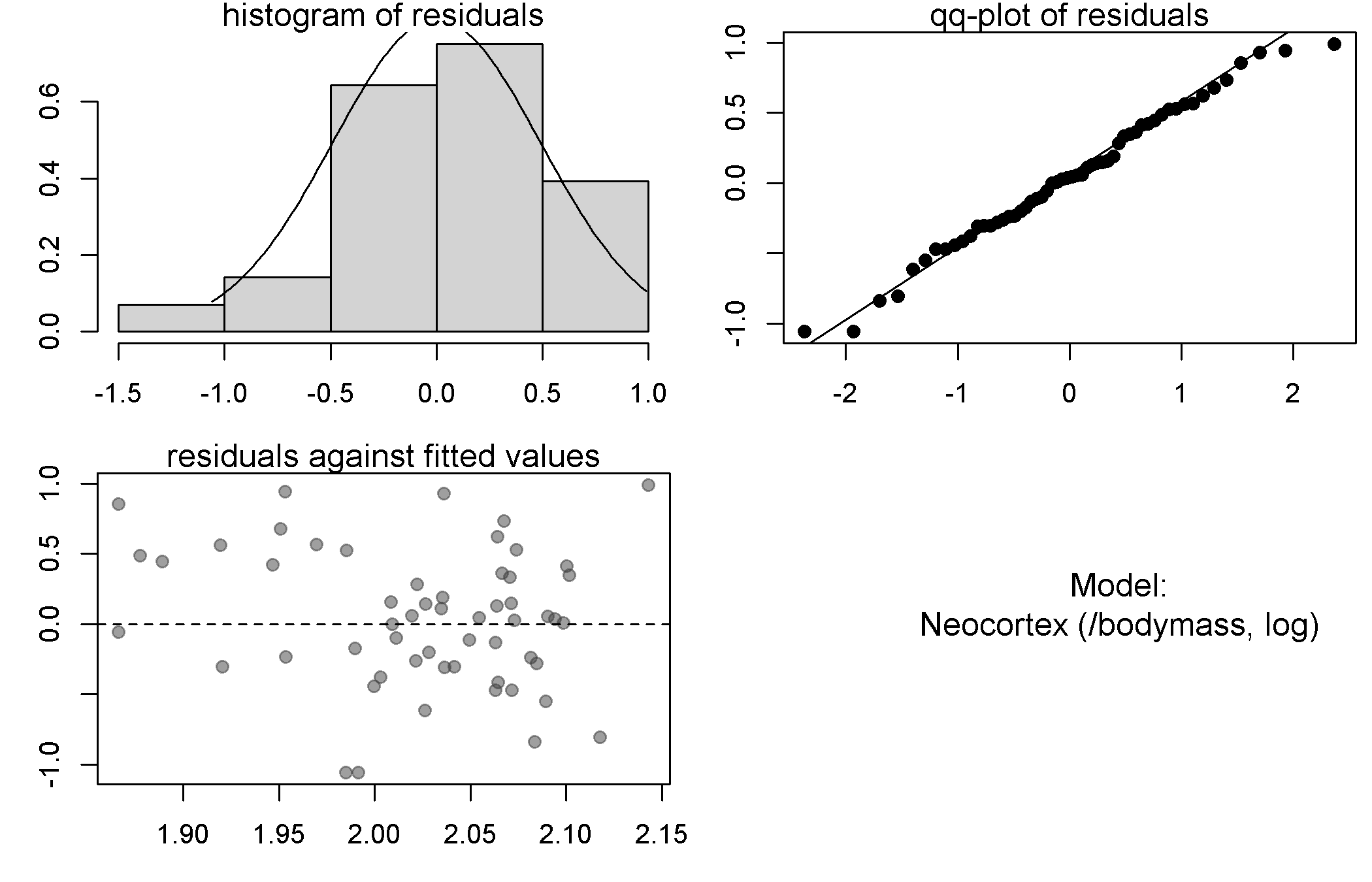

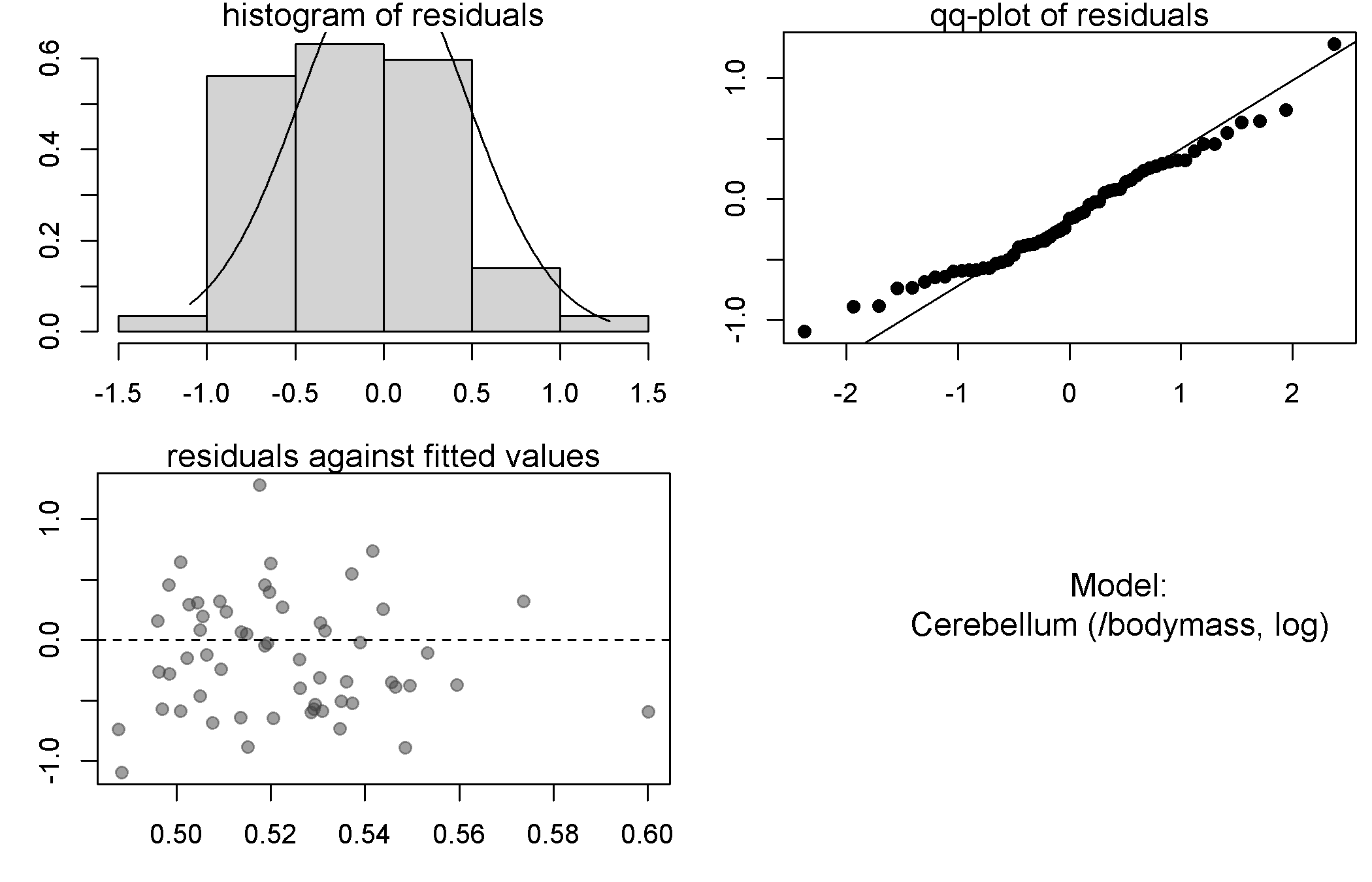

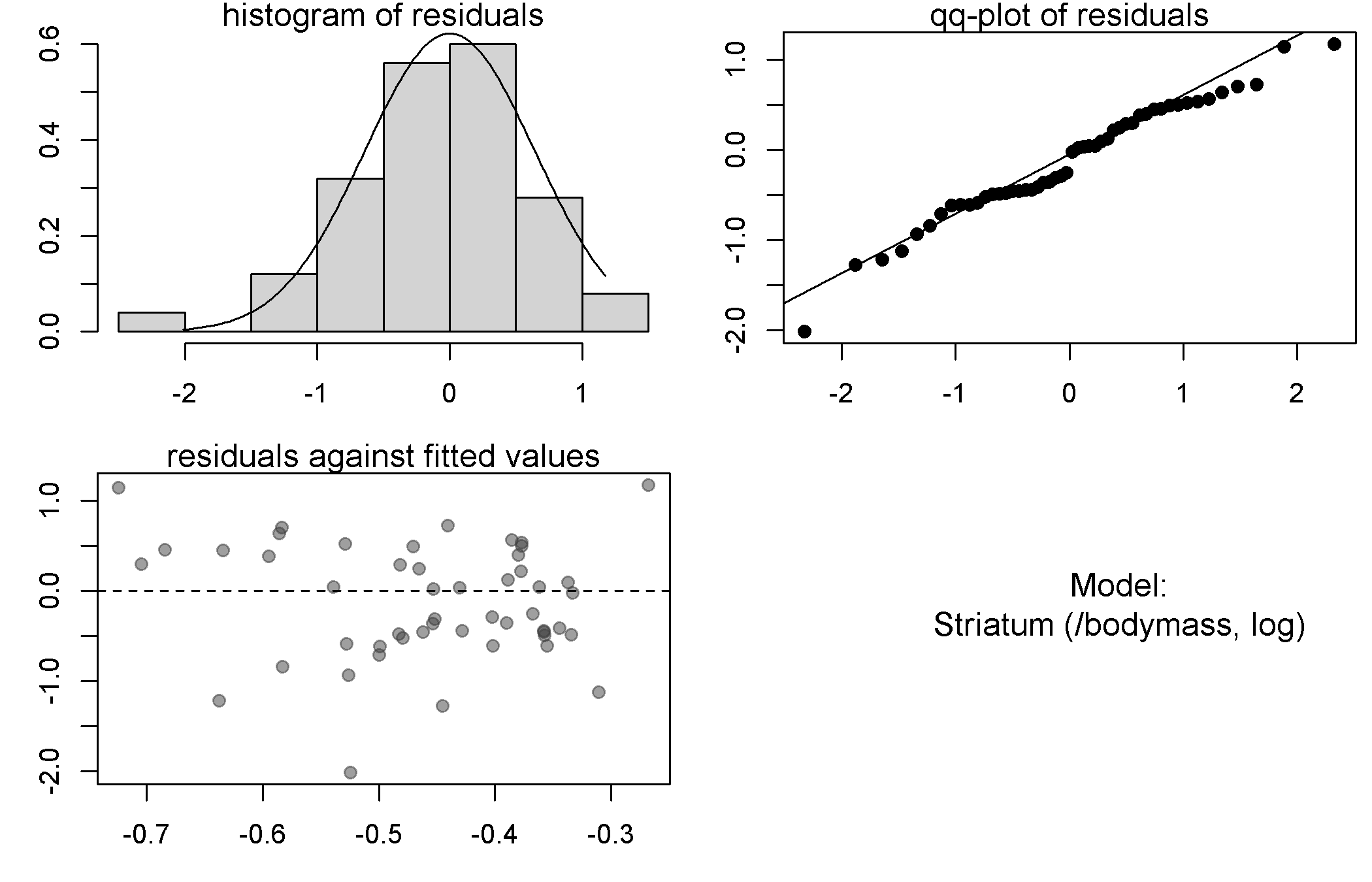

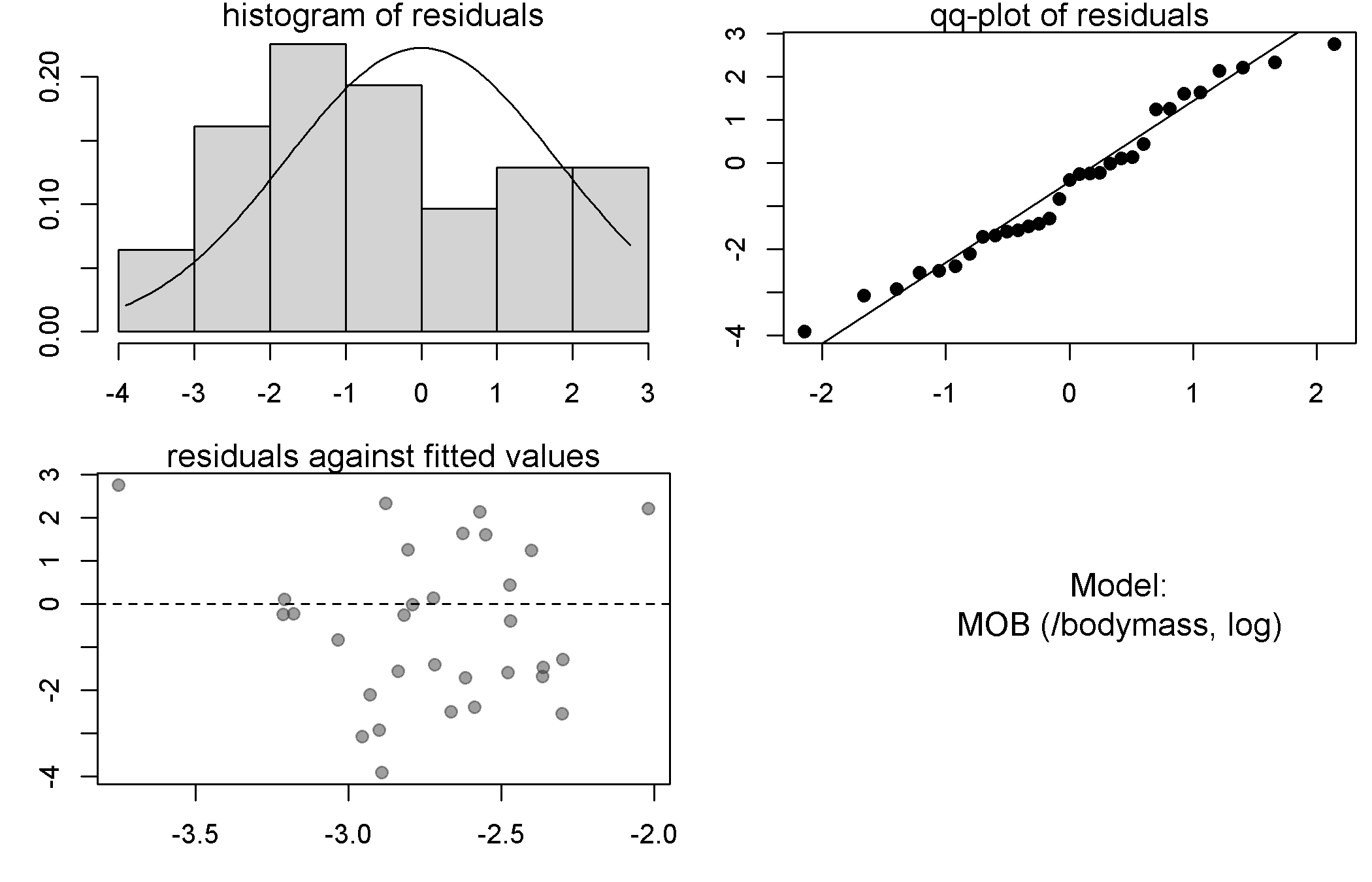

(a.2) Phylogenetic regressions: effect of sympatry on brain sizes (weighting by brain size)

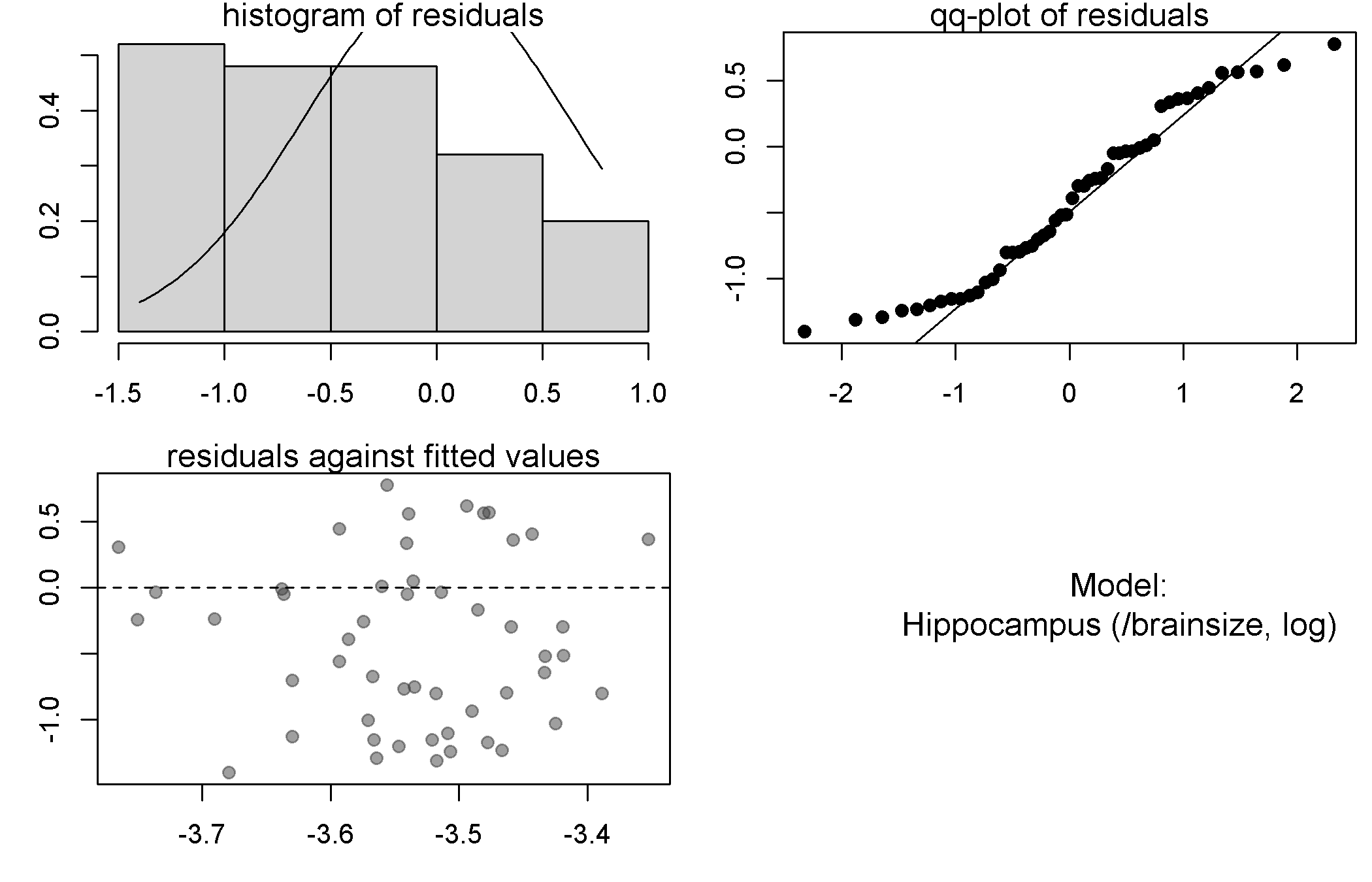

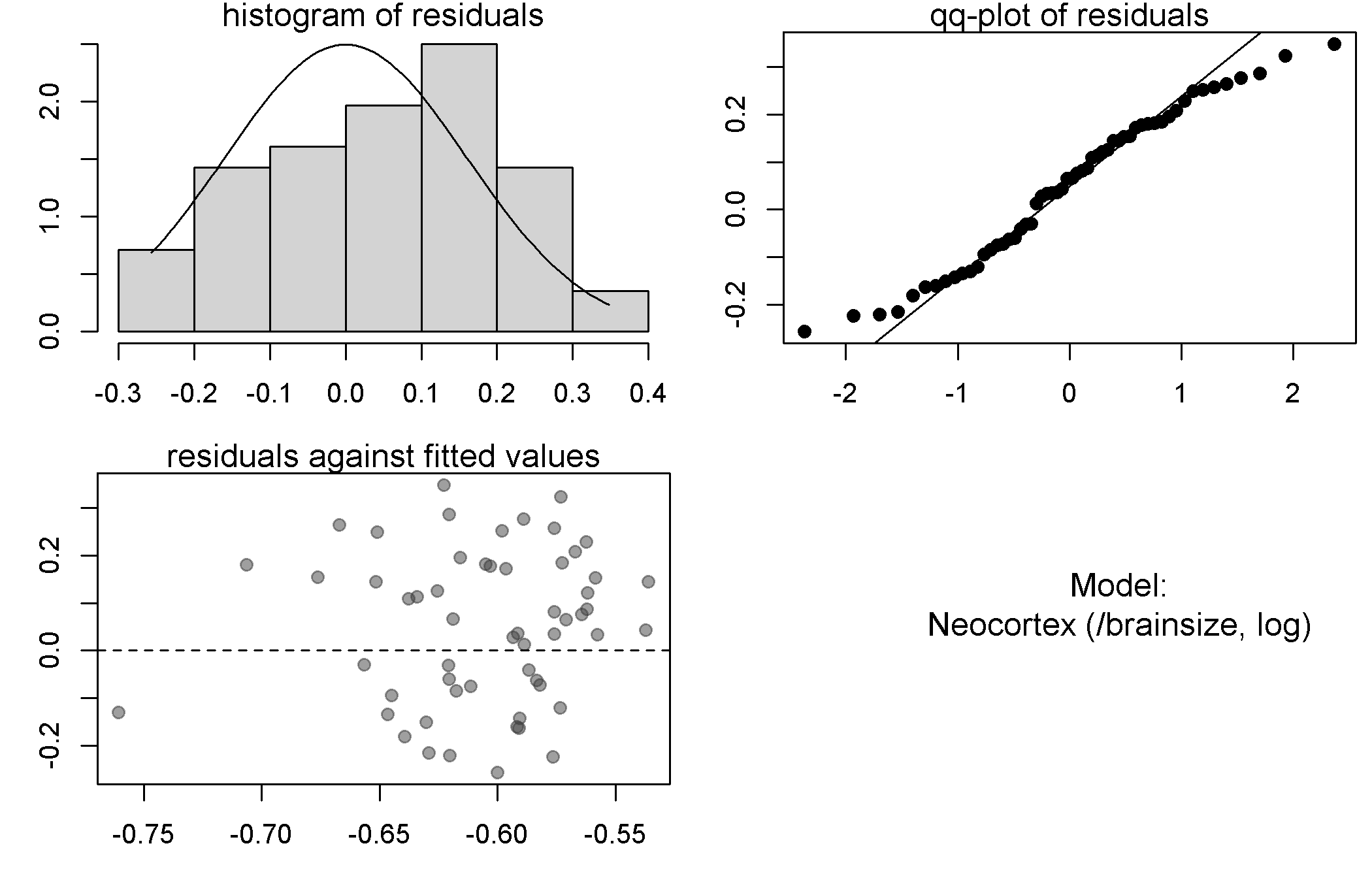

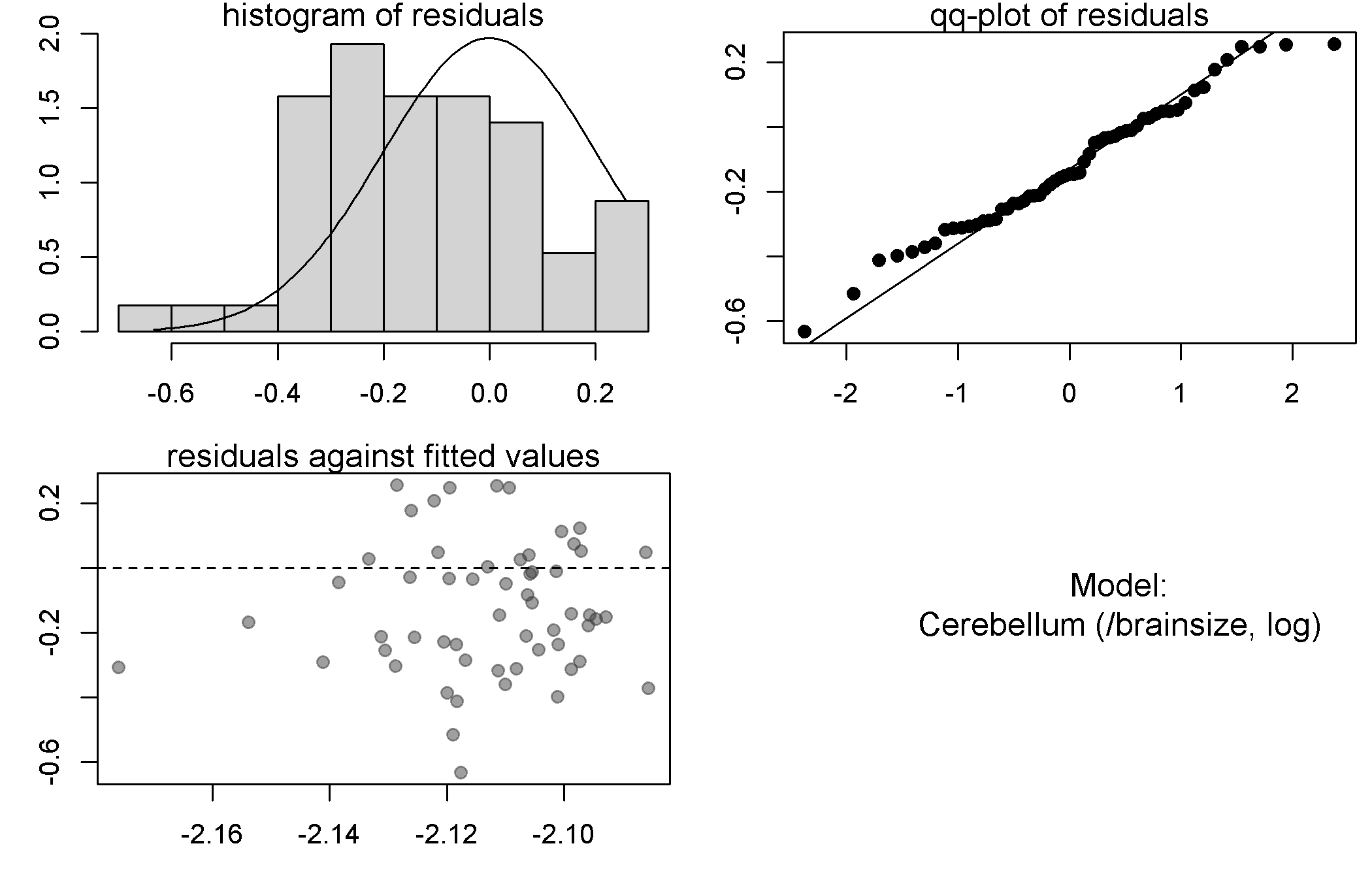

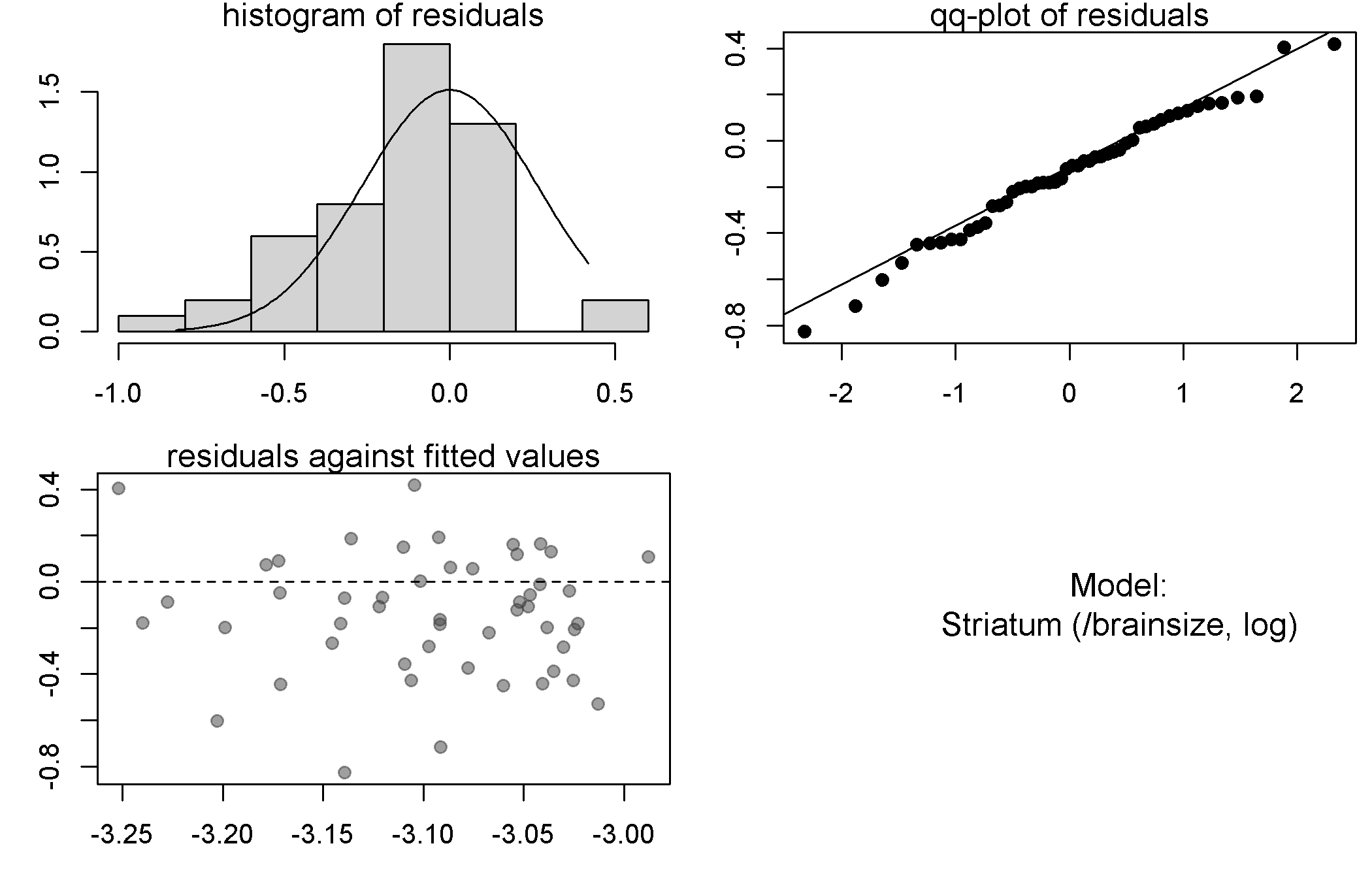

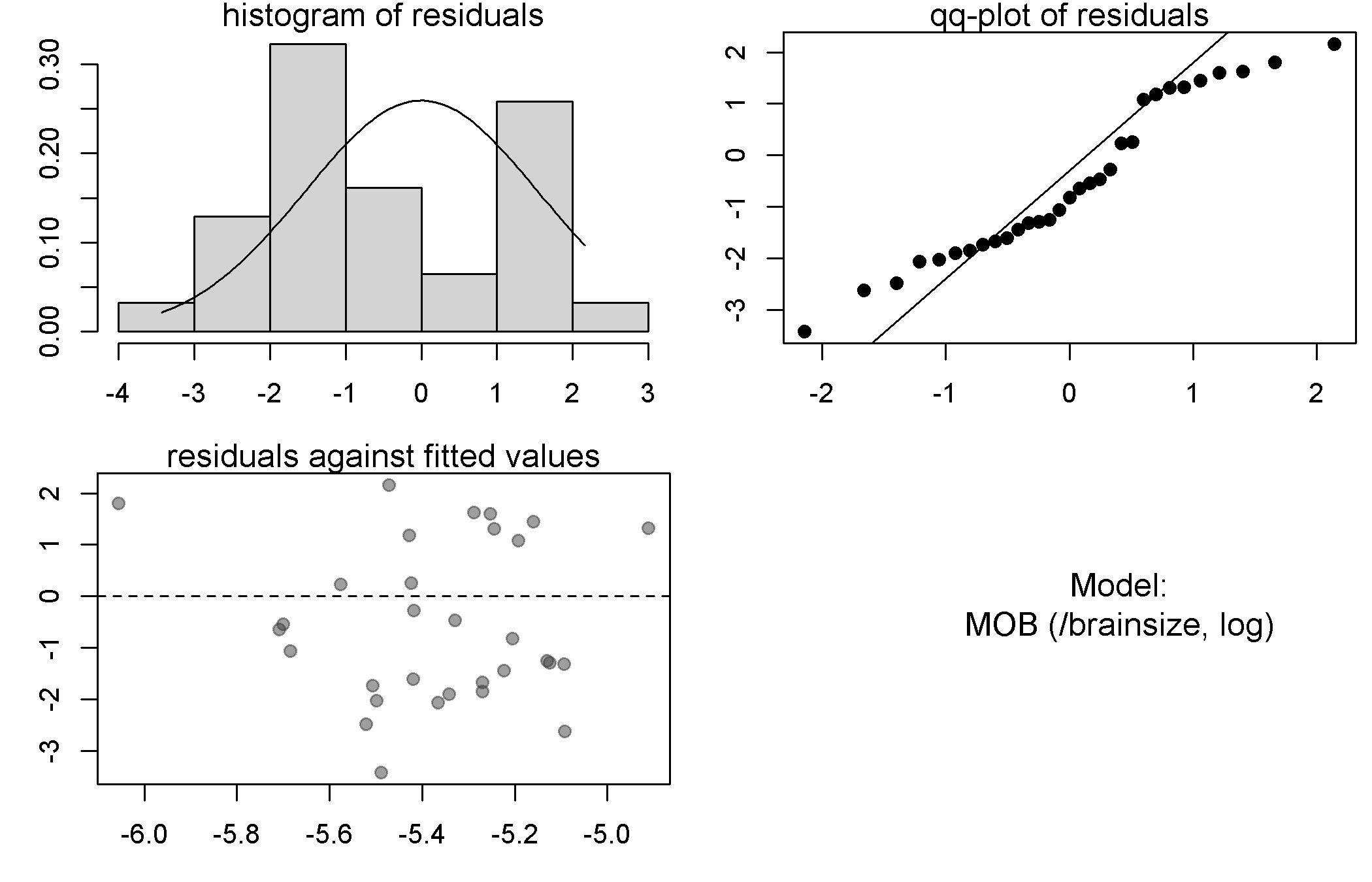

(b) Phylogenetic regressions: diversification and brain size

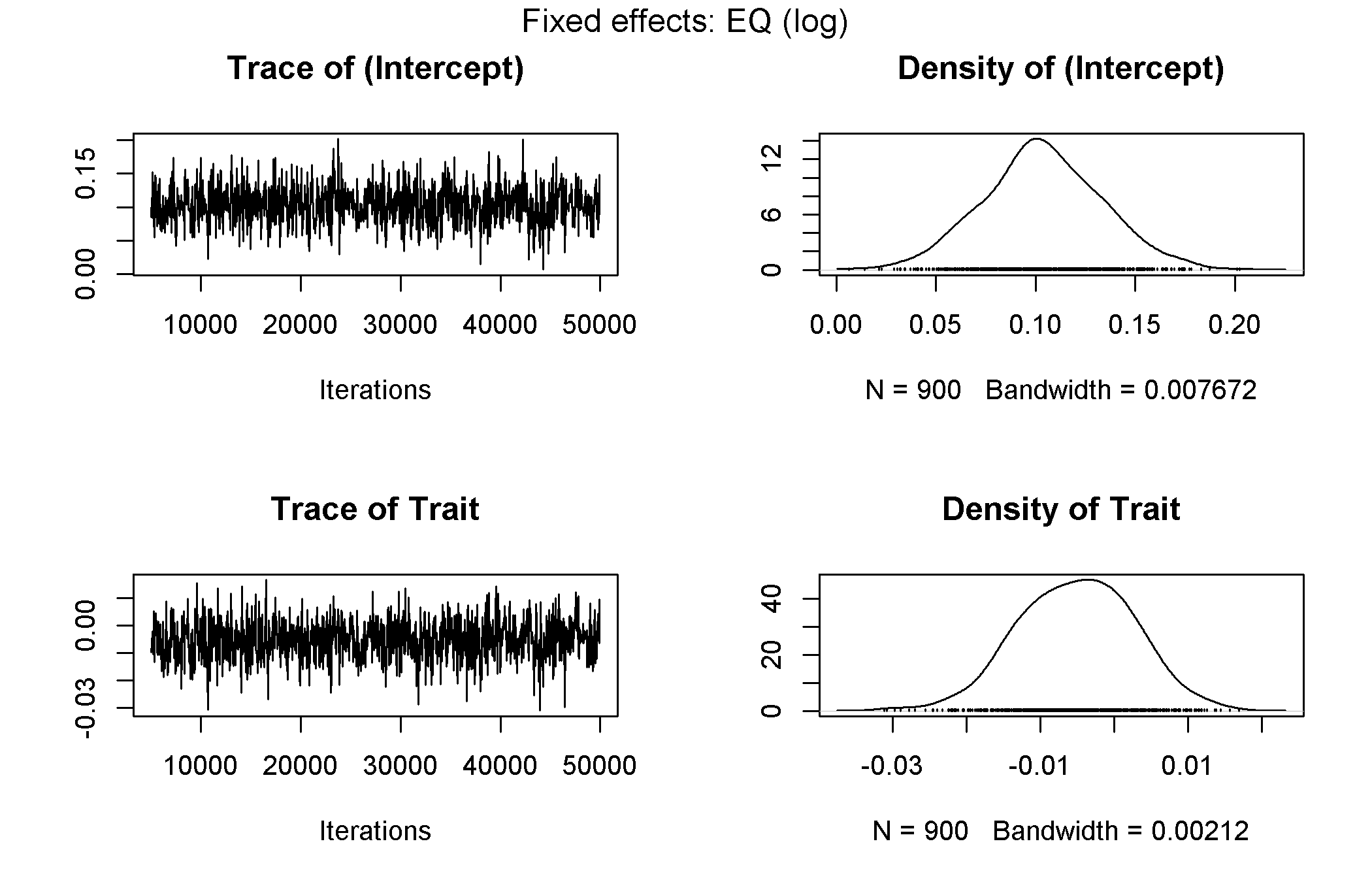

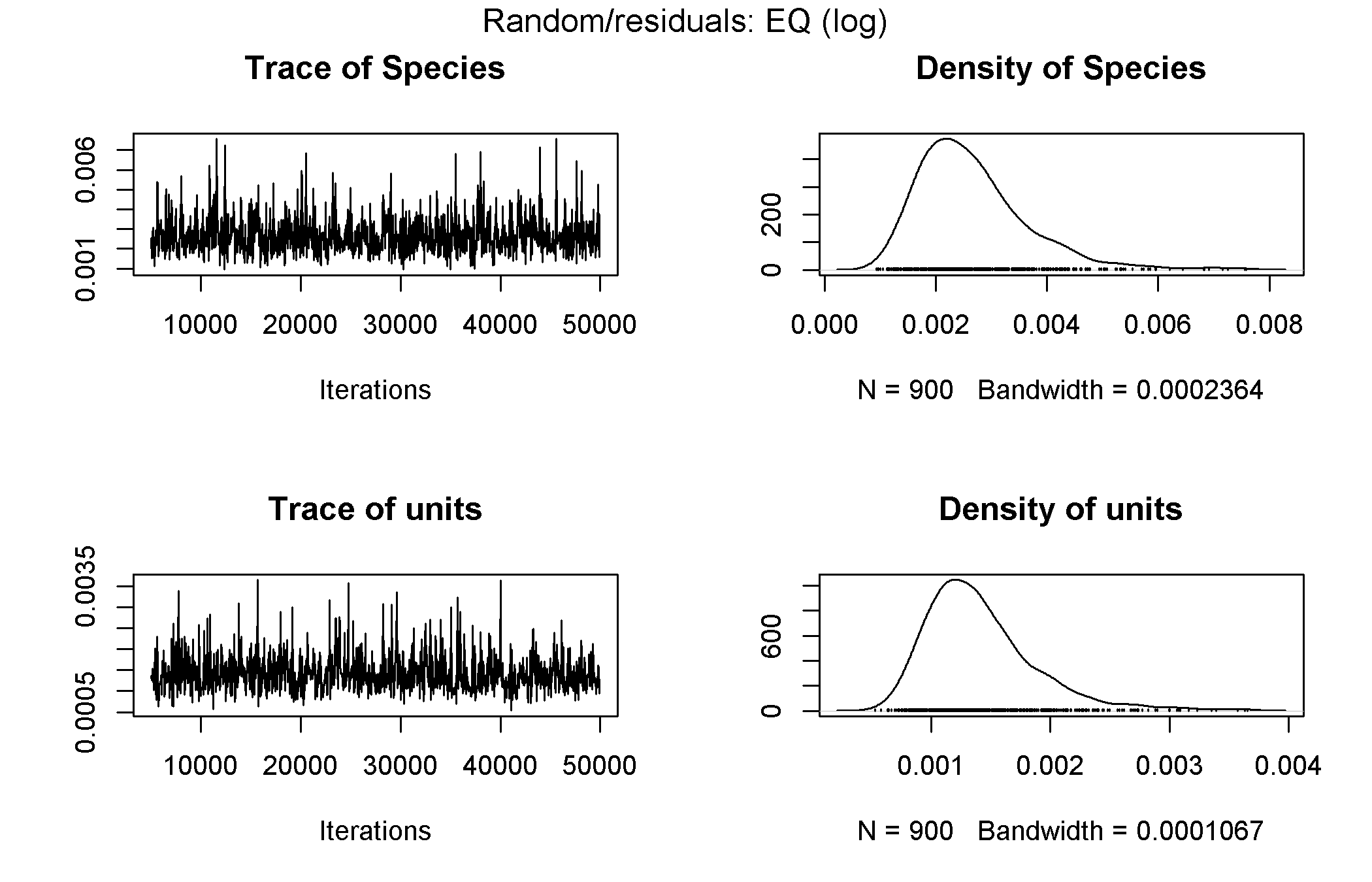

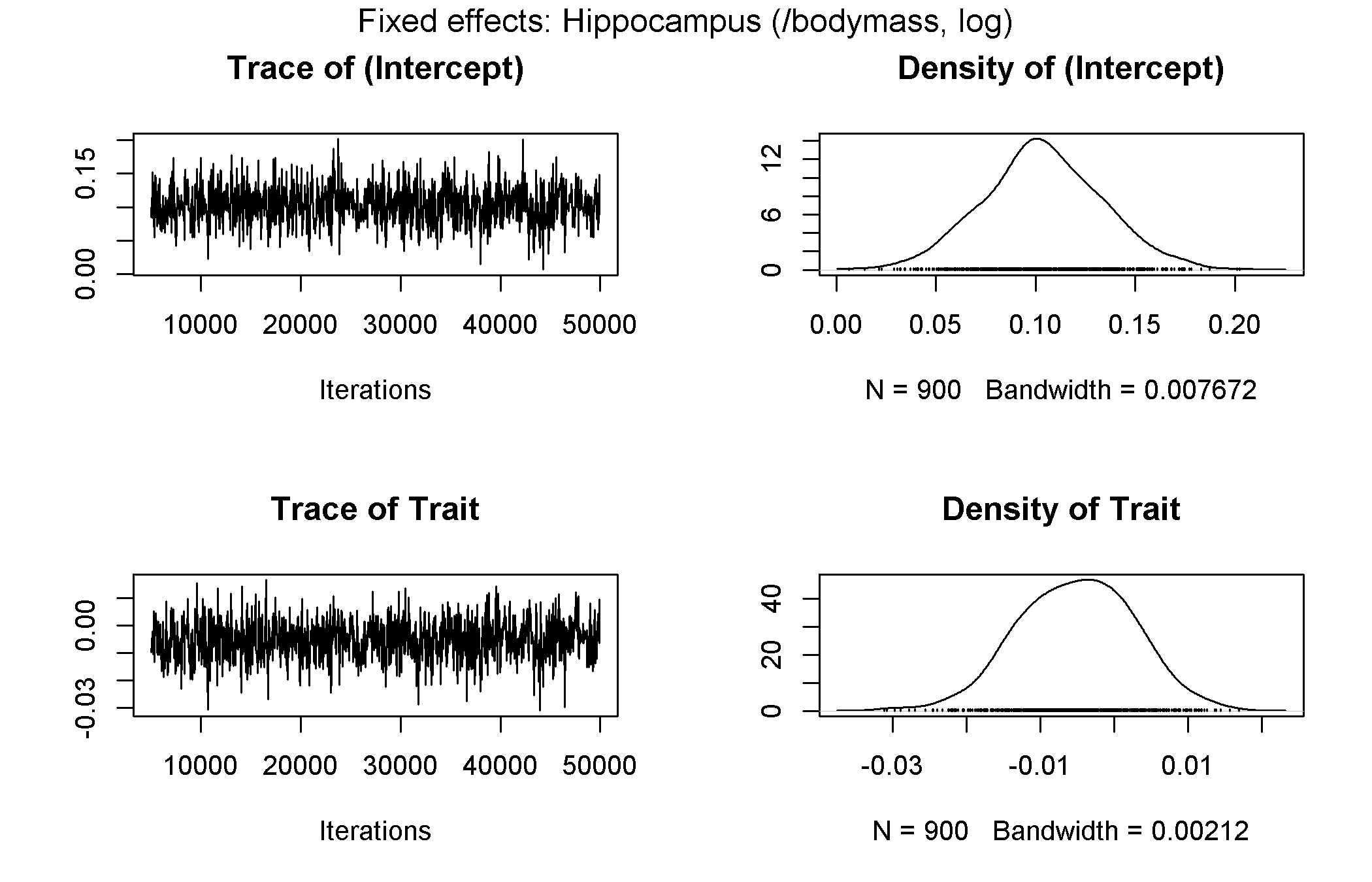

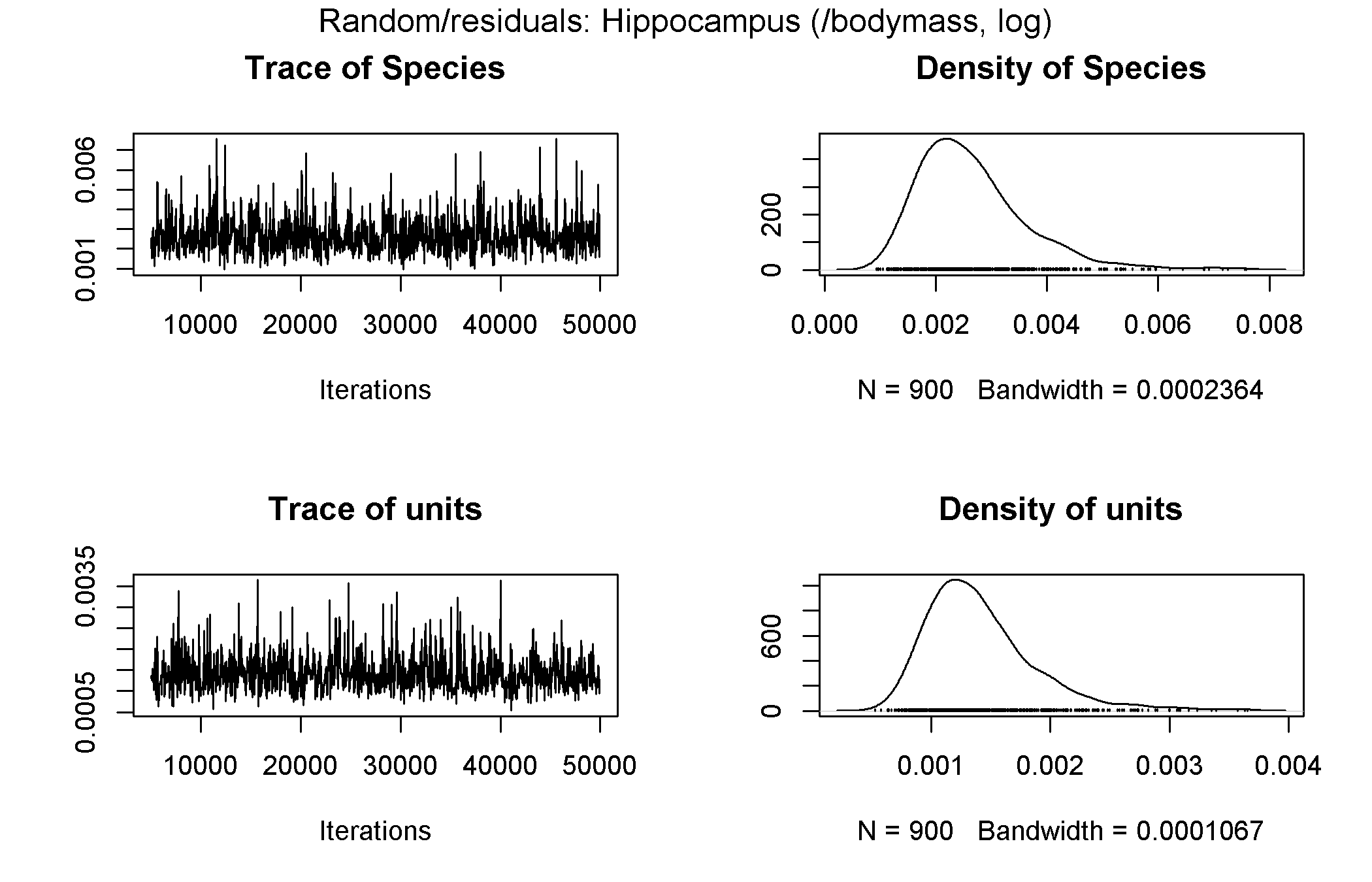

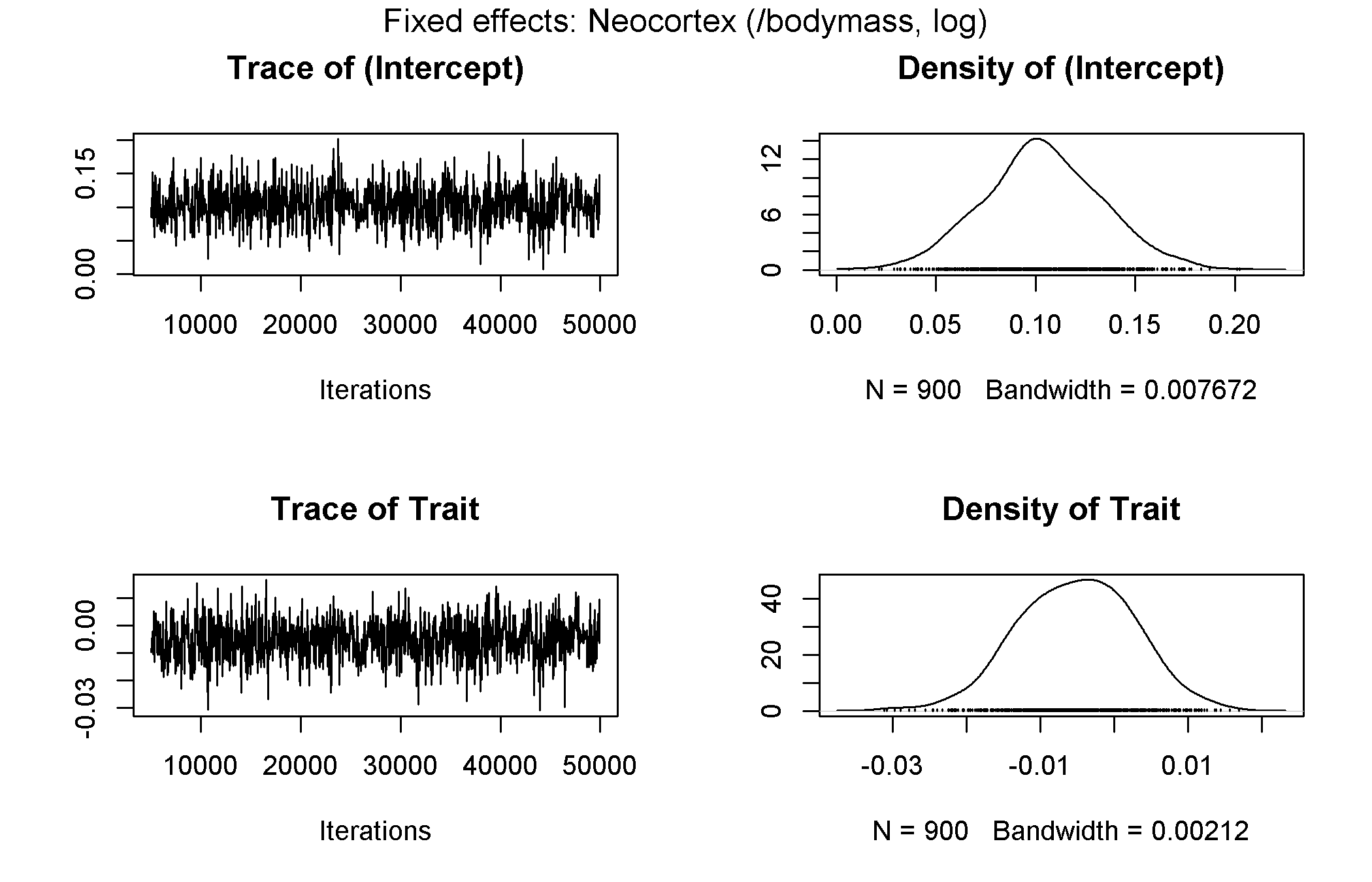

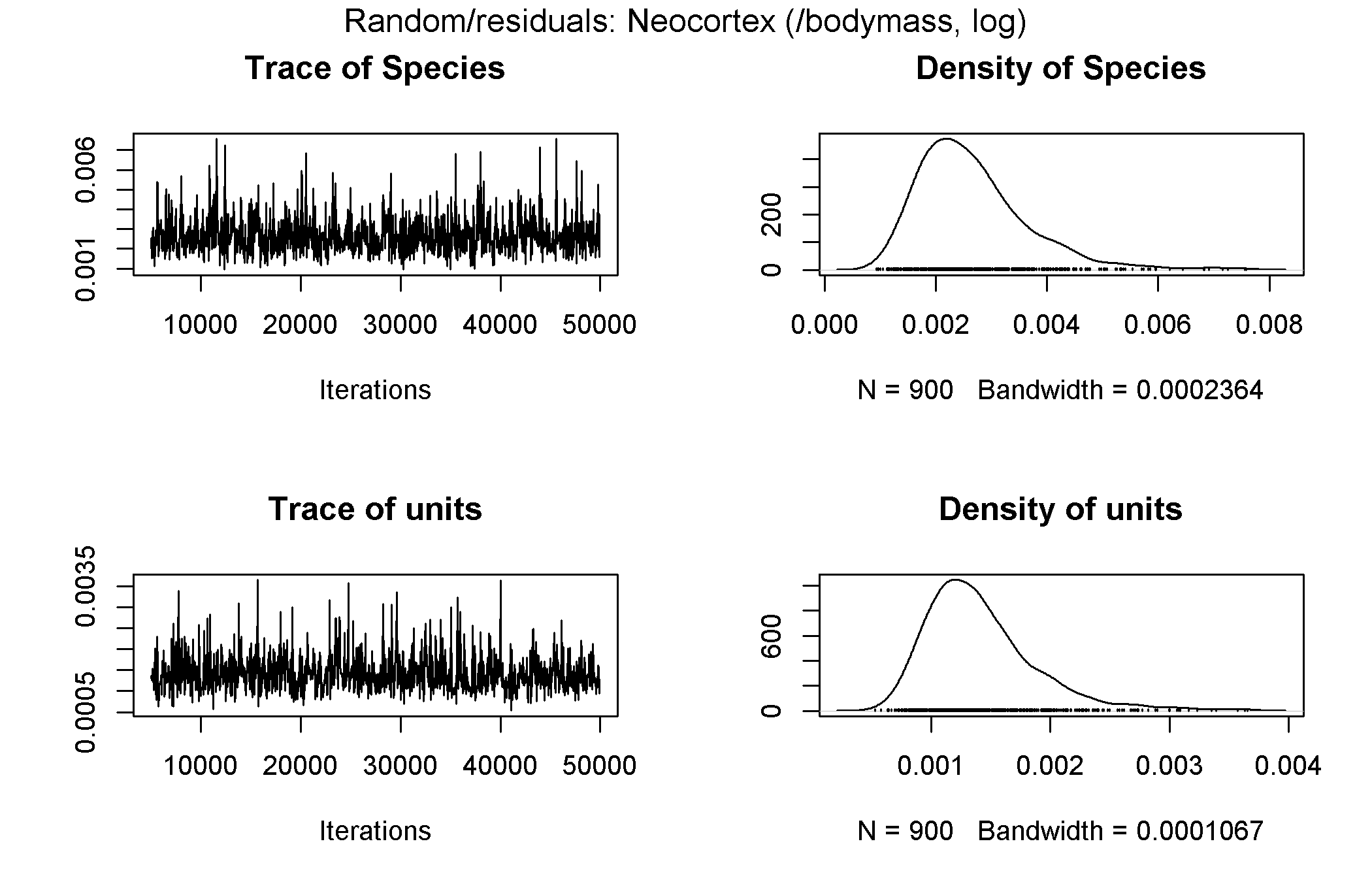

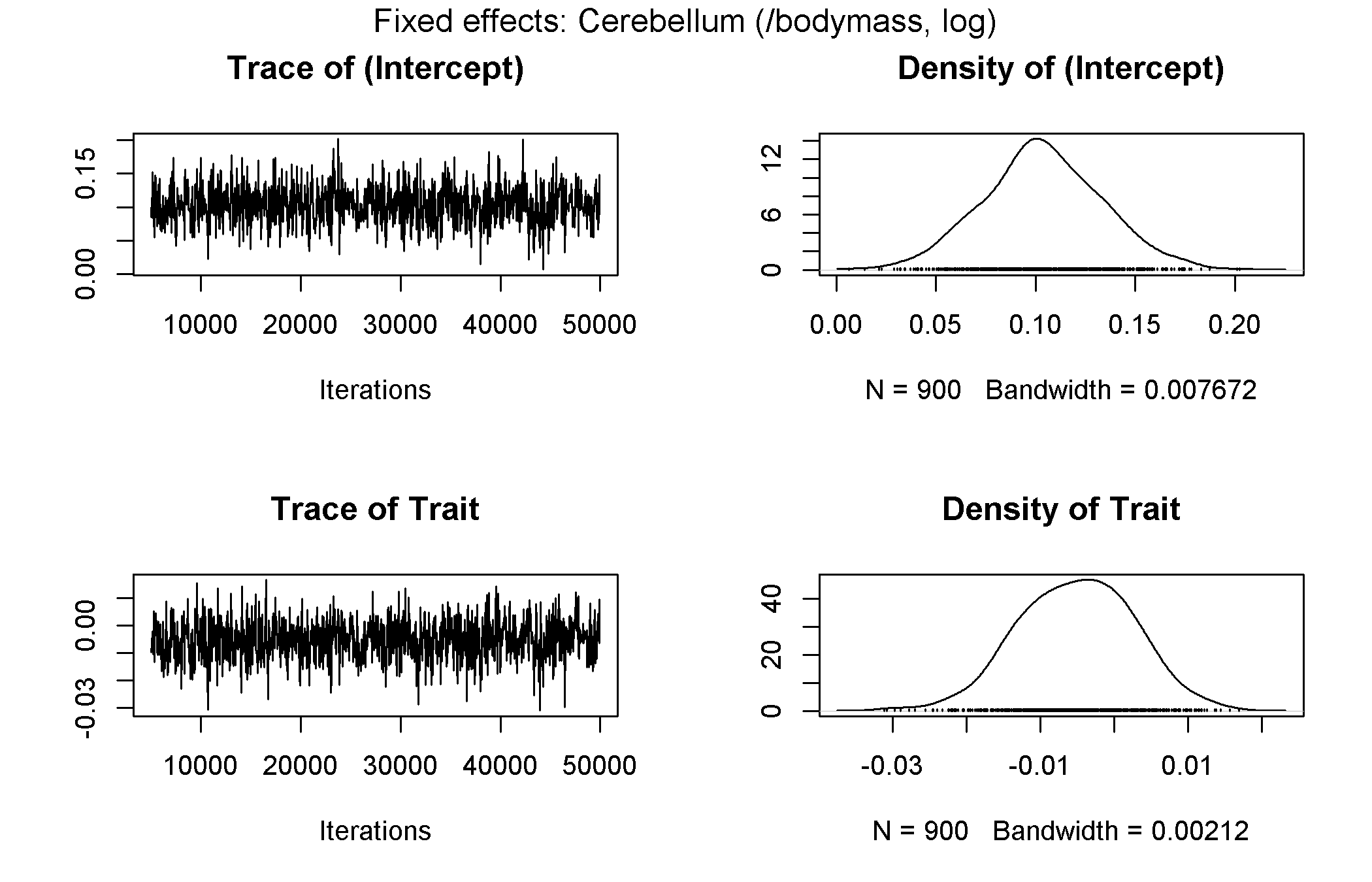

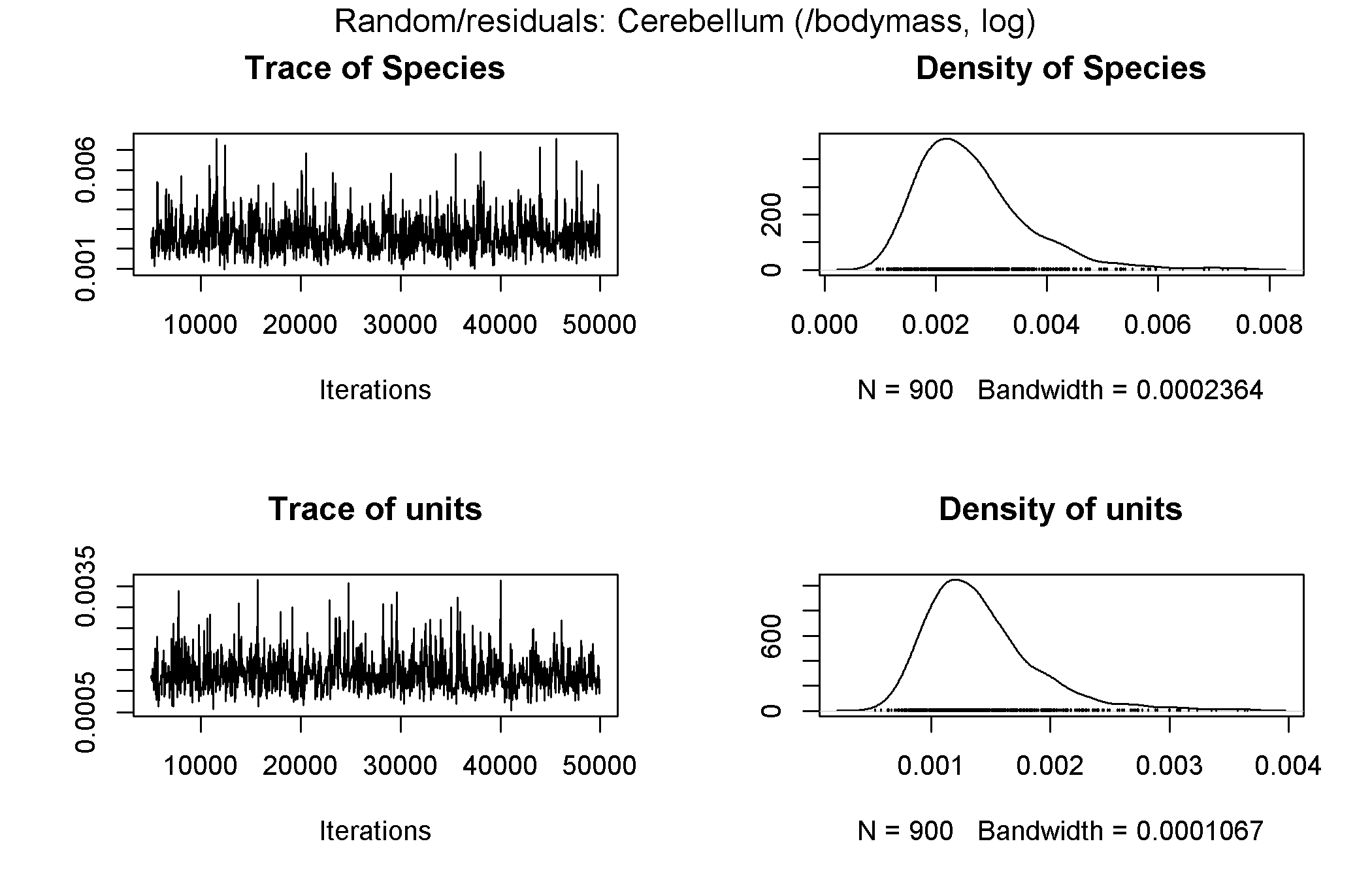

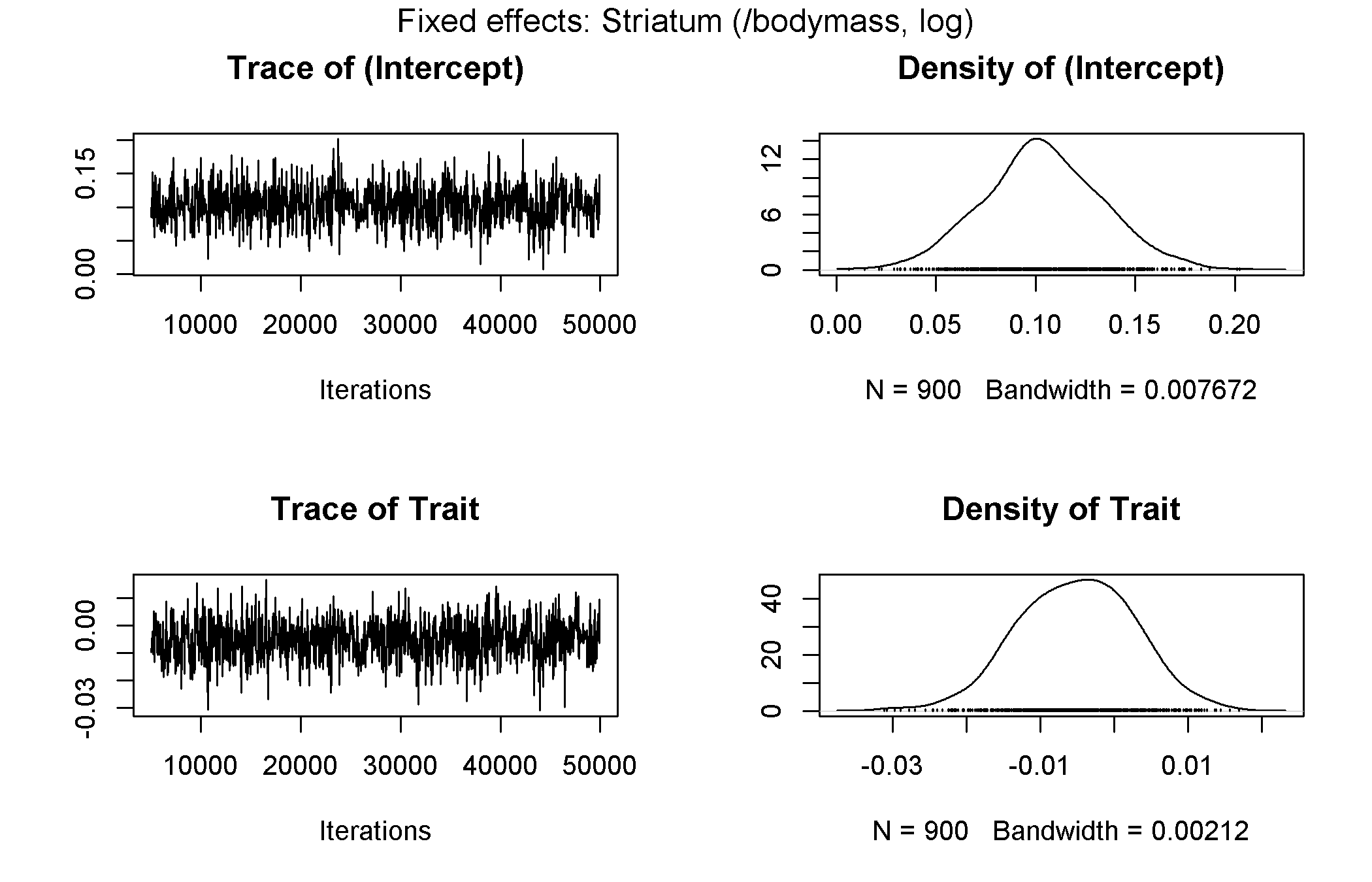

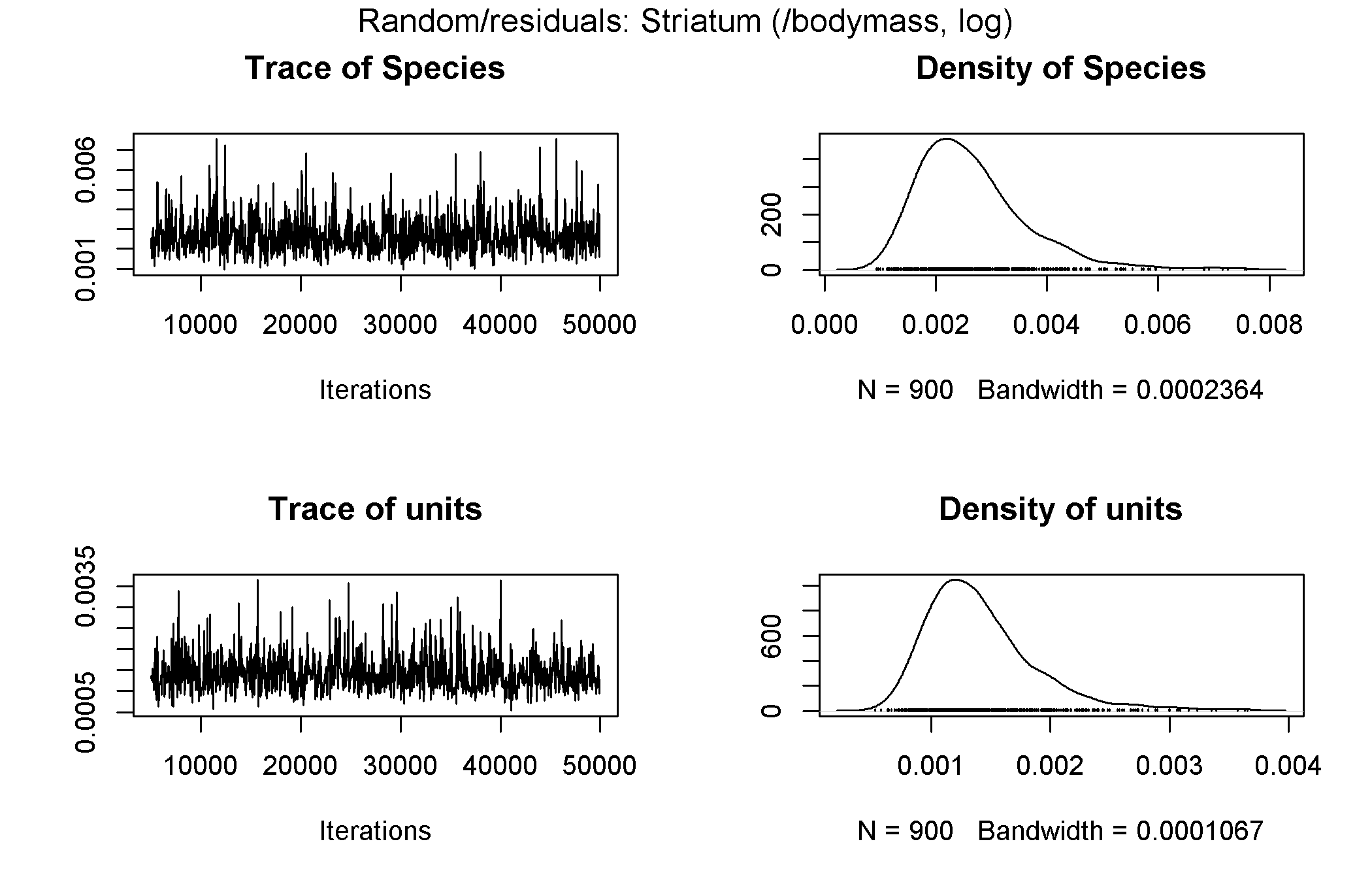

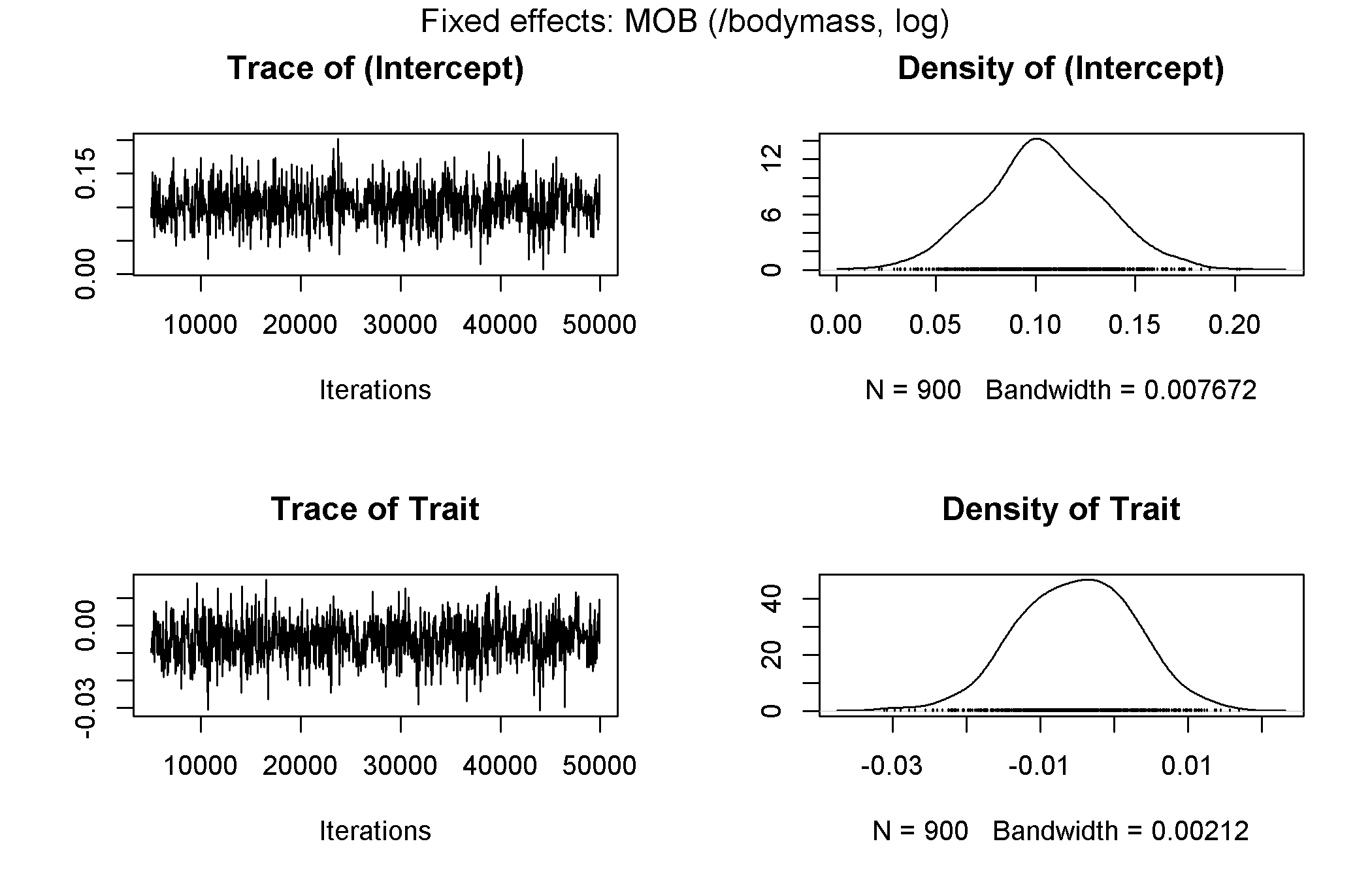

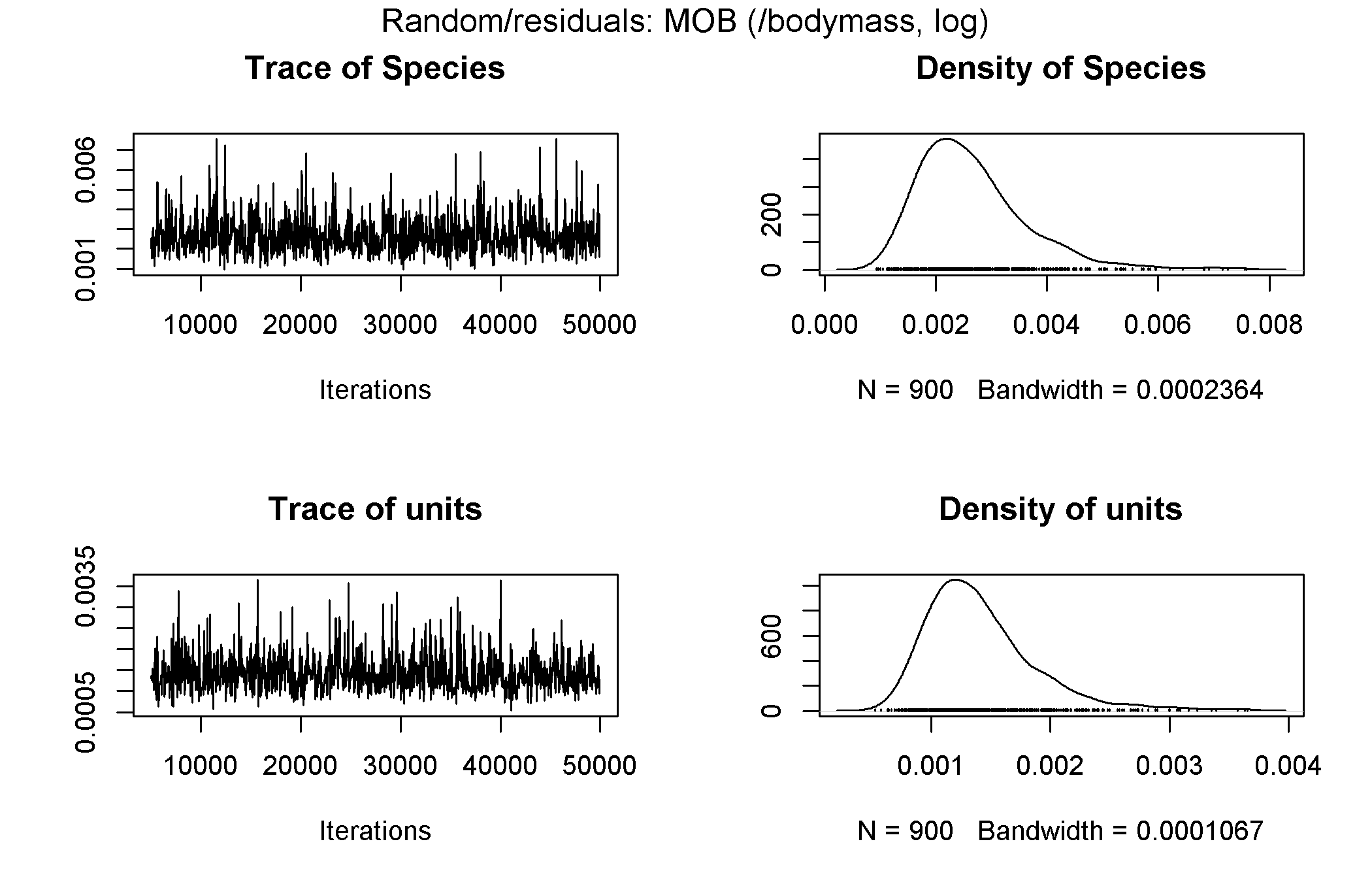

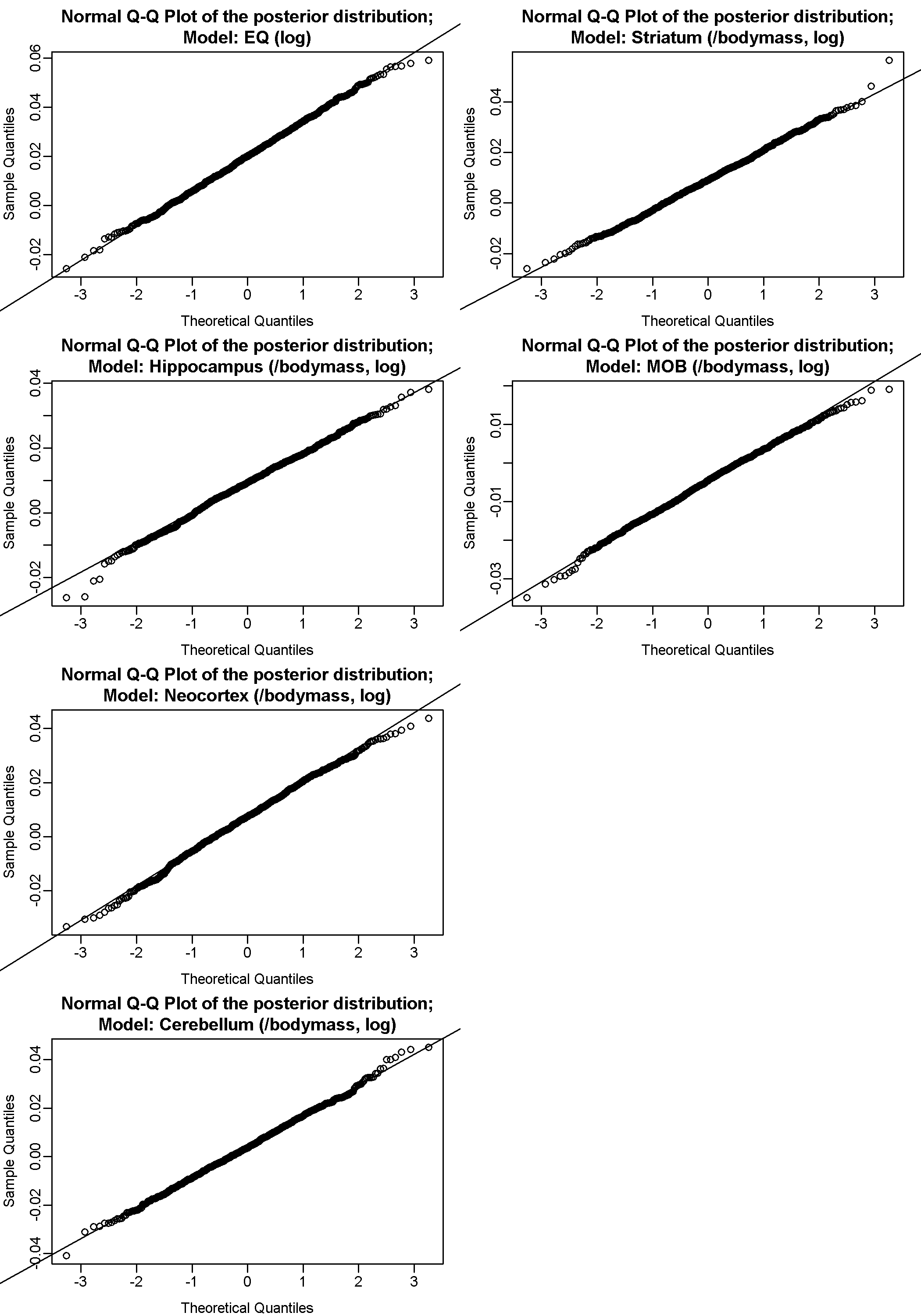

(c) Phylogenetic regressions: diversification and sympatry

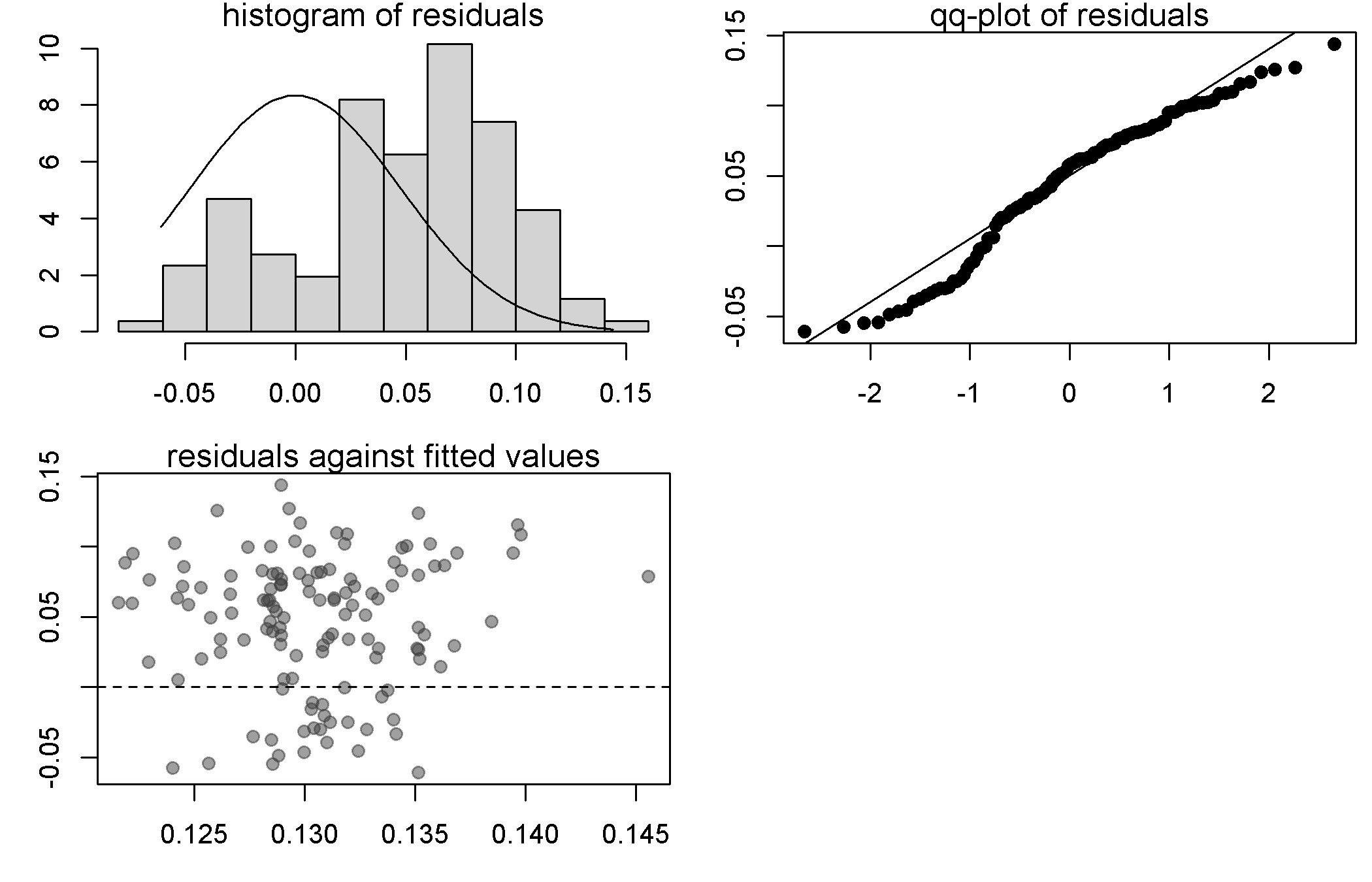

(d) Phylogenetic regressions: body mass and sympatry

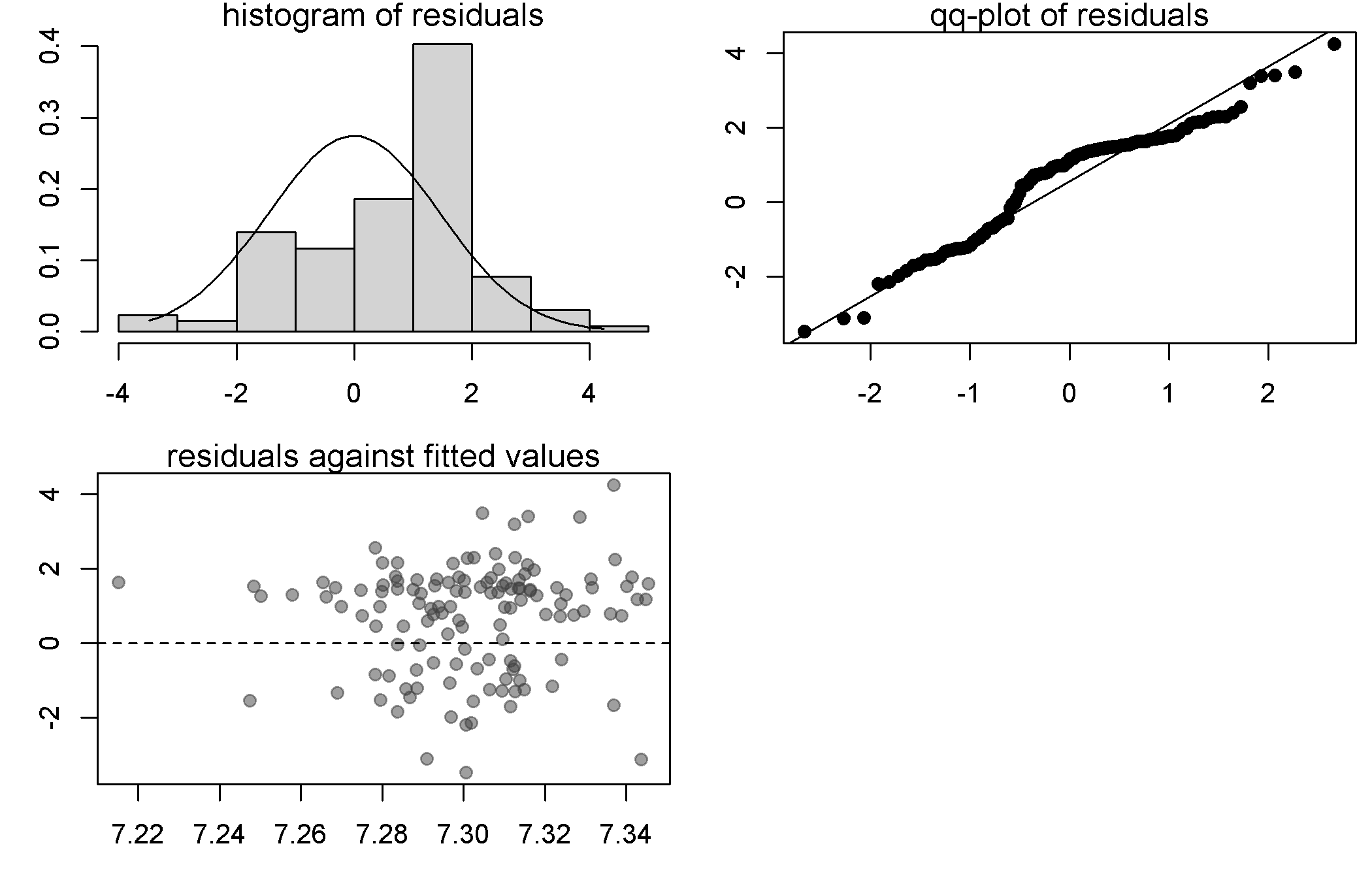

### R packages used

| **Package** | **Version** | **Reference** |
| --- | --- | --- |
| ape | 5.6-2 | Paradis E, Schliep K (2019). “ape 5.0: an environment for modern phylogenetics and evolutionary analyses in R.” Bioinformatics, 35, 526-528. |
| base | - | R Core Team (2020). R: A Language and Environment for Statistical Computing. R Foundation for Statistical Computing, Vienna, Austria. <https://www.R-project.org/>. |
| caper | 1.0.1 | Orme D, Freckleton R, Thomas G, Petzoldt T, Fritz S, Isaac N, Pearse W (2018). caper: Comparative Analyses of Phylogenetics and Evolution in R. R package version 1.0.1, <https://CRAN.R-project.org/package=caper>. |
| circlize | 0.4.15 | Gu Z, Gu L, Eils R, Schlesner M, Brors B (2014). “circlize implements and enhances circular visualization in R.” Bioinformatics, 30, 2811-2812. |
| cleangeo | 0.2-4 | Blondel E (2021). cleangeo: Cleaning Geometries from Spatial Objects. R package version 0.2-4, <https://CRAN.R-project.org/package=cleangeo>. |
| coda | 0.19-4 | Plummer M, Best N, Cowles K, Vines K (2006). “CODA: Convergence Diagnosis and Output Analysis for MCMC.” R News, 6(1), 7-11. <https://journal.r-project.org/archive/>. |
| datasets | - | R Core Team (2020). R: A Language and Environment for Statistical Computing. R Foundation for Statistical Computing, Vienna, Austria. <https://www.R-project.org/>. |
| doParallel | 1.0.17 | Corporation M, Weston S (2022). doParallel: Foreach Parallel Adaptor for the 'parallel' Package. R package version 1.0.17, <https://CRAN.R-project.org/package=doParallel>. |
| dplyr | 1.0.10 | Wickham H, François R, Henry L, Müller K (2022). dplyr: A Grammar of Data Manipulation. R package version 1.0.10, <https://CRAN.R-project.org/package=dplyr>. |
| ellipsis | 0.3.2 | Wickham H (2021). ellipsis: Tools for Working with .... R package version 0.3.2, <https://CRAN.R-project.org/package=ellipsis>. |
| foreach | 1.5.2 | Microsoft, Weston S (2022). foreach: Provides Foreach Looping Construct. R package version 1.5.2, <https://CRAN.R-project.org/package=foreach>. |
| geiger | 2.0.10 | Alfaro M, Santini F, Brock C, Alamillo H, Dornburg A, Rabosky D, Carnevale G, Harmon L (2009). “Nine exceptional radiations plus high turnover explain species diversity in jawed vertebrates.” Proceedings of the National Academy of Sciences of the United States of America, 106, 13410-13414.  Eastman J, Alfaro M, Joyce P, Hipp A, Harmon L (2011). “A novel comparative method for identifying shifts in the rate of character evolution on trees.” Evolution, 65, 3578-3589.  Slater G, Harmon L, Wegmann D, Joyce P, Revell L, Alfaro M (2012). “Fitting models of continuous trait evolution to incompletely sampled comparative data using approximate Bayesian computation.” Evolution, 66, 752-762.  Harmon L, Weir J, Brock C, Glor R, Challenger W (2008). “GEIGER: investigating evolutionary radiations.” Bioinformatics, 24, 129-131.  Pennell M, Eastman J, Slater G, Brown J, Uyeda J, Fitzjohn R, Alfaro M, Harmon L (2014). “geiger v2.0: an expanded suite of methods for fitting macroevolutionary models to phylogenetic trees.” Bioinformatics, 30, 2216-2218. |
| geosphere | 1.5-14 | Hijmans R (2021). geosphere: Spherical Trigonometry. R package version 1.5-14, <https://CRAN.R-project.org/package=geosphere>. |
| graphics | - | R Core Team (2020). R: A Language and Environment for Statistical Computing. R Foundation for Statistical Computing, Vienna, Austria. <https://www.R-project.org/>. |
| grDevices | - | R Core Team (2020). R: A Language and Environment for Statistical Computing. R Foundation for Statistical Computing, Vienna, Austria. <https://www.R-project.org/>. |
| iterators | 1.0.14 | Analytics R, Weston S (2022). iterators: Provides Iterator Construct. R package version 1.0.14, <https://CRAN.R-project.org/package=iterators>. |
| lattice | 0.20-45 | Sarkar D (2008). Lattice: Multivariate Data Visualization with R. Springer, New York. ISBN 978-0-387-75968-5, <http://lmdvr.r-forge.r-project.org>. |
| librarian | 1.8.1 | Quintans D (2021). librarian: Install, Update, Load Packages from CRAN, 'GitHub', and 'Bioconductor' in One Step. R package version 1.8.1, <https://CRAN.R-project.org/package=librarian>. |
| maps | 3.4.1 | Becker R, Minka T, Deckmyn. A (2022). maps: Draw Geographical Maps. R package version 3.4.1, <https://CRAN.R-project.org/package=maps>. |
| maptools | 1.1-4 | Bivand R, Lewin-Koh N (2022). maptools: Tools for Handling Spatial Objects. R package version 1.1-4, <https://CRAN.R-project.org/package=maptools>. |
| MASS | 7.3-58 | Venables WN, Ripley BD (2002). Modern Applied Statistics with S, Fourth edition. Springer, New York. ISBN 0-387-95457-0, <https://www.stats.ox.ac.uk/pub/MASS4/>. |
| Matrix | 1.5-1 | Bates D, Maechler M, Jagan M (2022). Matrix: Sparse and Dense Matrix Classes and Methods. R package version 1.5-1, <https://CRAN.R-project.org/package=Matrix>. |
| MCMCglmm | 2.34 | Hadfield JD (2010). “MCMC Methods for Multi-Response Generalized Linear Mixed Models: The MCMCglmm R Package.” Journal of Statistical Software, 33(2), 1-22. <https://www.jstatsoft.org/v33/i02/>. |
| methods | - | R Core Team (2020). R: A Language and Environment for Statistical Computing. R Foundation for Statistical Computing, Vienna, Austria. <https://www.R-project.org/>. |
| mvtnorm | 1.1-3 | Genz A, Bretz F, Miwa T, Mi X, Leisch F, Scheipl F, Hothorn T (2021). mvtnorm: Multivariate Normal and t Distributions. R package version 1.1-3, <https://CRAN.R-project.org/package=mvtnorm>.  Genz A, Bretz F (2009). Computation of Multivariate Normal and t Probabilities, series Lecture Notes in Statistics. Springer-Verlag, Heidelberg. ISBN 978-3-642-01688-2. |
| nlme | 3.1-159 | Pinheiro J, Bates D, R Core Team (2020). nlme: Linear and Nonlinear Mixed Effects Models. R package version 3.1-159, <https://CRAN.R-project.org/package=nlme>.  Pinheiro JC, Bates DM (2000). Mixed-Effects Models in S and S-PLUS. Springer, New York. doi:10.1007/b98882 <https://doi.org/10.1007/b98882>. |
| optimx | 2022-4.30 | Nash JC, Varadhan R (2011). “Unifying Optimization Algorithms to Aid Software System Users: optimx for R.” Journal of Statistical Software, 43(9), 1-14. doi:10.18637/jss.v043.i09 <https://doi.org/10.18637/jss.v043.i09>.  Nash JC (2014). “On Best Practice Optimization Methods in R.” Journal of Statistical Software, 60(2), 1-14. doi:10.18637/jss.v060.i02 <https://doi.org/10.18637/jss.v060.i02>. |
| parallel | - | R Core Team (2020). R: A Language and Environment for Statistical Computing. R Foundation for Statistical Computing, Vienna, Austria. <https://www.R-project.org/>. |
| permute | 0.9-7 | Simpson G (2022). permute: Functions for Generating Restricted Permutations of Data. R package version 0.9-7, <https://CRAN.R-project.org/package=permute>. |
| phylolm | 2.6.2 | Ho LST, Ane C (2014). “A linear-time algorithm for Gaussian and non-Gaussian trait evolution models.” Systematic Biology, 63, 397-408. |
| phytools | 1.2-0 | Revell LJ (2012). “phytools: An R package for phylogenetic comparative biology (and other things).” Methods in Ecology and Evolution, 3, 217-223. |
| picante | 1.8.2 | Kembel S, Cowan P, Helmus M, Cornwell W, Morlon H, Ackerly D, Blomberg S, Webb C (2010). “Picante: R tools for integrating phylogenies and ecology.” Bioinformatics, 26, 1463-1464. |
| plotrix | 3.8-2 | Lemon J (2006). “Plotrix: a package in the red light district of R.” R-News, 6(4), 8-12. |
| raster | 3.6-3 | Hijmans R (2022). raster: Geographic Data Analysis and Modeling. R package version 3.6-3, <https://CRAN.R-project.org/package=raster>. |
| RColorBrewer | 1.1-3 | Neuwirth E (2022). RColorBrewer: ColorBrewer Palettes. R package version 1.1-3, <https://CRAN.R-project.org/package=RColorBrewer>. |
| rgdal | 1.5-32 | Bivand R, Keitt T, Rowlingson B (2022). rgdal: Bindings for the 'Geospatial' Data Abstraction Library. R package version 1.5-32, <https://CRAN.R-project.org/package=rgdal>. |
| rgeos | 0.5-9 | Bivand R, Rundel C (2021). rgeos: Interface to Geometry Engine - Open Source ('GEOS'). R package version 0.5-9, <https://CRAN.R-project.org/package=rgeos>. |
| RPANDA | 2.2 | Morlon H, Lewitus E, Condamine F, Manceau M, Clavel J, Drury J (2016). “RPANDA: an R package for macroevolutionary analyses on phylogenetic trees.” Methods in Ecology and Evolution, 7, 589-597. R package version 1.4, <https://CRAN.R-project.org/package=RPANDA>. |
| rworldmap | 1.3-6 | South A (2011). “rworldmap: A New R package for Mapping Global Data.” The R Journal, 3(1), 35-43. ISSN 2073-4859, <http://journal.r-project.org/archive/2011-1/RJournal2011-1South.pdf>. |
| sf | 1.0-9 | Pebesma E (2018). “Simple Features for R: Standardized Support for Spatial Vector Data.” The R Journal, 10(1), 439-446. doi:10.32614/RJ-2018-009 <https://doi.org/10.32614/RJ-2018-009>, <https://doi.org/10.32614/RJ-2018-009>. |
| snow | 0.4-4 | Tierney L, Rossini AJ, Li N, Sevcikova H (2021). snow: Simple Network of Workstations. R package version 0.4-4, <https://CRAN.R-project.org/package=snow>. |
| sp | 1.5-1 | Pebesma EJ, Bivand RS (2005). “Classes and methods for spatial data in R.” R News, 5(2), 9-13. <https://CRAN.R-project.org/doc/Rnews/>.  Bivand RS, Pebesma E, Gomez-Rubio V (2013). Applied spatial data analysis with R, Second edition. Springer, NY. <https://asdar-book.org/>. |
| stats | - | R Core Team (2020). R: A Language and Environment for Statistical Computing. R Foundation for Statistical Computing, Vienna, Austria. <https://www.R-project.org/>. |
| stringr | 1.5.0 | Wickham H (2022). stringr: Simple, Consistent Wrappers for Common String Operations. R package version 1.5.0, <https://CRAN.R-project.org/package=stringr>. |
| svMisc | 1.2.3 | Grosjean P (2022). SciViews-R. UMONS, MONS, Belgium. <https://www.sciviews.org/SciViews-R/>. |
| tidyr | 1.2.1 | Wickham H, Girlich M (2022). tidyr: Tidy Messy Data. R package version 1.2.1, <https://CRAN.R-project.org/package=tidyr>. |
| utils | - | R Core Team (2020). R: A Language and Environment for Statistical Computing. R Foundation for Statistical Computing, Vienna, Austria. <https://www.R-project.org/>. |
| vegan | 2.6-4 | Oksanen J, Simpson G, Blanchet F, Kindt R, Legendre P, Minchin P, O'Hara R, Solymos P, Stevens M, Szoecs E, Wagner H, Barbour M, Bedward M, Bolker B, Borcard D, Carvalho G, Chirico M, De Caceres M, Durand S, Evangelista H, FitzJohn R, Friendly M, Furneaux B, Hannigan G, Hill M, Lahti L, McGlinn D, Ouellette M, Ribeiro Cunha E, Smith T, Stier A, Ter Braak C, Weedon J (2022). vegan: Community Ecology Package. R package version 2.6-4, <https://CRAN.R-project.org/package=vegan>. |
